# Microbiome-Derived Prion-Like Proteins and Their Potential to Trigger Cognitive Dysfunction

**DOI:** 10.1101/2023.10.19.563052

**Authors:** Jofre Seira Curto, Adan Dominguez Martinez, Paula Sotillo Sotillo, Martina Serrat Garcia, Monica Girona del Pozo, Maria Rosario Fernandez, Natalia Sanchez de Groot

## Abstract

Our life is intricately connected to microorganisms through infection or symbiotic relationships. While the inter-species propagation of prion-like proteins is well-established, their presence in the microbiome and impact on the host remains largely unexplored. To address this, we conducted a systematic study integrating *in silico*, *in vitro,* and *in vivo* analyses, showing that 63% of the gastrointestinal tract microbiome encodes prion-like sequences. These sequences can form amyloid fibrils capable of interfering with the aggregation of the Amyloid-beta-peptide and promoting the aggregation and propagation of the Sup35 prion. Finally, when *C. elegans* were fed with bacteria expressing chimeras of our prion candidates, it resulted in the loss of sensory memory, reproducing the Alzheimer’s model phenotype. In our model, memory impairment is linked to aggregate fragmentation and its susceptibility to degradation. Taken together, these findings show that the gut microbiota serves as a potential reservoir of prion-like sequences, supporting the idea that microbial products may influence the pathogenesis of neurodegenerative diseases.

## Introduction

One hallmark shared by many neurodegenerative diseases is the presence of insoluble aggregates of misfolded proteins. These aggregates frequently consist of amyloid characterized by a distinct cross-beta structure. Some of these amyloid-forming proteins exhibit prion-like properties, enabling transmission between cells. This phenomenon is notably observed in Alzheimer’s (AD) and Parkinson’s disease (PD), where amyloid-associated proteins (amyloid-β-peptide (Aβ) and tau for AD, and alpha-synuclein for PD) can propagate within the neuronal system ^1^, and, more importantly, this phenomenon can also occur between organisms of different species ^2^.

Traditionally, researchers have primarily focused on genetic and environmental factors as triggers for neurodegenerative diseases. However, in recent decades, a new area of study has emerged, suggesting that the gut microbiota, an ecosystem of trillions of microorganisms residing in the intestinal tract in symbiosis with the host, may play a pivotal role in the onset and progression of these diseases ^3^.

The brain-gut microbiota axis is believed to act as a bidirectional link between the gastrointestinal tract and the central nervous system through three key mechanisms: endocrine signaling, the immune system, and direct neuronal communication^4^. The vagus nerve, responsible for transmitting signals in both directions, mediates the neuronal connection between the gut and the brain ^5^. Previous studies proposed that amyloid proteins and other metabolites from the gastrointestinal tract can be transported along the vagus nerve to the brain^6^, thereby supporting their potential impact on the development and progression of neurodegenerative diseases^3,7^. More recently, it has been reported that also gut bacteria can travel through the vagus nerve to the brain, when dysbiosis and intestinal permeability are induced, but also as a consequence of a neurological disease^8^.

In this context, diverse studies conducted over the last decade have linked alterations in the composition and function of the gut microbiota with neurodegenerative diseases like AD and PD. Gut dysbiosis can be triggered by factors such as diet, antibiotic use, and stress, and these changes can have significant consequences on host health, contributing to various disorders, from gastrointestinal issues to metabolic disorders ^3,9^. However, it’s only recently that the gut microbiome has gained recognition as a potential biomarker and disease modifier concerning neurodegenerative diseases.

This connection is supported by causal evidence in animal model studies, where antibiotic treatments and microbiota transplantation can modulate the pathology^10^. Hence, it has been proposed that the microbiota amyloid protein can cross-seed host proteins aggregation. In agreement, the expression of the amyloid protein curli in *Escherichia coli* accelerates alpha-synuclein aggregation in a PD mouse model^11^.

To evaluate the existence of other bacterial proteins capable of interacting with and influencing the aggregation of host proteins, we conducted a computational analysis that revealed that 63% of the species within the gut microbiome encode prion-like sequences. Our bench investigation confirmed their ability to form and propagate amyloid aggregates through their amyloid-forming cores, and to cause sensory memory impairment in *C. elegans*.

All in all, our results provide insights into the intricate connection between microbiota and neurodegenerative diseases, contributing to the knowledge required for future prevention and treatment of these detrimental conditions.

## Results and discussion

### Screening of prion-like sequences in the gut microbiome

Previous publications have reported an average of 0.3% of prion-like sequences per genome in the bacterial domain^12–19^. In this context, it would be interesting to investigate their prevalence within the gut microbiome. With this aim, we screened the genes collected in the NIH Human Microbiome Project (NIH-HMP)^20,21^ using three different algorithms: PAPA^17^ (prion aggregation prediction algorithm), PLAAC (Prion-Like Amino Acid Composition)^16^ and pWALTZ^18^ (Figure 1A; Methods). These approaches identify disordered sequences with prion-like composition and provide a score that reflects the probability of a sequence behaving as a prion.

**Figure 1.**
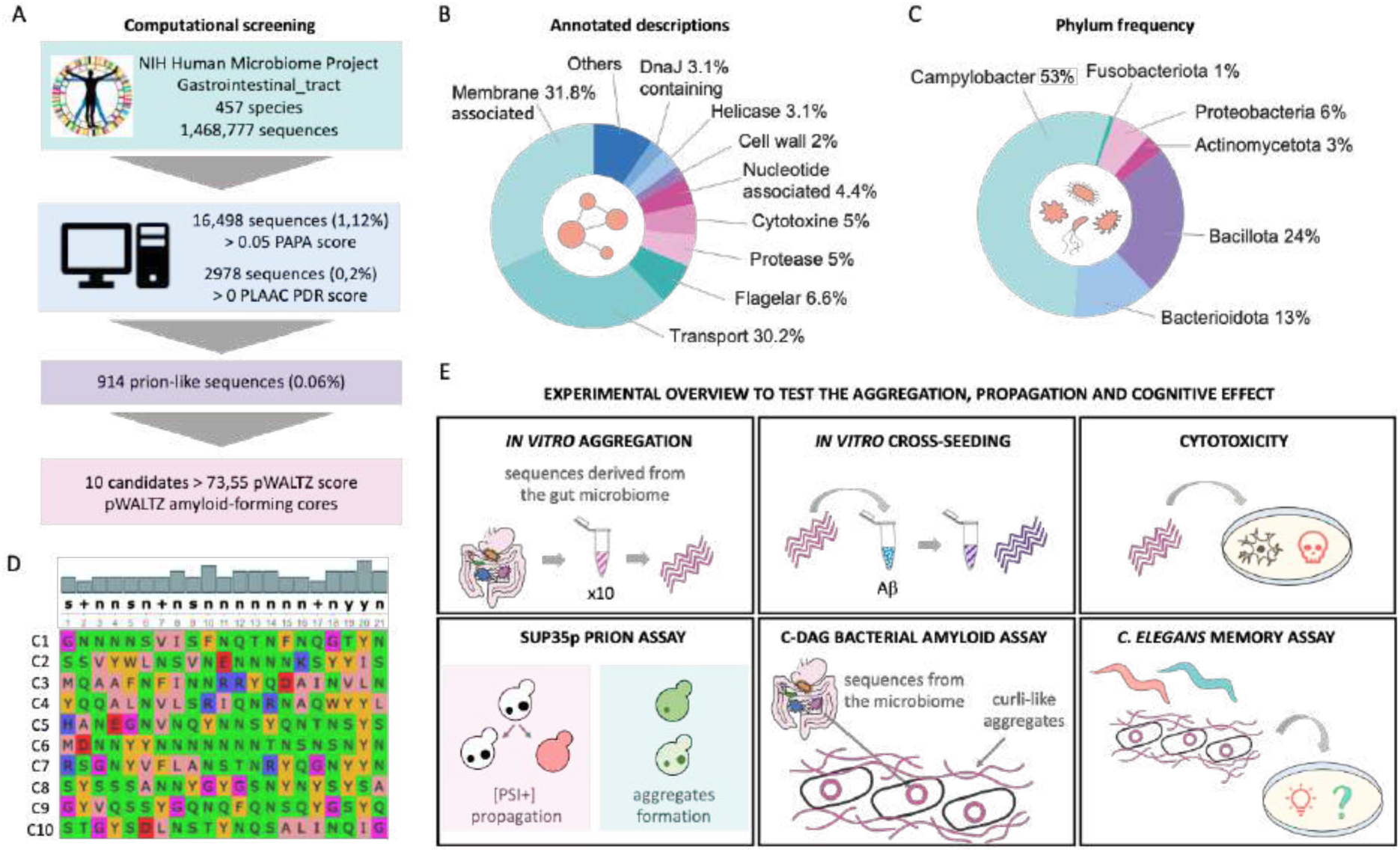
Screening of Prion-Like Sequences in the Gut Microbiome. A) Flowchart illustrating the procedure for screening prion-like sequences in the gut microbiome and selecting amyloid-forming core candidates. B) Pie chart depicting the distribution of annotated descriptions with the 914 prion-like sequences identified in the gut microbiome. It just shows those annotations found with at least 2% of frequency. C) Pie chart illustrating the distribution of phyla associated with the 914 prion-like sequences detected in the gut microbiome. It just shows those phyla found with at least 1% of frequency. D) Sequence alignment displaying the 10 peptide sequences studied in this work and the consensus sequence. E) Outline of the experimental methods used to analyze the potential influence of selected prion-like candidates from the gut microbiota on the development of neurodegenerative diseases: i) amyloid forming ability, ii) interference with a host protein, iii) cytotoxic potential, iv) functional propagation in vivo, v) aggregation in a bacterial, vi) cognitive effects upon ingestion.

The original list, derived from the NIH-HMP^20,21^, contains 457 species and 2,540,626 sequences, from which 1,468,777 are unique entries (Figure 1A, Supplementary Table 1). Using PAPA algorithm, we obtained 16,498 sequences (1.12%) positive in prion aggregation propensity, while using a more stringent score of PLAAC we obtained 2,978 sequences (0.20%) positive in prion-like domain (Methods). Merging these two sets, we identified 914 sequences (0.06%) meeting both criteria, which we refer to as the positive prion-like set (Supplementary Data 1). The proportion of prion-like sequences obtained with our approach (0.06%) aligns with previous reports of prion-like sequences in the bacterial domain^12–19^. These 914 sequences have been found in 284 species, showing that 63% of the species within the gut data set encode for prion-like sequences. Given the ability of prion-like proteins to spread across species, this finding sheds light on the role of gut microbiota as a reservoir of these proteins with infectious potential.

### Prion-like proteins in the gut microbiome

One striking feature of the 914 positive prion-like sequences is the high proportion of hypothetical proteins (uncharacterized)^20,21^, which constitute 40% of the set (Supplementary Table 1). Most of the sequences collected in the NIH-HMP have been identified via genome shotgun sequencing, and those identified as hypothetical are open reading frames without a characterized homolog in the protein databases. With a simple examination of their disorder regions predicted by MobiDB-lite (UniProt)^20,21^ it is possible to detect repeated patterns (Supplementary Figure 1 and Supplementary Table 1), which suggests that these proteins may have conserved, similar, yet unidentified, cellular roles that require a prion-like composition. This highlights the importance of further investigating the molecules produced by the human microbiota^22–24^.

The remaining 60% of positive prion-like sequences, with estimated homology, can be grouped into 10 categories according to their annotated descriptions (Figure 1B, Supplementary Table 1). The two main categories, membrane associated (31.8%) and transporter proteins (30.2%) (Figure 1B), play direct roles in facilitating interactions between bacterial cells and the extracellular milieu. These categories include functions such as membrane receptors, transporters, and adhesins, which require the formation of contacts where the sticky properties of the prion-like domains could be important^22–24^. For example, some transporter proteins need to interact with other proteins to form the scaffolding structures that enable the transport of molecules across the membrane (10.3389/fmicb.2018.01737). Similarly, other membrane proteins contribute to external environment sensing or cellular attachment to surfaces, such as adhesion proteins. The sequences associated with flagella may be involved in bacterial movement and migration, which are also functions related to interactions with the external space^25,26^. Other abundant functions are associated with the protein quality control machinery (DnaJ and proteases) and nucleotide binding (such as helicase or single-strand binding), which have been widely associated with prion-like proteins^12–19^.

Membrane associated proteins may also be involved in aggregation and transmissibility events. In fact, the interaction with membranes is an important source of conformational alteration and aggregation^27^. Additionally, considering that the amyloid precursor protein (APP) itself is an integral membrane protein^28–30^, it raises the intriguing possibility of yet undiscovered bacterial proteins sharing a similar fate. Many bacterial membrane proteins are involved in interacting with host cells and the environment, which can be a possible way of triggering or interfering in the cascade of events that can contribute to the emergence of diseases. For example, curli amyloid fibers formed by certain bacteria, including *E. coli*, have a dual role as both biofilm components and potential virulence factors^31–35^. The adhesive properties of curli enhance the bacteria’s ability to colonize diverse environments. Importantly, there is substantial evidence indicating that biofilm-forming proteins can influence the aggregation of neuronal proteins, thereby impacting the development of neurodegeneration^31,32,36,37^.

### Gut microbial species encoding for prion-like sequences

The analysis of the 914 positive prion-like sequences revealed that most of them belong to phyla previously related to neurodegenerative pathologies, particularly Alzheimer’s disease^38^. More than half of them were found in the phylum Campylobacter (53%) (Figure 1C). In our list, this phylum is mostly composed of *Helicobacter pylori* species, known for causing peptic ulcers and infecting approximately 50% of the global population^39^. Remarkably, many Alzheimer’s patients have experienced active and/or latent *Helicobacter* infections, suggesting a potential role for this pathogen in the development of the pathology^40,41^.

The second most abundant phylum, in terms of prion-like sequences, was Bacillota, accounting for 24% of the identified sequences. This phylum includes *Coprobacillus* (8%), *Eubacterium* (9%), *Ruminococcus* (9%), *Lactobacillus* (25%) and *Clostridium* (17%). Consistently, in a previous analysis of the whole Bacteria Domain^13^, we observed a high abundance of prion-like sequences in bacteria belonging to the *Clostridium* genus. Moreover, the Rho protein from *Clostridium botulinum* was the first prion-like protein experimentally confirmed in bacteria^42^.

Another significant finding was that 13% of the prion-like sequences corresponded to the phylum Bacteroidetes, which is one of the most prevalent phyla in the human gut microbiome^43^. An increase in the proportions of *Bacteroides* and *Prevotella* (genus included in this phylum) has also been observed in individuals with Alzheimer’s disease^38,44–46^. Therefore, the correlation between the increased proportion of these bacteria and their high number of prion-like sequences may indicate their potential role in the development of the pathology.

In contrast, we did not detect prion-like sequences in Firmicutes, the other prevalent phylum in the human gut microbiome^43^. This observation is particularly noteworthy, as Firmicutes are associated with a healthy microbiota that becomes altered and decreases in individuals with Alzheimer’s disease^38,44^.

Furthermore, Proteobacteria, which includes primarily non-pathogenic strains as well as the widely studied *E. coli*, was found to be associated with a significant proportion of putative prion-like proteins. A recent study indicates that *E. coli* is present in more than 90% of the population tested^47^. *Escherichia* also exhibits increased proportions in Alzheimer’s patients^47^, and our predictors have identified several prion-like sequences associated with this genus.

### Exploring prion-like candidates within the gut microbiota

To validate our finding, we selected ten candidates from the set of 914 positive prion-like proteins and assessed their potential to trigger neurodegenerative disease *in vitro* and *in vivo* (Figure 1D-E). We first arranged all the sequences by their pWALTZ score to prioritize the candidates with higher values, and, to guarantee a robust predisposition for aggregation, we enforced a minimum score of 73.5^18^. To facilitate the experimental procedure, we excluded sequences labeled as hypothetical, those containing cysteine residues, and those with multiple transmembrane domains. Following the conventional prion composition, we selected sequences with high content of asparagine and glutamine, above 20% (Supplementary Data 1). In agreement with the prion prediction algorithms (Supplementary Table 1)^16–18^, AlphaFold^48^ analysis consistently identified the presence of disordered regions in the sequence of all ten selected candidates (Supplementary Figures 2-11).

We also specifically selected sequences from bacterial strains associated with the genera previously reported to be altered in Alzheimer’s disease patients^40,41,38^. Utilizing the PSORTb tool^47^, we primarily chose candidates with a high likelihood of an extracellular location, enhancing their potential for interaction with the host environment. Overall, we aimed to construct a diverse collection of proteins from both Gram groups, from different genera, and with annotated descriptions (Supplementary Data 1, Supplementary Table 2).

### *In vitro* aggregation of the amyloid-forming cores

To assess the aggregation properties of the selected prion-like candidates, our investigation focused on their amyloid-forming cores, regions capable of self-aggregation but also of driving the aggregation of the entire protein^18,42^. We identified these sequences by combining PAPA and pWALTZ algorithms (Methods). In this way, we identified the 21-amino acid segment with the highest aggregation propensity within the main prion-like domain^13,18,19^. Subsequently, these 21 amino-acid segments were chemically synthesized and their ability to self-assemble into fibrillar aggregates was tested.

The synthetic peptides were diluted in the aggregation buffer and then incubated under non-agitation conditions at 37 °C (Methods). The presence of amyloid aggregates was assessed using Transmission Electron Microscopy (TEM), Fourier-transform infrared spectroscopy (FT-IR), and binding to the amyloid dyes Thioflavin-T and Congo red. All peptides exhibited the ability to form fibrillar structures (Figure 2A), rich in beta-sheet conformation (Figure 2B). Additionally, all peptides exhibited positive binding for at least one of the amyloid-specific dyes (Figure 2C-D and Supplementary Figures 2-11). Taken together, these findings demonstrate the capacity of the predicted amyloid-forming cores from the gut microbiome, to catalyze self-nucleation into amyloid fibrils.

**Figure 2.**
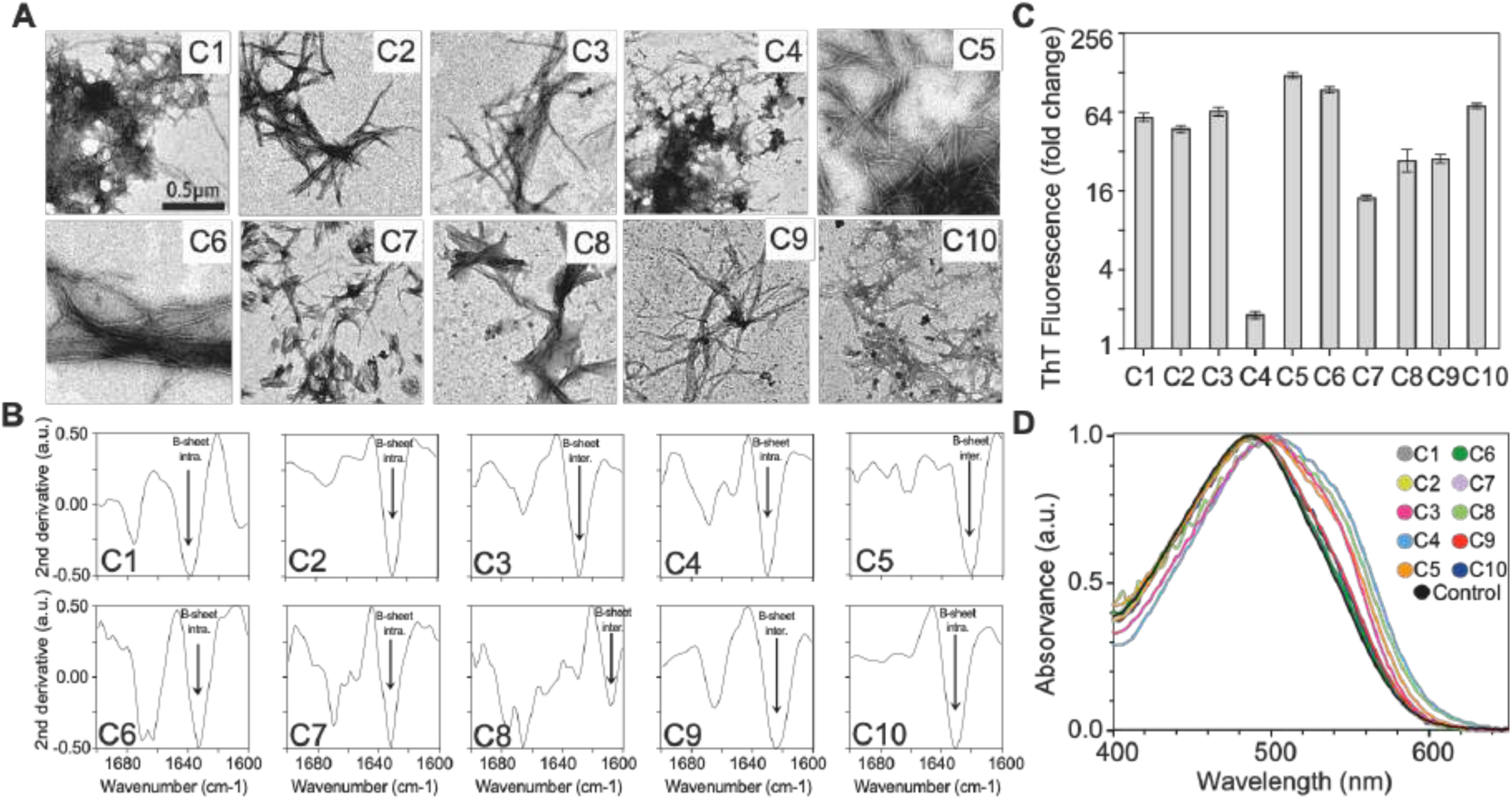
Amyloid Aggregation of Sequences from the Gut Microbiome. A) TEM images showing fibrillar aggregates formed by the selected amyloid-forming cores. B) FTIR spectrum of the aggregated peptides. C) Thioflavin-T fluorescence increase (fold change) upon binding the aggregated peptides (N=3). D) Congo Red absorbance in the presence of the aggregate’s peptides (see Supplementary Figures 2-11 for individual scans).

### Interference with a host protein aggregation

As noted before, the peptides selected are derived from the genomes of bacteria found altered in Alzheimer’s disease patients^38^. In this line, we also wanted to study their ability to interfere with the aggregation process of the amyloid-β-peptide (Aβ). Pre-aggregated peptides were incubated with soluble Aβ40 and the aggregation kinetics were monitored following the Th-T fluorescence intensity (Figure 3A, Supplementary Figure 12). The half-life (Figure 3B) and lag times (Figure 3C) exhibited a significant influence in 9 out of 10 peptides tested. Most peptides accelerated the aggregation kinetics, except for C10, which slowed it down (Figure 3A-C and Supplementary Figure 12). Overall, this result supports the existence of prion-like sequences in the gut microbiome with the potential to interfere with host proteins and contribute to pathogenesis^11^.

**Figure 3.**
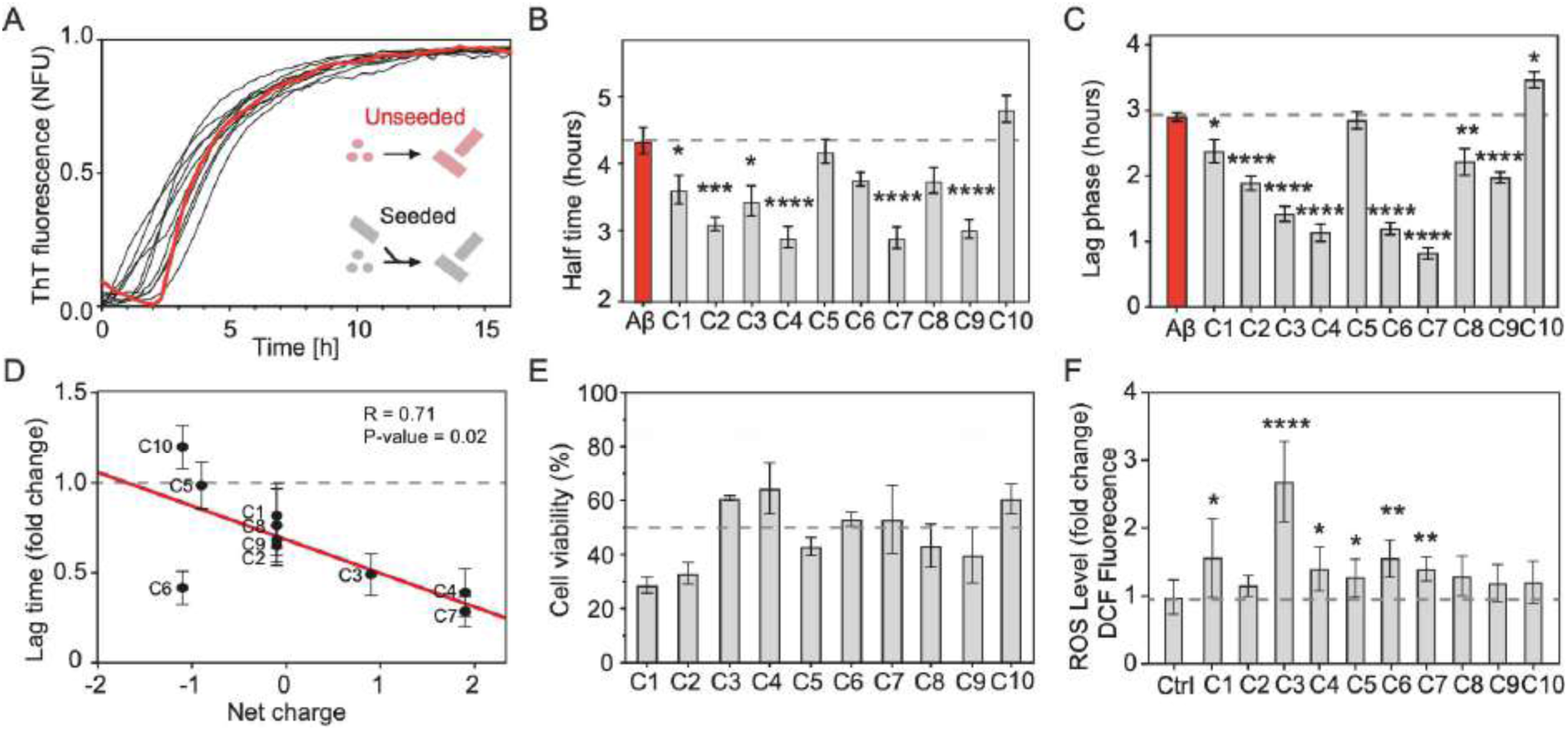
Seeding and Toxicity Potential of Sequences from the Gut Microbiome. A) Aggregation kinetics of Aβ40 peptide seeded by amyloid-forming cores from the gut microbiome (for separated kinetics, see Supplementary Figure 12). All kinetic assays were performed four times in triplicates. B) Half-times of seeded aggregation kinetics (one-way ANOVA, SEM). C) Lag phase of seeded aggregation kinetics (one-way ANOVA, SEM). D) Correlation between the lag time and the net charge of the amyloid-forming cores used for seeding. Without C6, R increases to 0.95. E) The viability of neuron-differentiated SH-SYS5 cells was assessed by the MTT assay, incubating them with 10 µM of peptide aggregates (N=3; for lower concentrations see Supplementary Figure 16). Error bars are SEM. F) DCFDA/H2DCFDA Cellular ROS assay. Fluorescence was observed after 4 hours of exposure to the different peptides. The bars show the fluorescence fold change in comparison to the negative control (Ctrl). The significance against the control assay was calculated using an unpaired t-test (N=3).

Looking for a relationship between the peptide’s composition and its effect on Aβ40 aggregation, we observed a robust correlation between the peptide net charge and lag-time kinetics, except for C6 (Figure 3D). A more positive net charge results in a faster Aβ40 aggregation, whereas a more negative one results in a slower one. Similar findings have been reported previously in studies investigating inhibitory peptides against Aβ^49–51^, which bears a negative charge (-2.9) at pH 7.4^52,53^.

To show that not just the amyloid-forming core, but also the full protein can form fibrillar aggregates and influence Aβ40 aggregation kinetics, the parental protein that contains the C4 peptide (C9L6N5, a protein with a DNAJ domain) was also tested. This protein, composed of 212 amino acids, was expressed in *E. coli* and subsequently purified (Supplementary Figure 15, Methods). We observed that the purified protein was able to aggregate into fibrillar structures (Supplementary Figure 13). Subsequent incubation of Aβ40 with pre-aggregated C9L6N5 accelerated the aggregation kinetics, in a similar way as its amyloid-forming core (Supplementary Figure 13). This result supports that not only the amyloid-forming cores but also the proteins from which these sequences originate, can form fibrillar aggregates and can seed host molecules.

### Toxicity and oxidative stress on neuron-differentiated cells

We also assessed whether peptides derived from the gut microbiome could induce phenotypic hallmarks associated with neurodegenerative diseases in neuronal differentiated cells (SH-SY5Y) (Methods). After proving their stability in cell culture conditions (Supplementary Figure 14-15), we evaluated their potential to induce cell death using the MTT assay and oxidative stress using the DCFDA/H2DCFDA assay (Figure 3E-F, Supplementary Figures 16).

The analysis of the MTT assays showed that peptides with a net charge close to 0 exhibit greater toxicity compared to those with a higher charge (Supplementary Figure 17). This suggests a potential role for electrostatic interactions in protecting against cytotoxicity. Polar, uncharged polypeptides tend to have an amphipathic nature that facilitates their interaction with membranes, which, as in the case of antimicrobial peptides, may result in cell toxicity. Taking all the results into account, the presence of these peptides causes a general drop in cell viability below 70% (Figure 3F), indicating that the selected amyloid-forming cores from the gut microbiome hold the potential to induce cellular damage upon aggregation.

The oxidative stress test revealed an increase in H2DCFDA fluorescence across all cases (Figure 3F), with significance in six out of ten peptides. This data suggests that the peptides’ cytotoxicity (Figure 3E) can, in part, be attributed to oxidative stress. However, we cannot rule out that other mechanisms, such as membrane disruption, autophagy dysfunction, or proteasomal dysfunction, may also be contributing to^27,56^.

Among all the peptides tested, the aggregates of C1 and C2 exhibited especially high toxicity (Figure 3E and Supplementary Figure 16), reducing cell viability to nearly 30% at 10 uM. Interestingly, C1 and C2 are fragments of two vacuolating cytotoxins (VacA) implicated in pathogenesis and gastric cancer^57,58^. Both are located in P33 domain, a segment required to form the oligomeric conformation able to bind and internalize into mammalian cells^57,58^. Notably, C1 and C2, despite their short length, retain a significant portion of VacA toxicity. This result also suggests that the prion-like composition of C1 and C2 may contribute to the oligomerization and toxic potential of their parental proteins.

### Amyloid-forming cores from the gut microbiome are functional in yeast

To evaluate the ability of the ten amyloid-forming cores^18,42^ to lead to protein aggregation and propagation *in vivo* (Figure 1E), we opted to use a common and well-known framework, the yeast prion protein Sup35 (Figure 4A) ^53,59,60^. We removed its first forty amino acids, the nucleation domain ^61,62^, and replaced them with our collection of amyloid-forming cores (Supplementary Data 1). The sequential analysis of the resultant chimeras indicates similar aggregation and prion-like properties as the original Sup35 (Supplementary Data 2).

**Figure 4.**
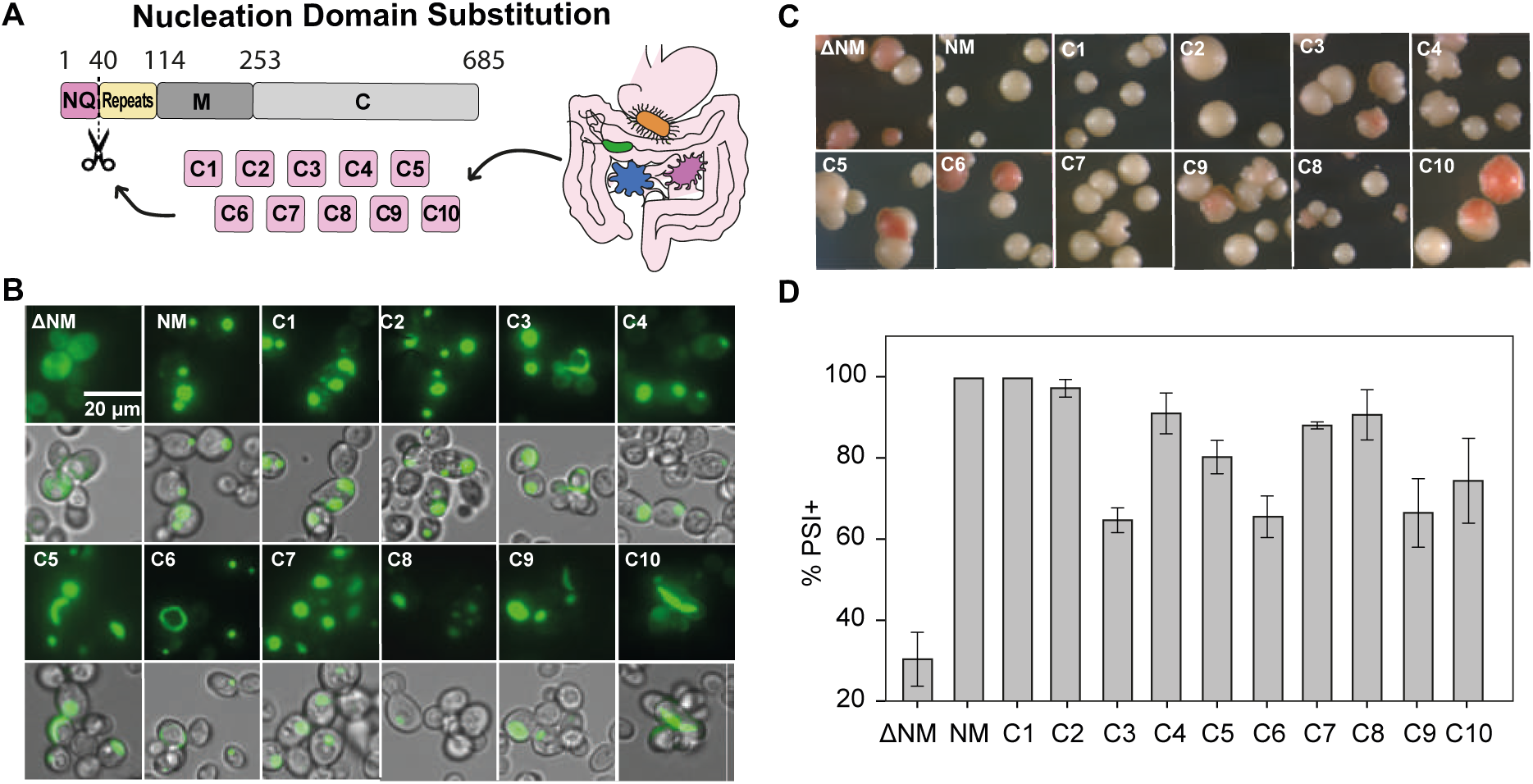
Aggregation and Propagation Analysis in Yeast. A) Diagram illustrating the substitution of the Sup35 nucleation region (NQ) with 10 amyloid-forming cores from the gut microbiome. B) Fluorescent images showing intracellular aggregates. C) Representative images of white/red colonies from the non-sense suppression assay. D) Percentage of [PSI+] for each yeast strain analyzed. All assays were performed at least in triplicates. All are significantly more [PSI+] concerning ΔNM (≤0.05, one-way ANOVA). Error bars are SEM. For more details about the yeast phenotype see Supplementary Data 4.

We used a nonsense suppression assay to evaluate the formation and propagation of the Sup35 aggregates ([*PSI*^+^])^61^. In this system, the *ADE1* gene (*ade*1-14), causes the accumulation of a red pigment (an adenine precursor) that colors colonies in red when Sup35 is soluble (non-prion colonies[*psi*^-^]), and white or pink when it becomes insoluble (prion colonies [*PSI*^+^]). Since the conversion to [*PSI*^+^] is a rare event, we induced it by over-expressing the NM segment of Sup35 fused to GPF (Supplementary Data 3); this increases the rate of prion conversion and facilitates monitoring the aggregation inside the cell. As positive and negative controls, we employed cells expressing unmodified Sup35 (Sup35NM) and Sup35 lacking the first forty amino acids (ΔSup35), respectively. Microscopy analysis has revealed that Sup35NM concentrates the fluorescence in bright foci, while ΔSup35 exhibits a homogeneous distribution throughout the cytosol (Figure 4B, Supplementary Figure 18). In addition, all the cells expressing a Sup35 variant, carrying the predicted amyloid-forming cores from the gut microbiome, exhibit fluorescent foci.

Consistently, the [*PSI*^+^] assay displays 100% white colonies in Sup35NM, dropping to 30% in ΔSup35 (Figure 4C-D). In the case of the cells expressing a Sup35 variant, all show a significant increase in the number of white colonies ([*PSI*^+^]) compared to ΔSup35 (Figure 4C-D).

Overall, both assays underscore the significance of the nucleation domain for driving Sup35 aggregation and propagation. They also show that all the amyloid-forming cores selected from the microbiome can trigger Sup35 nucleation and replicate its prion transmission *in vivo* in yeast.

A deeper analysis of the microscopy images shows differences in the colonies and the aggregated forms between Sup35 variants. One characteristic of the [*PSI*^+^] colonies is their color stability, which indicates the capacity of the prion-like protein to keep the aggregated conformation through different generations. We observed that some Sup35 chimeras form long ring intracellular structures (Figure 4B, Supplementary Data 4, Supplementary Figure 18), and that the presence of more than 10% of cells with ring forms is associated with the formation of color revertant colonies. It has been proposed that ring forms are large early aggregated stages that can progress to a mature punctate conformation through the fragmentation activity of HSP104 chaperone^63,64^. In agreement with our results, the formation of long intracellular fibers has been associated with less stable [*PSI^+^*], possibly due to lower amounts of prion seeds (propagons), meanwhile, the punctate foci are associated with the formation of efficient transmissible [*PSI^+^*]^65–67^.

The Sup35 variants C1, C2, C4, C7, and C8 demonstrate the highest percentage of [*PSI*^+^] conversion and, like Sup35NM, robust stability, evident by the absence of revertant colonies. Between them, C1, C2, and C8, also display multiple fluorescent punctate foci, mirroring the pattern observed in Sup35NM (Figure 4B, Supplementary Data 4, Supplementary Figure 18). *In vitro*, upon incubation with SH-5YSY, these three amyloid-forming cores also lead to some of the lowest cell viabilities. In this line, recent studies associated the presence of multiple punctate foci with more toxic variants of the huntingtin protein^61^. Interestingly, C1 and C2 are segments of two different putative VacA from *Helicobacter pylori*, and C8 is a segment of an aggregation promoting factor from a *Lactobacillus delbrueckii* (Supplementary Table 2). Overall, these results support an enhanced toxicity associated with these putative extracellular microbial proteins.

### Aggregation in bacteria-based C-DAG system

We introduced the same set of Sup35NM chimeras expressed in yeast, into the bacteria-based C-DAG system (Figure 5A). In this system, the CsgAss exportation signal facilitates to study protein aggregation propensity by measuring the formation of extracellular aggregates^68^. Importantly, it provides us with a means to simulate protein aggregation within a bacterial context.

**Figure 5.**
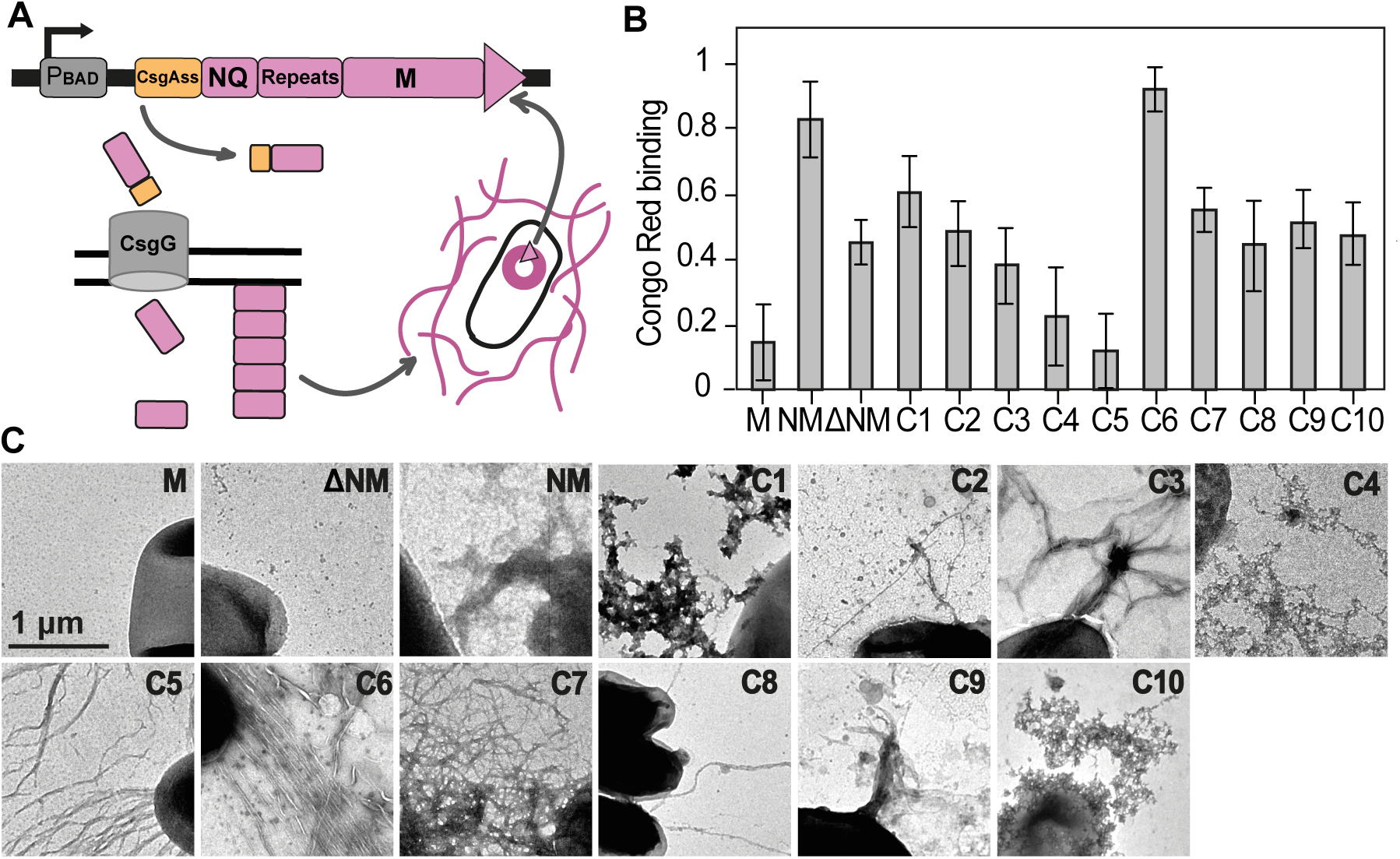
Aggregation in Bacteria. A) Diagram showing the C-DAG construct integrated into E. coli cells and how the expression of Sup35p chimeras can form extracellular protein aggregates. B) Measurement of the colony red intensity using ImageJ, normalized between maximum and minimum. All strains except C3 and C6 show a significant increase in red color in comparison with Sup35M (one-way ANOVA, N=4). C) TEM images showing fibrillar aggregated structures around E. coli cells. D-H) Analysis of the cognitive abilities of C. elegans N2 fed with the E. coli C-DAG collection.

To detect the formation of extracellular aggregates, we grew *E. coli* C-DAG cells on Congo Red (CR) plates, where the presence of extracellular aggregates is revealed by the development of reddish colonies. Accordingly, all strains, except for C4 and C5, displayed a robust signal, notably surpassing that exhibited by Sup35M, which comprises only the M domain (see Figure 5B, Supplementary Figure 19). We also observed, through transmission electron microscopy (TEM), that all strains except Sup35M and ΔSup35 displayed abundant large and fibrillar aggregated structures surrounding the cells (Figure 5C). As observed in yeast, these findings indicate the essential role of the nucleation core in driving Sup35 aggregation, also in a bacterial context.

### Bacteria extracellular fibrils induce loss of associative memory

After demonstrating that there are multiple sequences from the gut microbiome able to form amyloid fibrils and propagate in a prion-like manner, we went a step further and explored whether these proteins could stimulate a neurodegenerative disease phenotype upon ingestion (Figure 1E). To assess this, we focused on monitoring alterations in the sensory memory of two different strains of *C. elegans*: the laboratory wild-type N2 (Figure 6) and the Alzheimer’s disease model CL2355, which has a pan-neuronal expression of Aβ42 (Supplementary Figure 20).

**Figure 6.**
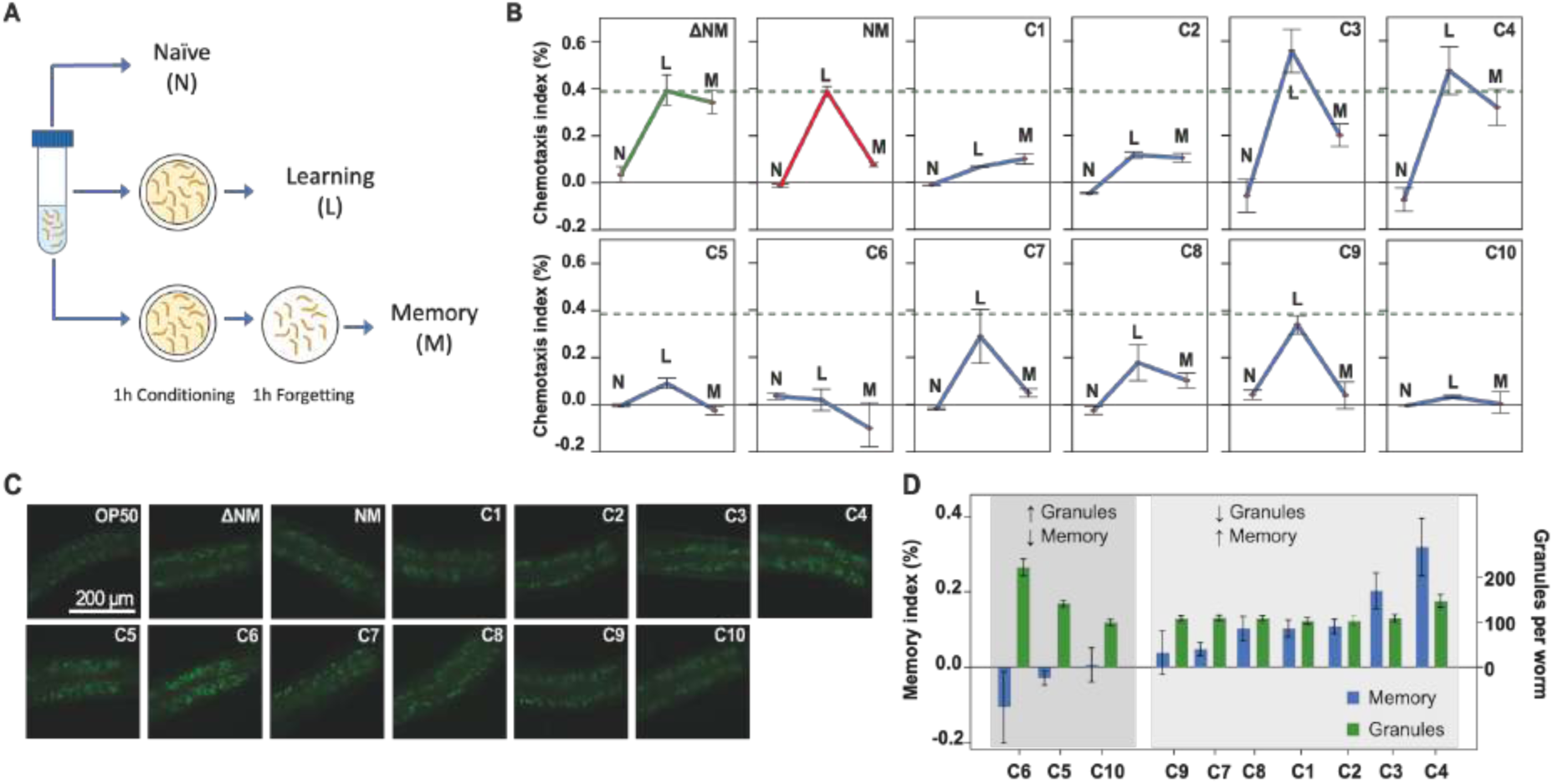
Cognitive Effects on C. elegans. A) Diagram showing the three different conditions of the STAM assay. B) Plots showing the three chemotaxis indexes measured: naïve (N), learning (L) and memory (M). Each data point corresponds to three replicates of 400-600 worms. For individual measurements and comparisons, please refer to Supplementary Figures 22 and 23. The green dashed line indicates the learning index measured when ingesting ΔSup35 (ΔNM). C) Fluorescent images showing the presence of gut granules in the analyzed worms (see Supplementary Figure 24 for transmitted-light images). Fluorescent intensity is indicative of gut granule activity. D) Comparison between memory index and number of gut granules.

We first feed the animals with the *E. coli* C-DAG collection expressing the Sup35NM chimeras (Figure 5A). Then, we performed a short term associative memory assay (STAM) that links the presence of food with the chemoattractant butanone, a volatile odorant (Figure 6A). Thus, we could analyze the cognitive status in those worms^69^.

The STAM employed three replicates of a starting population of approximately 400 to 600 starved worms, which were subsequently divided and subjected to three distinct conditions prior to the execution of the chemotaxis assay (Figure 6A). These conditions included (i) a naïve population that had not been conditioned with the odorant, (ii) a population tested immediately after conditioning (learning index), and (iii) a population tested an hour after conditioning (memory index). The chemotaxis assay itself was conducted on plates where each differently treated population was positioned equidistantly between a spot containing butanone (the chemoattractant) and another spot containing ethanol, a compound to which the worms had not been previously exposed^69^.

We initially compared the phenotype of N2 with the CL2355, to determine whether a STAM assay could be used to measure the ability of *E. coli* C-DAG strains, which form extracellular aggregates, to induce a neurodegenerative disease phenotype (Supplementary Figure 20). Both *C. elegans* models were fed with *E. coli* expressing either the ΔNM or NM variants of Sup35. When compared to N2, CL2355 always presented lower indices. Interestingly, N2 fed with bacteria forming aggregates (Sup35NM), exhibited a memory index similar to that of CL2355 fed with bacteria unable to form extracellular aggregates (ΔSup35). Since both sets of bacteria share the same genotype, these results suggest that the ingestion of the extracellular aggregates produced by Sup35NM causes cognitive impairments in the wild-type animal similar to those associated with the model of Alzheimer’s disease. Furthermore, this assay also showed that the disease phenotype in CL2355 was exacerbated when Sup35NM bacteria were included in its diet.

After validating the STAM assay to measure a neurodegenerative phenotype in N2, we fed the animals with *E. coli* expressing the Sup35NM variants containing the amyloid-forming sequences from the gut microbiota. As a control, we first compared the non-conditioned worms (naïve), and as expected, no significant differences were observed between the animals fed with Sup35NM and those fed with the strains capable of forming extracellular aggregates (Supplementary Figure 21). This indicates that, without prior conditioning, all the animals respond similarly to the chemoattractant. However, during the learning assay (Figure 4E), those animals fed with bacteria expressing C1, C2, C5, C6, and C10 exhibited less than half of the index measured for ΔSup35 or Sup35NM. This suggests a potential decrease in their learning capability due to the consumption of these bacterial strains and their extracellular aggregates.

8 out of 10 memory assays display a significant reduction in the chemotaxis index, compared to ΔSup35 (Figure 6B and Supplementary Figure 22). This suggests that most of the amyloid-forming peptides selected from the gut microbiome can promote the formation of aggregates that, upon ingestion, may lead to a decline in the cognitive abilities of the worm. Putting all together, these assays also indicate that the different aggregates may differentially impact the cognitive abilities of the worm (Figure 6B and Supplementary Figure 23). In this regard, some may have a more pronounced effect on learning (C1, C2, C5, C6, C10), while others may predominantly influence memory (C7, C8, C9)^69^. Intriguingly, these analyses on cognitive abilities highlight the far-reaching influence of the microbiota’s amyloidogenic sequences on cognitive function.

### The aggregates’ properties define the neurodegeneration severity

To analyze the causes of the cognitive decrease, we focused on the gut granules, which are intestinal lysosome-related organelles^70,71^ (Figure 6C, Supplementary Figure 24). These organelles are involved in digestion and nutrient storage and are able to sequester and neutralize ingested toxins and pathogens to prevent damage^72–74^. Recently, they have also been associated with stress response and with the reduction of protein aggregation in a Huntington’s disease model^75^.

The gut granules are autofluorescent, and their intensity can vary with oxidative stress and cellular damage ^76,77^. In our case, the ingestion of the *E. coli* C-DAG collection increased the number of gut granules (Supplementary Figure 25), and their fluorescence intensity (Supplementary Figure 26), compared to the consumption of the *E. coli* OP50 strain, a biofilm-defective mutant^78^. This observation indicates an increase in gut granule activity^79^ and suggests that the *E. coli* C-DAG bacteria may be interfering with the digestion more than the OP50 strain.

When we compare the number of gut granules with the memory index, we observe that those worms with lower memory indexes also have a higher number of gut granules (Figure 6D). This result suggests that the accumulation of components in the gut granules due to the intake of *E. coli* C-DAG may be related to the severity of memory loss. However, this effect is detected to a lesser extent in worms fed with C4, which did not show significant cognitive effects.

Similarly, our results also revealed that the worms ingesting *E. coli* cells, which resulted in the formation of intensely red colonies, exhibited a lower memory index (Supplementary Figure 27), except for C5, which produced a remarkably faint red color. Furthermore, the proteins associated with the formation of abundant ring structures in yeast resulted in lower memory indexes when expressed in *E. coli* and ingested by *C. elegans* (Supplementary Figure 28).

Here we observed a parallel progression between three different assays (number of gut granules, intensity of red colonies, ring aggregates in yeast) and measurements of memory loss (Figure 4H, Supplementary Figures 27-28). The intensity of red in the *E. coli* colonies is linked with the capacity of the extracellular aggregates to bind CR. The ring structures are precursors of foci and their impaired fragmentation is associated with the accumulation of elongated structures and poorer propagation^65,67,80^. Similarly, an increase in gut granules fluorescence has been associated with enhanced activity of these organelles^79^. Collectively, these three assays suggest that specific properties of the aggregates may be linked with the cognitive phenotype of the worms. One theory could be related to the number of aggregates, as indicated by the CR intensity. This may also be associated with the difficulty in degrading and eliminating these aggregates, as suggested by the results in yeast and gut granules. We hypothesize that a higher number of resilient aggregates may decrease the worms’ association memory, which will be interesting to investigate further.

Our research demonstrates the presence of sequences within the gut microbiota that have the potential to form fibrillar aggregates, interfere with host proteins, propagate in a prion-like manner, and influence the cognitive capabilities of the host (Figure 1E). Importantly, our computational analysis also highlights that many of these sequences remain a mystery despite their potential implications in neurodegenerative diseases (Supplementary Figure S1). Furthermore, the analysis of the cognitive effects reveals that aggregates formed from different sequences can have different impacts on associative learning and memory. This underscores the importance of conducting further experiments to understand the mechanisms of these detrimental effects. In this context, our results have led us to hypothesize that differences in the capacity to degrade or fragment these aggregates may be linked to the exacerbation of the disease phenotype. Therefore, the data collected here not only enhances our understanding of the microbiota-host relationship but also unveils potential new targets for the treatment of neurodegenerative diseases.

## Methods

### Screening of Prion-Like Proteins with Amyloid-Forming Cores

The proteins encoded in the gut microbiome’s genome were sourced from the NIH Human Microbiome Project^20,21^. After refining the list to remove duplicate entries, we identified 457 distinct organism names and 1,468,778 protein sequences. This compilation of proteins underwent screening to identify prion-like proteins using three distinct algorithms: PAPA^17^, PLAAC^16^, and pWALTZ^18^. These approaches seek proteins featuring prion-like domains and assign a score to each, indicating the probability of behaving as a prion.

Based on prior studies and our expertise^13^, we identified potential prion-like proteins by employing a combination of criteria from the three different algorithms. Specifically, we employed a PAPA score threshold of 0.05 to denote a positive prion aggregation propensity. Additionally, we considered a PLAAC PRD score above 0 as indicative of sequential domains capable of initiating the aggregation process. Subsequently, we selected 10 candidates for experimental validation, ranking the prion-like sequences according to their pWALTZ scores. This score evaluates the potential of a 21-amino acid sequence to act as an amyloid-forming core, leading to the aggregation of an entire protein (Supplementary Table 1). The pWALTZ algorithm was applied to the 80-amino acid regions previously predicted by PAPA^13^.

### Protein alignment and consensus sequence

The alignment between the 10 different amyloid-forming core candidates and the consensus sequence has been obtained using the Unipro UGENE software v43.0^81^ for sequence visualization.

### Protein Location Prediction using PSORTb

For protein location prediction, we employed PSORTb v3.0^82^. This allowed us to anticipate the potential locations of the protein candidates. The algorithm was executed with ’Bacteria’ selected as the organism type, and the appropriate Gram stain was chosen for each organism. The best Localization Scores were used to generate Supplementary Table 2.

### Peptide preparation

All samples were prepared in low protein-binding microcentrifuge tubes (ThermoFisher Scientific, Waltham, MA, USA).

The amyloid-forming cores were acquired lyophilized from the Peptide Synthesis Facility, Department of Experimental and Health Sciences, Universitat Pompeu Fabra (UPF). Peptides were then solubilized in 1,1,1,3,3,3-hexafluoroisopropanol (HFIP) and separated in different aliquots. Afterward, they were dried O/N under the fume hood at room temperature. Then, different buffers and concentrations were tested to find the best conditions for amyloid aggregate formation (Supplementary Table 1). The best results were obtained by redissolving the peptides with DMSO, to maintain them monomeric, and diluting them in 50 mM Phosphate Buffer (PB) pH 7.4.

Synthetic Aβ40 (DAEFRHDSGYEVHHQKLVFFAEDVGSNKGAIIGLMVGGVV-NH2) was purchased from GenScript (Rijswijk, Netherlands). Stock solutions were prepared by dissolving 1 mg of the peptide to a final concentration of 250 μM and adding 20 mM sodium phosphate buffer, 0.04% NH3, and NaOH to a final pH of 11. Then, the peptide was sonicated for ten minutes (Fisherbrand Pittsburgh, PA, USA, FB15051) without sweep mode and stored at −80 °C until needed. For toxicity and seeding experiments, the peptide was left to aggregate at RT for 24 hours.

### Congo Red binding to protein aggregates

Congo Red (CR) binding to aggregated peptides was analyzed by acquiring the absorbance spectra in the 400-650 nm range using a Cary 300 spectrophotometer (Varian). The assay was performed in a total volume of 150μL (15μL of aggregated peptides samples, 15μL of CR 200μM and 120μL of 50mM PB pH 7.4). Before dye addition, all samples were sonicated for 5 minutes in the ultrasonic bath. Once the dye was added, they were incubated at room temperature for 5 min before the measurement. The peptide spectrums were compared with the spectrum of CR alone, without peptide. To avoid buffer interference, the absorbance spectrum of the buffer was subtracted from all measurements prior to their analysis. A result is considered positive if the maximum absorbance peak of CR shifts towards higher wavelengths.

### Thioflavin-T binding and aggregation kinetics

Thioflavin T was dissolved in Milli-Q water at 5 mM stock, filtered with a 0.2 μm filter, diluted to 0.5 mM, and stored at −20 °C before use. ThT fluorescence for aggregation kinetics was measured every 10 min and at the endpoint for ThT binding of the peptides using a 440 nm excitation filter and a 480 nm emission filter using bottom-optics in a plate reader (TECAN Infinite+ NANO). Samples were placed in a flat-bottom, black, non-binding 96-well plate (Greiner bio-one). A total of 100 µL of sample was added per well. Each condition was measured in triplicate.

For the ThT binding experiments, the fluorescence of the peptides was measured after 72 hours of aggregation of the peptides at a final concentration of 10 μM. Measurements were then compared to the negative control to determine the fold-change in ThT fluorescence intensity.

For the seeding experiments the Aβ40 peptide stock, initially at pH 11, was diluted to 25 µM in sodium phosphate buffer 20 mM and 100 mM NaCl and the corresponding preaggregated peptide was added at 2.5 µM after 5 mins of sonication. Controls without Aβ40 and with peptide and without both Aβ40 and peptides were prepared at the same conditions and by adding the same volumes of the corresponding buffers. All samples and controls were prepared in triplicate. The pH in the wells was measured both at the start and end of the experiment to ensure the pH stability throughout the aggregation process and that there were no differences in the pH of the different conditions. ThT was added to 20 mM final concentration. The aggregation reaction was performed at 37 °C, under quiescent conditions. Four independent replicates were done. The t1/2 is the time necessary, at a given condition, to reach 50% of the final fluorescence signal. The lag phase was calculated by considering its ending as the point at which it reached 10% of the final fluorescence. For each condition, we measured four experimental replicates with three technical replicates, in each.

### Transmission Electron Microscopy

A 10 µl sample of aggregated peptide or *E. coli* (REF) was placed onto carbon-coated copper grids, incubated for 1 min, and dried with Whatman paper. The grids were washed with distilled water, negatively stained with 2% (*w*/*v*) uranyl acetate for 1 min, and then dried. Micrographs were obtained using a JEM-1400 (JEOL, Tokyo, Japan) transmission electron microscope (TEM) operated at an accelerating voltage of 80 keV.

### Infrared Spectroscopy

FT-IR spectra were collected using a Bruker Tensor 27 FT-IR spectrometer (Bruker Optics Inc) with a Golden Gate MKII ATR accessory. Each spectrum consists of 256 independent scans measured at a spectral resolution of 4 cm-1 within the 1400-1800 cm-1 range. To obtain the infrared spectrum of the peptide, the spectrum of the solvent was subtracted from the sample, and all spectra were corrected for the atmospheric water vapor contribution. Each experiment was repeated three times. Afterward, we normalized the results and calculated the 2nd derivative. All data acquisition and analysis were conducted with OPUS MIR Tensor 27 software.

SR-μFTIR of the peptides *in vitro* was performed at the MIRAS beamline at ALBA synchrotron (Catalonia, Spain), using a Hyperion 3000 Microscope that was equipped with a 36× magnification objective coupled to a Vertex 70 spectrometer (Bruker). The measuring range was 650–4000 cm^−1^ and the spectra collection was carried out in transmission mode at 4 cm^−1^ resolution, 10 μm × 10 μm aperture dimensions and coadded from 128–256 scans. Zero filling was performed with fast Fourier transform (FFT), so that, in the final spectra, there was one point every 2 cm^−1^. Background spectra were collected from a clean area of the CaF_2_ window every 15 min. A mercury–cadmium–telluride (MCT) detector was used, and the microscope and spectrometer were continuously purged with nitrogen gas.

### Fourier Transform Infrared (FTIR) Spectra Analysis

Spectra were acquired in two different ways: (a) at least 50 spectra on single cells were acquired for each sample in a given sample region; (b) maps with a dimension of minimum 50 μm × 50 μm with a step size of 6 μm × 6 μm. Fourier transform infrared (FTIR) spectra of single independent cells, the spectra from the different cell maps, and the independent spectra of amyloid aggregates without cells were analyzed using Thermo Omnic 7.1 (Thermo Scientific, Inc.) and Opus 7.5 (Bruker) software. The spectra exhibiting a low signal-to-noise ratio were eliminated. For data processing, the second derivative of the spectra was calculated using a Savitsky–Golay algorithm with a nine-point filter and a polynomial order of 2, to eliminate the baseline contribution. Unscrambler X software (CAMO Software, Oslo, Norway) was used to perform PCA in the dataset. PCA analysis was applied for the second derivative of the spectra. Unit vector normalization was applied after secondary derivation for PCA analysis. Principal components (PCs) were calculated using the Nonlinear Iterative Partial Least Squares (NIPALS) algorithm on mean centered data. Since the PCA procedure allows weighting the individual variables relative to each other, a constant value (1.00, equal weight) was assigned to all variables (the different wavenumbers in the 650–4000 cm^−1^ region), as the recommended value. For Ratios were calculated over the following peaks of interest: 1627 cm^−1^ for Amide I β-sheet structures (noted as A_1627_), 1665 cm^−1^ as a wavelength at which both the amyloid peptide and the cells have (in the derivative spectrum) a signal clearly distinct from zero (noted as A_1665_), 1740 cm^−1^ for ν(C═O) (carbonyl) (noted as A_1740_), 2925 cm^−1^ for CH_2_ asymmetric stretching vibrations (noted as A_2925_), and 2960 cm^−1^ for CH_3_ asymmetric stretching vibrations (noted as A_2960_). It is important to stress out that, since the peptide alone signal at 1657 cm^−1^ in the derivative spectra is close to zero (Figure SX), choosing the 1665 cm^−1^ wavelength permits one to estimate peptide aggregation as the ratio A_1627_/A_1665_. Origin 9.1 software was used for the ratios calculation, *t*-test analysis, and graphical representation.

### Cell Cytotoxicity Assay

Neuroblastoma SH-SY5Y cells were cultured in Dulbecco’s modified Eagle’s medium, F-12 supplemented with Glutamax (DMEM/F12 Glutamax) supplemented with 10% (v/v) heat-inactivated fetal bovine serum and 1% (v/v) penicillin/streptomycin. The cells were maintained at 37 °C and 5% CO2 in a 75 cm2 cell culture flask. Differentiation to neuronal cells started 24 h after plating by replacement of the maintenance medium with differentiation culture medium for 7 days and refreshment every 72 h. The differentiation culture medium consisted of DMEM/F-12 Glutamax supplemented with 2.5% inactivated FBS and 10 μM retinoic acid. Differentiation was monitored microscopically by morphological assessment.

For the cytotoxicity assay, the cells were seeded in 96-well plates and treated at a density of 1 × 104 cells/well. The different peptides were added at the determined concentrations (10-0.25 μM), and after 24 h of incubation, cell viability was detected by MTT assay. After removing the medium, 10 μL of 3-(4,5-dimethylthiazol-2-yl)-2,5-diphenyltetrazolium bromide (MTT) solution (5 mg/mL) and 100μL of the medium were added to each well and incubated at 37 °C for 4 h. Then, 150 μL of dimethyl sulfoxide was added to each well to dissolve the formazan after discarding the supernatant. Absorbance values were quantified using a plate reader (TECAN Spark) at a wavelength of 580 nm. For each pH, the data are expressed as a percentage of viability with respect to non-treated cells. The non-treated cells were grown in a medium containing the same amount of buffer, at the corresponding pH, but without Aβ40 nor the corresponding peptide.

For the ROS assay, the cells were seeded in 96-well plates and treated at a density of 1x 10^4^ cells / well. The different peptides were added at 10 μM and after 4h of incubation, ROS were detected by DCFDA/H2DCFDA assay.

### Expression and purification

A pET21b vector (Novagen) containing the full sequence of C9L6N5 (C4) with a C-terminal His-tag was introduced into *E. coli* BL21. The bacteria were grown in LB at 37 °C. At an OD600 of 0.5, IPTG 1mM was added and then the culture was left incubating O/N at 22 °C. Afterward, the cells were centrifugated, and the pellet was resuspended in PBS with 5 mM imidazole, RNAse A, and DNAse I and sonicated. Then the lysate was ultracentrifuged, and we kept the supernatant which was added to a His-Trap FF (Cytiva, Barcelona, Spain) column. Which was washed with a gradient of PBS with imidazole buffer, reaching a maximum concentration of imidazole of 200 mM. Afterward, the protein purity was assessed with a SDS-PAGE gel and a Western blot to ensure that it was the protein of interest.

The purified samples were separated by SDS-PAGE gels in duplicate, one was stained with Coomassie Blue and the other was transferred to nitrocellulose membranes. After the transference, the membranes were blocked with 4% bovine serum albumin. Then, the membrane was incubated with an anti-His Rabbit primary antibody (Thermo Scientific, Massachusetts, United States) (1:1000). The secondary antibody was an anti-rabbit horseradish peroxidase conjugate (Bio-Rad) (1:3000). The reaction was developed with an ECL kit (Thermo Scientific).

### Yeast strain and non-sense suppression system

The yeast strain used in this project was derived from 74D-694 *MATa, ade1-14UGA, trp1-289, his3*1′*-200, ura3-52, leu2-3,112 sup35::loxP* [pYK810]) (REF). The N-terminal Sup35p modified chimeras (where the first 40 residues are replaced with 21 residues peptides) are encoded in a centromeric pUCK1620 plasmid bearing a *HIS3* selection marker.

In our assay, the full-length Sup35 version and their peptide-containing chimeras (Figure 3A) are expressed constitutively under their respective promoters at physiological levels to maintain essential terminator factor activity. To prompt aggregation, an additional copy of only the N and M domains fused to GFP is transiently overproduced under the control of a galactose-inducible promoter. This strategy allows us to induce prion conversion when necessary and simultaneously monitor protein aggregation using fluorescent microscopy. The initial 40 residues of Sup35p, and the NM domain fused to GFP, were replaced with the different selected amyloid cores (Figure 3A).

### Fluorescent foci detection in yeast

Yeast cells grown for 3–5 days on SD medium plates were inoculated into a synthetic medium with raffinose (instead of glucose) and without histidine nor uracil (SRaf -His, -Ura), and subsequently grown at 30 °C under vigorous agitation for 3 days. To induce expression of NM-GFP chimeras, yeast was inoculated into a synthetic medium with 2% galactose and 2% raffinose (instead of glucose) one day before the GFP assay, as they are under the *GAL10* promoter. On the day of the experiment, the cells were examined and recorded using fluorescence microscopy (EVOS M5000) on a 96-well plate.

### Non-sense suppression assay

In this assay, a preculture was initiated in SD -His -Ura medium, using 2% glucose as the carbon source, for three days at 30 °C. This three-day interval serves to repress expression prior to commencing the assay. Subsequently, yeast cells were washed with PBS buffer and induced to express the NMGFP fusions by inoculating fresh media containing 2% galactose and 2% raffinose, and incubated for two days at 30 °C. All cultures were adjusted to an OD_600_ of 1 and plated at a 1/1000 dilution on ¼ YPD plates (1% bacto-yeast extract, 2% bacto-peptone, 2% glucose, 2% bacto-agar), then incubated for 4–5 days at 30 °C. After this incubation, plates were transferred to 4°C for an additional 5-7 days and subsequently examined for the red and white colony phenotypes. Images were captured for each plate using a stereomicroscope Leica MZFLIII (Leica Microsystems GmbH, Mannheim, Germany), set at 8x magnification. All assays were conducted in triplicate at a minimum, with colony counts exceeding 200 per plate. Colony counting was executed for each assay, followed by t-student analysis for comparison against the negative control. Red colonies were indicative of the [*psi*^-^] phenotype, while white colonies indicated a stable [*PSI*^+^] phenotype characterized by efficient transmission between cells. Pink colonies were observed as indicative of an unstable [*PSI*^+^] phenotype, attributed to factors including low transmission between cells and/or high reversion between the aggregated and soluble forms of Sup35p.

### Colony-color phenotype assay to detect extracellular aggregates

For the expression and extracellular aggregation of the Sup35p chimeras, we utilized the *E. coli* strain VS39 (kindly provided by Ann Hochschild). This particular strain is deficient in curli genes (*csgA*, *csgB*, and *csgC*), resistant to kanamycin, and contains a pACYC-derived plasmid named pVS76. This plasmid orchestrates the synthesis of the outer-membrane curli protein, CsgG, regulated by an IPTG-inducible promoter. Additionally, VS39 carries a cat gene, constitutively transcribed, which provides resistance to chloramphenicol.

The *Sup35NM* genes were obtained by modifying the pEXPORT plasmid pVS72, which harbors an intact *Sup35NM*. The finalized plasmids were subsequently transformed into the VS39 strain. The detailed sequence information of the final constructs is provided in the Supplementary Material. In this study, we also incorporated the pEXPORT plasmid pVS105, encoding for the Sup35 medium region (*Sup35M*) gene.

To measure the capacity of forming extracellular aggregates among the various Sup35p variants, bacterial cells were spotted on CR-inducing plates. These plates consist of LB agar supplemented with 100 µg/ml carbenicillin, 25 µg/ml chloramphenicol, 0.2% w/v L-arabinose, 1 mM IPTG, and 10 µg/ml Congo Red (REF). Following a 5-day incubation period at 22°C, the plates were subjected to imaging to assess the extent of CR binding. To obtain quantitative data, red color intensity values were derived from quadruplicate samples using ImageJ and normalized to Sup35ΔNM as in Brunquell, et al. 2018^75^.

### *C. elegans* strains and maintenance

In this study, we utilized the wild-type N2 strain, which was obtained from the Caenorhabditis Genetics Center (CGC). The worms were maintained at a temperature of 20 °C on standard nematode growth medium (NGM) plates that had been seeded with E. coli OP50 as their food source. To synchronize the nematode population, we employed the conventional method of subjecting them to a 5% hypochlorite treatment, followed by a 24-hour rotation at 80 rpm in M9 buffer without any food source. Following this procedure and until they reached the young adult stage, each group of worms was incubated with the respective bacterial strain.

### *C. elegans* butanone associative short-term learning assay

The butanone associative learning assay was performed following established methods (REF-6). Young adults were washed in M9 and collected by gravity sedimentation in a 15 mL conical tube to remove bacteria. Some animals were immediately transferred to the chemotaxis plate for assaying the naïve condition (CI (naïve)). The rest were starved in M9 for 1 hour. After the starvation, animals were conditioned for 1 hour on NGM plates seeded with the corresponding *E. coli* strain and containing a 2 μL drop of 10% butanone, diluted in absolute ethanol, on the lid.

The conditioned worms (right after the condition, Learning index (LI), or after 1 hour of starvation, Memory index (LI)) were washed with M9 and transferred to the chemotaxis plate. For the chemotaxis assay, 10 cm unseeded NGM plates were used, and three circles (each 1 cm in diameter) on the bottom and both sides of the plate were marked. Then 1 μL of 1 M sodium azide and 1 μL of 10% of butanone were added to the left spot, and the same amount of sodium azide and pure ethanol control were added to the right spot. The animals were placed in the bottom spot. After one hour, the number of worms located in the butanone and ethanol spots, as well as at the original bottom spots were counted. Each chemotaxis assay was performed in triplicates and contained about 400-600 worms.

Our standard formula to calculate a chemotaxis index (CI) is:

CI = [(Number of worms in the butanone zone) – (Number of worms in the ethanol zone)] / (Total number of worms)

With worms not conditioned, we calculated the CI (naïve):

CI (naïve) = [(Number of worms in the test butanone zone) – (Number of worms in the ethanol zone)] / (Total number of worms)

Right after conditioning, we calculated the Learning index (LI) as:

Learning index (LI) = CI (right after conditioning) – CI (naïve)

And after conditioning followed by 1 hour of starvation, we calculated the Memory index (LI) as:

Memory index (LI): CI (after conditioning and starvation) – CI (naïve).

### *C. elegans* gut granules autofluorescence detection

The EVOS microscope (EVOSTM M5000 Imaging System) was employed to capture autofluorescence images of *C. elegans* using a Green Fluorescent Protein (GFP) filter and a Transmission filter. The images were acquired from animals treated identically to those used in the chemotaxis assays. For visualizing gut granules, we centered the images on the first two intestine rings, which correspond to the region with the highest fluorescence intensity. Subsequently, ImageJ software was utilized for image analysis. The selected region for analysis was carefully chosen to avoid the presence of the gonads, mitigating potential interference.

### Statistics

The statistics were realized with GraphPad PRISM 8 software and the corresponding statistical analysis for the obtained data. In general, the data was analyzed with an Ordinary One-way ANOVA. Statistical significance was considered with p values lower than 0.05 and the data was shown as the Mean ± SEM.

## Acknowledgments

We extend our gratitude to the following individuals:

Ann Hochschild and Padraig Deighan for kindly providing the SV39 *E. coli* strain and plasmids pVS105 and PVS72.

Rafael Giraldo for generously providing the 74D-694 S. cerevisiae strain. Antonela Lavatelli for her assistance in resolving basic experimental issues.

Julian Cerón and Antonio Miranda-Vizuete for their invaluable responses to our inquiries.

Anna Laromaine for sharing her *C. elegans* laboratory expertise and aiding in the construction of the initial picker.

Esther Dalfo for enlightening us about the requirements of a *C. elegans* laboratory and for sharing valuable information regarding contacts and databases related to *C. elegans*.

Gian G. Tartaglia for his inestimable support to start the project. Nuria Benseny for her assistance in the microFTIR experiments.

Josep Cladera for his responses and discussions around the investigation.

*C. elegans* N2 and *E. coli* OP50 were provided by the CGC, which is funded by NIH Office of Research Infrastructure Programs (P40 OD010440).

μFTIR experiments were performed at MIRAS beamline at ALBA Synchrotron with the collaboration of ALBA staff.

## Author contributions

NSG conceived the research idea and developed the study design. MRF, JSC, ADM, MSG, MGP collected and compiled the primary data for the study. NSG, MRF, JSC conducted the statistical analysis and interpreted the results. NSG, MRF, JSC wrote the initial draft of the manuscript. NSG, MRF, JSC, ADM assisted in refining the research methodology. NSG and MRF provided oversight and guidance throughout the research project.

## References

1. Duyckaerts, C., Clavaguera, F. & Potier, M. C. The prion-like propagation hypothesis in Alzheimer’s and Parkinson’s disease. Curr. Opin. Neurol. 32, 266–271 (2019).

2. Ritchie, D. L. & Barria, M. A. Prion Diseases: A Unique Transmissible Agent or a Model for Neurodegenerative Diseases? Biomolecules 11, 1–23 (2021).

3. Fang, P., Kazmi, S. A., Jameson, K. G. & Hsiao, E. Y. The Microbiome as a Modifier of Neurodegenerative Disease Risk. Cell Host Microbe 28, 201–222 (2020).

4. Bauer, K. C., Huus, K. E. & Finlay, B. B. Microbes and the mind: Emerging hallmarks of the gut microbiota-brain axis. Cell. Microbiol. 18, 632–644 (2016).

5. Bonaz, B., Bazin, T. & Pellissier, S. The vagus nerve at the interface of the microbiota-gut-brain axis. Front. Neurosci. 12, 49 (2018).

6. Svensson, E. et al. Vagotomy and subsequent risk of Parkinson’s disease. Ann. Neurol. 78, 522–529 (2015).

7. Friedland, R. P. & Chapman, M. R. The role of microbial amyloid in neurodegeneration. PLoS Pathog. 13, (2017).

8. Thapa, M. et al. Translocation of gut commensal bacteria to the brain. bioRxiv 2023.08.30.555630 (2023) doi:10.1101/2023.08.30.555630.

9. Hashim, H. M. & Makpol, S. A review of the preclinical and clinical studies on the role of the gut microbiome in aging and neurodegenerative diseases and its modulation. Front. Cell. Neurosci. 16, (2022).

10. Sampson, T. R. et al. Gut Microbiota Regulate Motor Deficits and Neuroinflammation in a Model of Parkinson’s Disease. Cell 167, 1469–1480.e12 (2016).

11. Sampson, T. R. et al. A gut bacterial amyloid promotes α-synuclein aggregation and motor impairment in mice. Elife 9, (2020).

12. Seira Curto, J., et al. Microbiome Impact on Amyloidogenesis. Front. Mol. Biosci. 9, (2022).

13. Iglesias, V., de Groot, N. S. & Ventura, S. Computational analysis of candidate prion-like proteins in bacteria and their role. Front. Microbiol. 6, (2015).

14. Espinosa Angarica, V., Ventura, S. & Sancho, J. Discovering putative prion sequences in complete proteomes using probabilistic representations of Q/N-rich domains. BMC Genomics 14, 1–17 (2013).

15. Harrison, P. M. Evolutionary behaviour of bacterial prion-like proteins. PLoS One 14, e0213030 (2019).

16. Lancaster, A. K., Nutter-Upham, A., Lindquist, S. & King, O. D. PLAAC: A web and command-line application to identify proteins with prion-like amino acid composition. Bioinformatics 30, 2501–2502 (2014).

17. Toombs, J. A. et al. De novo design of synthetic prion domains. Proc. Natl. Acad. Sci. U. S. A. 109, 6519–6524 (2012).

18. Sabate, R., Rousseau, F., Schymkowitz, J. & Ventura, S. What Makes a Protein Sequence a Prion? PLOS Comput. Biol. 11, e1004013 (2015).

19. Gil-Garcia, M., Iglesias, V., Pallarès, I. & Ventura, S. Prion-like proteins: from computational approaches to proteome-wide analysis. FEBS Open Bio 11, 2400–2417 (2021).

20. University of Maryland. NIH Human Microbiome Project. Microbe Magazine vol. 4 393–393 https://www.hmpdacc.org/hmp/ (2009).

21. Park, S. K. R. et al. ComPIL 2.0: An Updated Comprehensive Metaproteomics Database. J. Proteome Res. 18, 616–622 (2019).

22. Visconti, A. et al. Interplay between the human gut microbiome and host metabolism. Nat. Commun. *2019 101* 10, 1–10 (2019).

23. Ijaq, J., Chandrasekharan, M., Poddar, R., Bethi, N. & Sundararajan, V. S. Annotation and curation of uncharacterized proteins-challenges. Front. Genet. 6, 115944 (2015).

24. Desler, C., Durhuus, J. A. & Rasmussen, L. J. Genome-wide screens for expressed hypothetical proteins. Methods Mol. Biol. 815, 25–38 (2012).

25. Javed, I. et al. Accelerated Amyloid Beta Pathogenesis by Bacterial Amyloid FapC. Adv. Sci. 7, 2001299 (2020).

26. Hartman, K. et al. Bacterial curli protein promotes the conversion of PAP248-286 into the amyloid SEVI: cross-seeding of dissimilar amyloid sequences. PeerJ 1, (2013).

27. De Groot, N. S. & Burgas, M. T. Is membrane homeostasis the missing link between inflammation and neurodegenerative diseases? Cell. Mol. Life Sci. 2015 7224 72, 4795–4805 (2015).

28. Marquer, C. et al. Local cholesterol increase triggers amyloid precursor protein-Bacel clustering in lipid rafts and rapid endocytosis. FASEB J. 25, 1295–1305 (2011).

29. Malnar, M. et al. Cholesterol-depletion corrects APP and BACE1 misstrafficking in NPC1-deficient cells. Biochim. Biophys. Acta - Mol. Basis Dis. 1822, 1270–1283 (2012).

30. Rivest, S. Regulation of innate immune responses in the brain. Nat. Rev. Immunol. *2009 96* 9, 429–439 (2009).

31. Chen, S. G. et al. Exposure to the Functional Bacterial Amyloid Protein Curli Enhances Alpha-Synuclein Aggregation in Aged Fischer 344 Rats and Caenorhabditis elegans. Sci. Reports 2016 61 6, 1–10 (2016).

32. Friedland, R. P., McMillan, J. D. & Kurlawala, Z. What Are the Molecular Mechanisms by Which Functional Bacterial Amyloids Influence Amyloid Beta Deposition and Neuroinflammation in Neurodegenerative Disorders? Int. J. Mol. Sci. *2020, Vol. 21, Page 1652* 21, 1652 (2020).

33. Van Gerven, N., Van der Verren, S. E., Reiter, D. M. & Remaut, H. The Role of Functional Amyloids in Bacterial Virulence. J. Mol. Biol. 430, 3657–3684 (2018).

34. Kuwajima, K. et al. Functional Bacterial Amyloids: Understanding Fibrillation, Regulating Biofilm Fibril Formation and Organizing Surface Assemblies. Mol. *2022, Vol. 27, Page 4080* 27, 4080 (2022).

35. Kosolapova, A. O., Antonets, K. S., Belousov, M. V. & Nizhnikov, A. A. Biological Functions of Prokaryotic Amyloids in Interspecies Interactions: Facts and Assumptions. Int. J. Mol. Sci. 21, 1–26 (2020).

36. Wang, C., Lau, C. Y., Ma, F. & Zheng, C. Genome-wide screen identifies curli amyloid fibril as a bacterial component promoting host neurodegeneration. Proc. Natl. Acad. Sci. U. S. A. 118, (2021).

37. Miller, A. L., Bessho, S., Grando, K. & Tükel, Ç. Microbiome or Infections: Amyloid-Containing Biofilms as a Trigger for Complex Human Diseases. Front. Immunol. 12, 638867 (2021).

38. Vogt, N. M. et al. Gut microbiome alterations in Alzheimer’s disease. Sci. Rep. 7, (2017).

39. Go, M. F. Natural history and epidemiology of Helicobacter pylori infection. Aliment. Pharmacol. Ther. 16, 3–15 (2002).

40. Panza, F., Lozupone, M., Solfrizzi, V., Watling, M. & Imbimbo, B. P. Time to test antibacterial therapy in Alzheimer’s disease. Brain 142, 2905–2929 (2019).

41. Khaled, M. et al. A Hairpin Motif in the Amyloid-β Peptide Is Important for Formation of Disease-Related Oligomers. J. Am. Chem. Soc. 145, 18340–18354 (2023).

42. Pallarès, I., Iglesias, V. & Ventura, S. The rho termination factor of Clostridium botulinum contains a prion-like domain with a highly amyloidogenic core. Front. Microbiol. 6, 177351 (2016).

43. Huttenhower, C. et al. Structure, function and diversity of the healthy human microbiome. Nat. *2012 4867402* 486, 207–214 (2012).

44. Cammann, D. et al. Genetic correlations between Alzheimer’s disease and gut microbiome genera. Sci. Reports *2023 131* 13, 1–15 (2023).

45. Haran, J. P. et al. Alzheimer’s Disease Microbiome Is Associated with Dysregulation of the Anti-Inflammatory P-Glycoprotein Pathway. MBio 10, (2019).

46. Xia, Y. Correlation and association analyses in microbiome study integrating multiomics in health and disease. Prog. Mol. Biol. Transl. Sci. 171, 309–491 (2020).

47. Tenaillon, O., Skurnik, D., Picard, B. & Denamur, E. The population genetics of commensal Escherichia coli. Nat. Rev. Microbiol. *2010 83* 8, 207–217 (2010).

48. Varadi, M. et al. AlphaFold Protein Structure Database: massively expanding the structural coverage of protein-sequence space with high-accuracy models. Nucleic Acids Res. 50, D439–D444 (2022).

49. Liu, H. et al. Negatively charged hydrophobic nanoparticles inhibit amyloid β-protein fibrillation: The presence of an optimal charge density. React. Funct. Polym. 103, 108– 116 (2016).

50. Liao, Y. H., Chang, Y. J., Yoshiike, Y., Chang, Y. C. & Chen, Y. R. Negatively charged gold nanoparticles inhibit Alzheimer’s amyloid-β fibrillization, induce fibril dissociation, and mitigate neurotoxicity. Small 8, 3631–3639 (2012).

51. Chan, H. M. et al. Effect of surface-functionalized nanoparticles on the elongation phase of beta-amyloid (1–40) fibrillogenesis. Biomaterials 33, 4443–4450 (2012).

52. Seira Curto, J., Fernandez, M. R., Cladera, J., Benseny-Cases, N. & Sanchez de Groot, N. Aβ40 Aggregation under Changeable Conditions. Int. J. Mol. Sci. 24, 8408 (2023).

53. Wood, S. J., Maleeff, B., Hart, T. & Wetzel, R. Physical, Morphological and Functional Differences between pH 5.8 and 7.4 Aggregates of the Alzheimer’s Amyloid Peptide A β. J. Mol. Biol. 256, 870–877 (1996).

54. Kremer, J. J., Pallitto, M. M., Sklansky, D. J. & Murphy, R. M. Correlation of β-amyloid aggregate size and hydrophobicity with decreased bilayer fluidity of model membranes. Biochemistry 39, 10309–10318 (2000).

55. Mannini, B. et al. Toxicity of Protein Oligomers Is Rationalized by a Function Combining Size and Surface Hydrophobicity. ACS Chem. Biol 9, (2014).

56. Van Schependom, J. & D’haeseleer, M. Advances in Neurodegenerative Diseases. J. Clin. Med. 12, (2023).

57. Foo, J. H. et al. Both the p33 and p55 Subunits of the Helicobacter pylori VacA Toxin Are Targeted to Mammalian Mitochondria. J. Mol. Biol. 401, 792–798 (2010).

58. Su, M. et al. Cryo-EM Analysis Reveals Structural Basis of Helicobacter pylori VacA Toxin Oligomerization. J. Mol. Biol. 431, 1956–1965 (2019).

59. Sivanathan, V. & Hochschild, A. A bacterial export system for generating extracellular amyloid aggregates. Nat. Protoc. *2013 87* 8, 1381–1390 (2013).

60. Sivanathan, V. & Hochschild, A. Generating extracellular amyloid aggregates using E. coli cells. Genes Dev. 26, 2659–2667 (2012).

61. Parham, S. N., Resende, C. G. & Tuite, M. F. Oligopeptide repeats in the yeast protein Sup35p stabilize intermolecular prion interactions. EMBO J. 20, 2111–2119 (2001).

62. Wickner, R. B. et al. Yeast Prions: Structure, Biology, and Prion-Handling Systems. Microbiol. Mol. Biol. Rev. 79, 1–17 (2015).

63. Tyedmers, J. et al. Prion induction involves an ancient system for the sequestration of aggregated proteins and heritable changes in prion fragmentation. Proc. Natl. Acad. Sci. U. S. A. 107, 8633–8638 (2010).

64. Sharma, J. et al. De novo [PSI +] prion formation involves multiple pathways to form infectious oligomers. Sci. Rep. 7, 76 (2017).

65. Chernova, T. A., Wilkinson, K. D. & Chernoff, Y. O. Prions, Chaperones, and Proteostasis in Yeast. Cold Spring Harb. Perspect. Biol. 9, a023663 (2017).

66. Liebman, S. W. & Chernoff, Y. O. Prions in yeast. Genetics 191, 1041–1072 (2012).

67. Derkatch, I. L. & Liebman, S. W. The story of stolen chaperones. Prion 7, 294–300 (2013).

68. Zhou, Y., Smith, D. R., Hufnagel, D. A. & Chapman, M. R. Experimental manipulation of the microbial functional amyloid called curli. Methods Mol. Biol. 966, 53–75 (2013).

69. Stein, G. M. & Murphy, C. T. C. elegans positive olfactory associative memory is a molecularly conserved behavioral paradigm. Neurobiol. Learn. Mem. 115, 86–94 (2014).

70. Hermann, G. J. et al. Genetic analysis of lysosomal trafficking in Caenorhabditis elegans. Mol. Biol. Cell 16, 3273–3288 (2005).

71. Dell’Angelica, E. C., Mullins, C., Caplan, S. & Bonifacino, J. S. Lysosome-related organelles. FASEB J. 14, 1265–1278 (2000).

72. Roh, H. C., Collier, S., Guthrie, J., Robertson, J. D. & Kornfeld, K. Lysosome-related organelles in intestinal cells are a zinc storage site in C. elegans. Cell Metab. 15, 88–99 (2012).

73. Chun, H. et al. The Intestinal Copper Exporter CUA-1 Is Required for Systemic Copper Homeostasis in Caenorhabditis elegans. J. Biol. Chem. 292, 1–14 (2017).

74. Ardelli, B. F. & Prichard, R. K. Inhibition of P-glycoprotein enhances sensitivity of Caenorhabditis elegans to ivermectin. Vet. Parasitol. 191, 264–275 (2013).

75. Brunquell, J., Morris, S., Snyder, A. & Westerheide, S. D. Coffee extract and caffeine enhance the heat shock response and promote proteostasis in an HSF-1-dependent manner in Caenorhabditis elegans. Cell Stress Chaperones 23, 65–75 (2018).

76. Coburn, C. et al. Anthranilate fluorescence marks a calcium-propagated necrotic wave that promotes organismal death in C. elegans. PLoS Biol. 11, (2013).

77. Coburn, C. & Gems, D. The mysterious case of the c. elegans gut granule: Death anthranilic acid and the kynurenine pathway. Front. Genet. 4, 57351 (2013).

78. Arata, Y., Oshima, T., Ikeda, Y., Kimura, H. & Sako, Y. OP50, a bacterial strain conventionally used as food for laboratory maintenance of C. elegans, is a biofilm formation defective mutant. microPublication Biol. (2020) doi:10.17912/micropub.biology.000216.

79. Huynh, J., Dang, H. & Fares, H. Measurement of Lysosomal Size and Lysosomal Marker Intensities in Adult Caenorhabditis elegans. BIO-PROTOCOL 8, (2018).

80. Barbitoff, Y. A., Matveenko, A. G. & Zhouravleva, G. A. Differential Interactions of Molecular Chaperones and Yeast Prions. J. Fungi *2022, Vol. 8, Page 122* 8, 122 (2022).

81. Okonechnikov, K., Golosova, O., Fursov, M. & UGENE team. Unipro UGENE: a unified bioinformatics toolkit. Bioinformatics 28, 1166–7 (2012).

82. Yu, N. Y. et al. PSORTb 3.0: improved protein subcellular localization prediction with refined localization subcategories and predictive capabilities for all prokaryotes. Bioinformatics 26, 1608–1615 (2010).

