## Supplementary Material for "Microbiome-Derived Prion-Like Proteins and Their Potential to Trigger Cognitive Dysfunction"

**Index**

**Supplementary Figures**


| | | | |
|---|---|---|---|
| Supplementary Figure 1 | **... Page 2** | Supplementary Figure 15 | **... Page 16** |
| Supplementary Figure 2 | **... Page 3** | Supplementary Figure 16 | **... Page 17** |
| Supplementary Figure 3 | **... Page 4** | Supplementary Figure 17 | **... Page 18** |
| Supplementary Figure 4 | **... Page 5** | Supplementary Figure 18 | **... Page 19** |
| Supplementary Figure 5 | **... Page 6** | Supplementary Figure 19 | **... Page 20** |
| Supplementary Figure 6 | **... Page 7** | Supplementary Figure 20 | **... Page 21** |
| Supplementary Figure 7 | **... Page 8** | Supplementary Figure 21 | **... Page 22** |
| Supplementary Figure 8 | **... Page 9** | Supplementary Figure 22 | **... Page 23** |
| Supplementary Figure 9 | **... Page 10** | Supplementary Figure 23 | **... Page 24** |
| Supplementary Figure 10 | **... Page 11** | Supplementary Figure 24 | **... Page 25** |
| Supplementary Figure 11 | **... Page 12** | Supplementary Figure 25 | **... Page 26** |
| Supplementary Figure 12 | **... Page 13** | Supplementary Figure 26 | **... Page 27** |
| Supplementary Figure 13 | **... Page 14** | Supplementary Figure 27 | **... Page 28** |
| Supplementary Figure 14 | **... Page 15** | Supplementary Figure 28 | **... Page 29** |


**Supplementary Data**


**Supplementary Data 1.** Table showing ten amyloid-forming cores selected from gut microbiome. **Page 30**

**Supplementary Data 2.** Predictions performed on the Sup35p variants studied in this work. **Pages 31 to 43**

**Supplementary Data 3.** Protein sequences of the Sup35p and the Sup35NM fused to GFP expressed in *S. cerevisiae*. **Page 44**

**Supplementary Data 4.** Table showing the phenotypic characteristics associated to Sup35 expression. **Page 45**

**Supplementary Data 5.** Sequence of the Sup35NM variants (Sup35NM, DNM and chimeras) expressed in the C-DAG *E. coli* system. **Page 46**

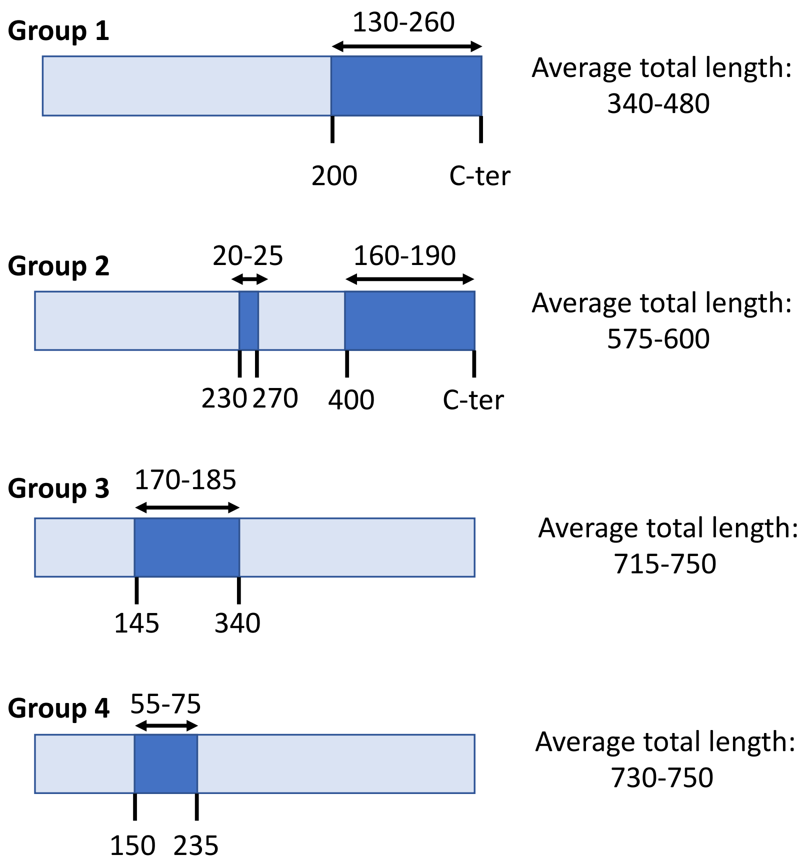


**Supplementary Figure S1. Common Disordered Regions Identified in Uncharacterized Proteins.** The diagram displays four sequences, highlighting the consensus locations identified within the gut microbiome. This classification is based on information sourced from MobiDB Lite (Supplementary Table 1). In Supplementary Table 1, there are 370 uncharacterized sequences, constituting 40% of the dataset. Among these, 214 sequences have been determined to contain disordered regions according to MobiDB Lite. The groups shown in this figure consist of the following numbers of sequences: Group 1, 78 sequences; Group 2, 39 sequences; Group 3, 35 sequences; Group 4, 23 sequences.

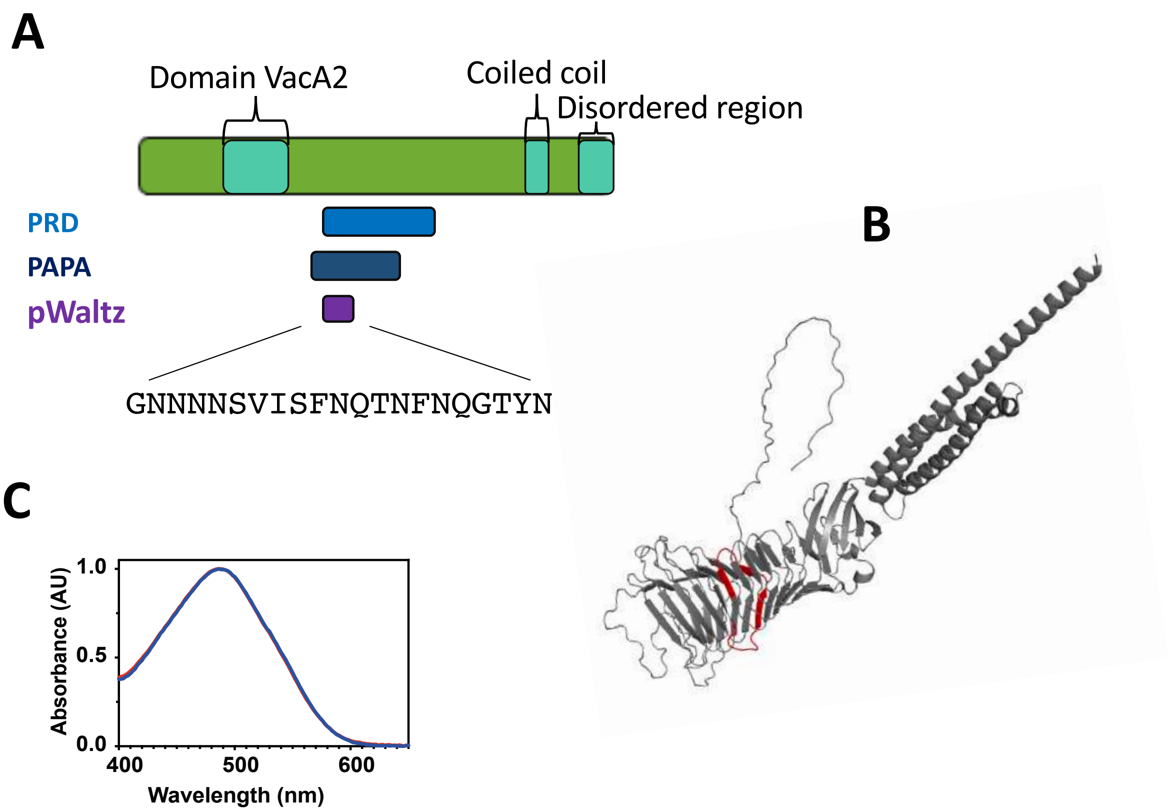


**Supplementary Figure 2. Predictive and experimental results associated to M5YJZ4 containing C1.** A) Diagram sowing the prion-like regions predicted. B) Alpha-fold prediction of the whole protein structure. C) Congo-red binding analysis.

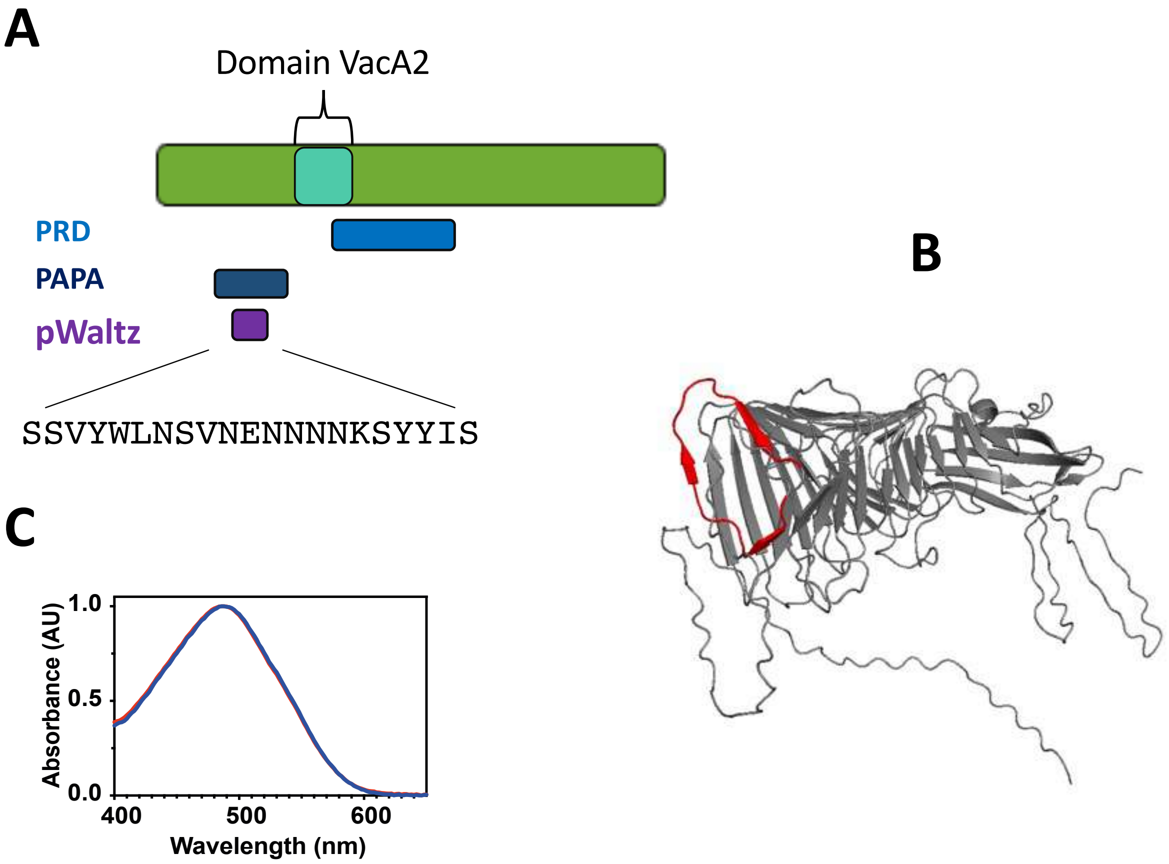


**Supplementary Figure 3. Predictive and experimental results associated to M3SJI9 containing C2.** A) Diagram sowing the prion-like regions predicted. B) Alpha-fold prediction of the whole protein structure. C) Congo-red binding analysis.

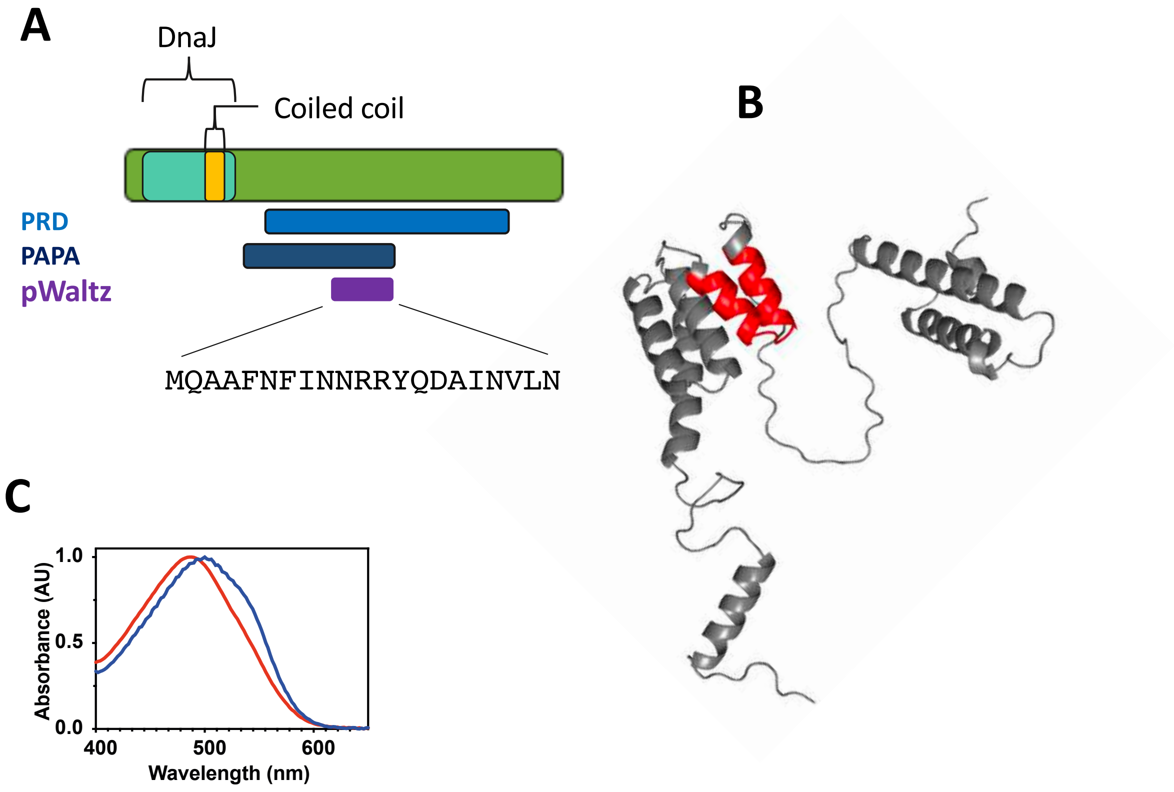


**Supplementary Figure 4. Predictive and experimental results associated to C0B555 containing C3.** A) Diagram sowing the prion-like regions predicted. B) Alpha-fold prediction of the whole protein structure. C) Congo-red binding analysis.

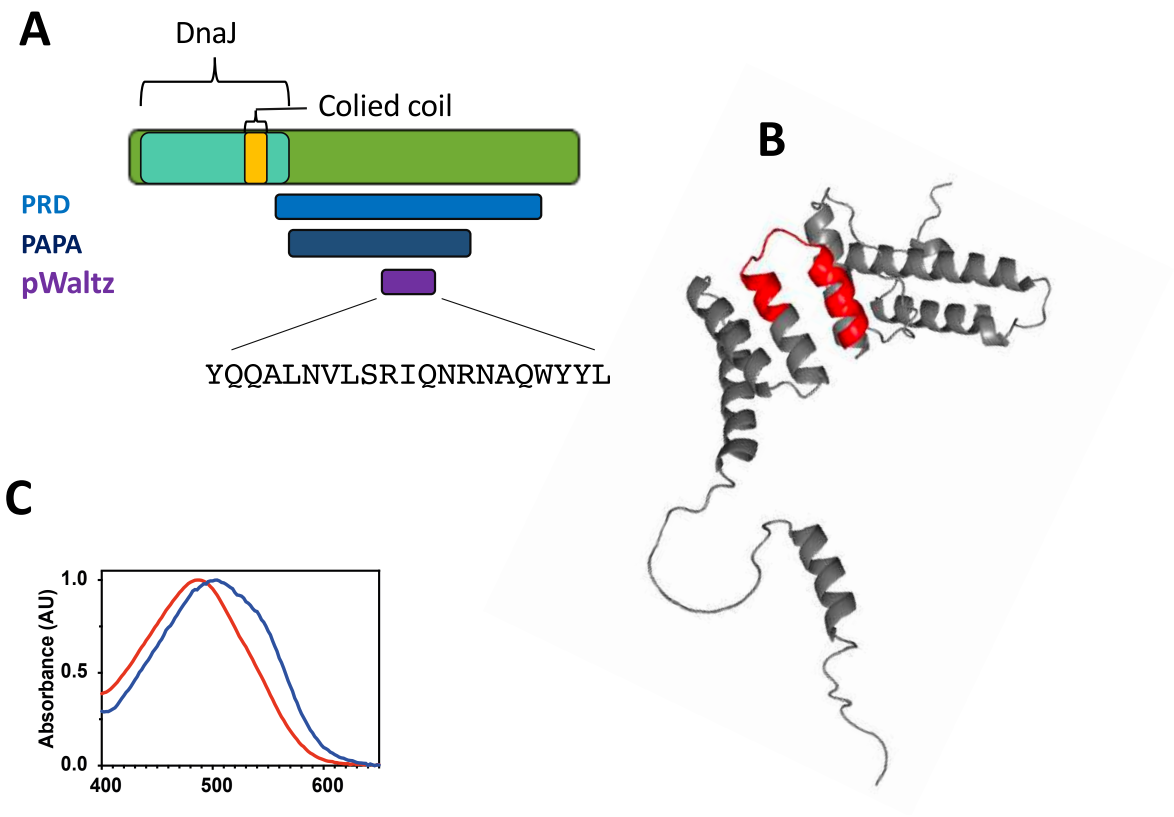


**Supplementary Figure 5. Predictive and experimental results associated to C9L6N5 containing C4.** A) Diagram sowing the prion-like regions predicted. B) Alpha-fold prediction of the whole protein structure. C) Congo-red binding analysis.

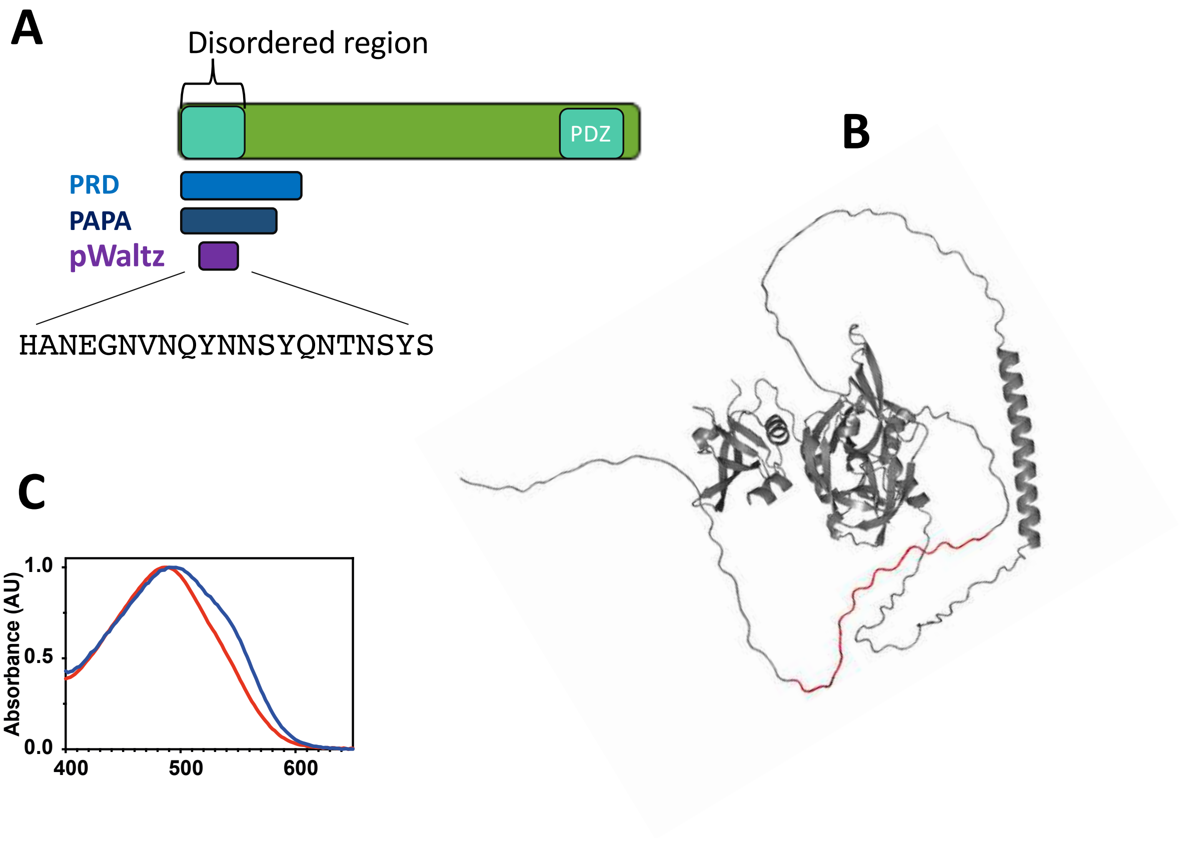


**Supplementary Figure 6. Predictive and experimental results associated to D4KZ46 containing C5.** A) Diagram sowing the prion-like regions predicted. B) Alpha-fold prediction of the whole protein structure. C) Congo-red binding analysis.

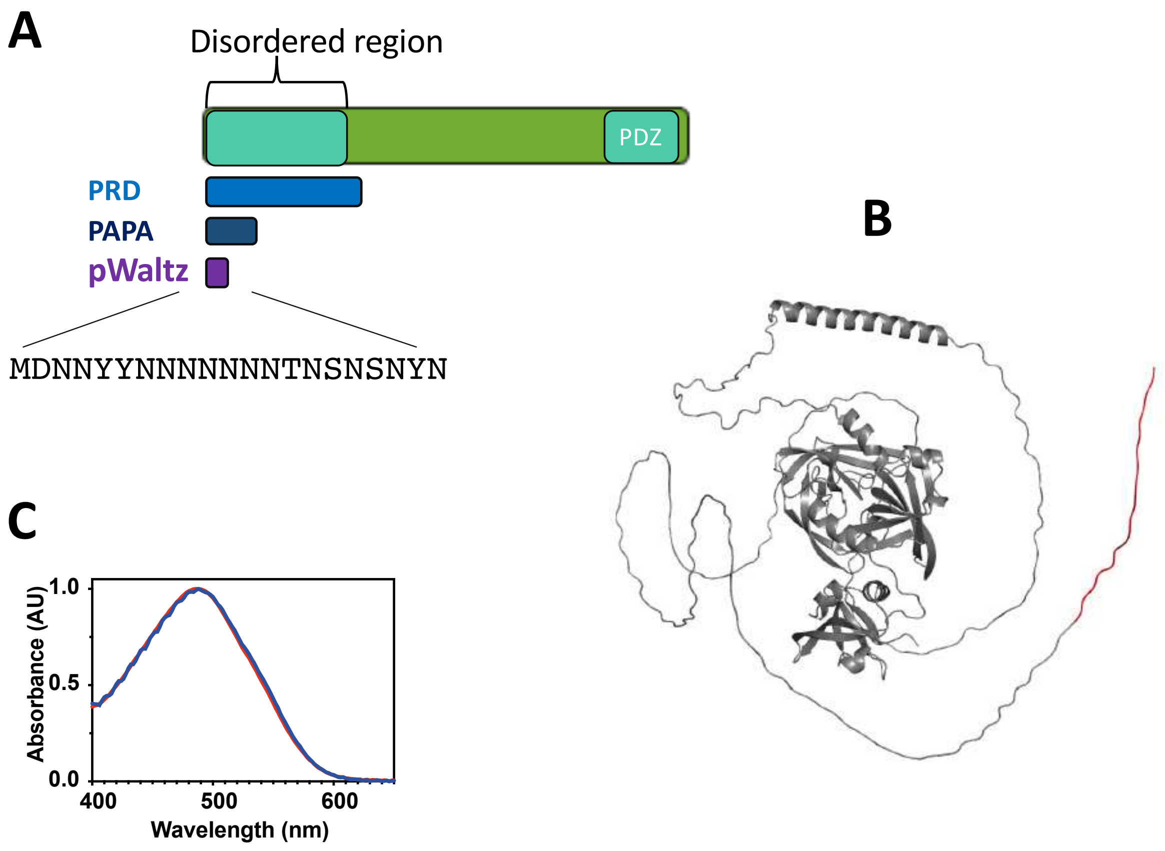


**Supplementary Figure 7. Predictive and experimental results associated to C0FUS6 containing C6.** A) Diagram sowing the prion-like regions predicted. B) Alpha-fold prediction of the whole protein structure. C) Congo-red binding analysis.

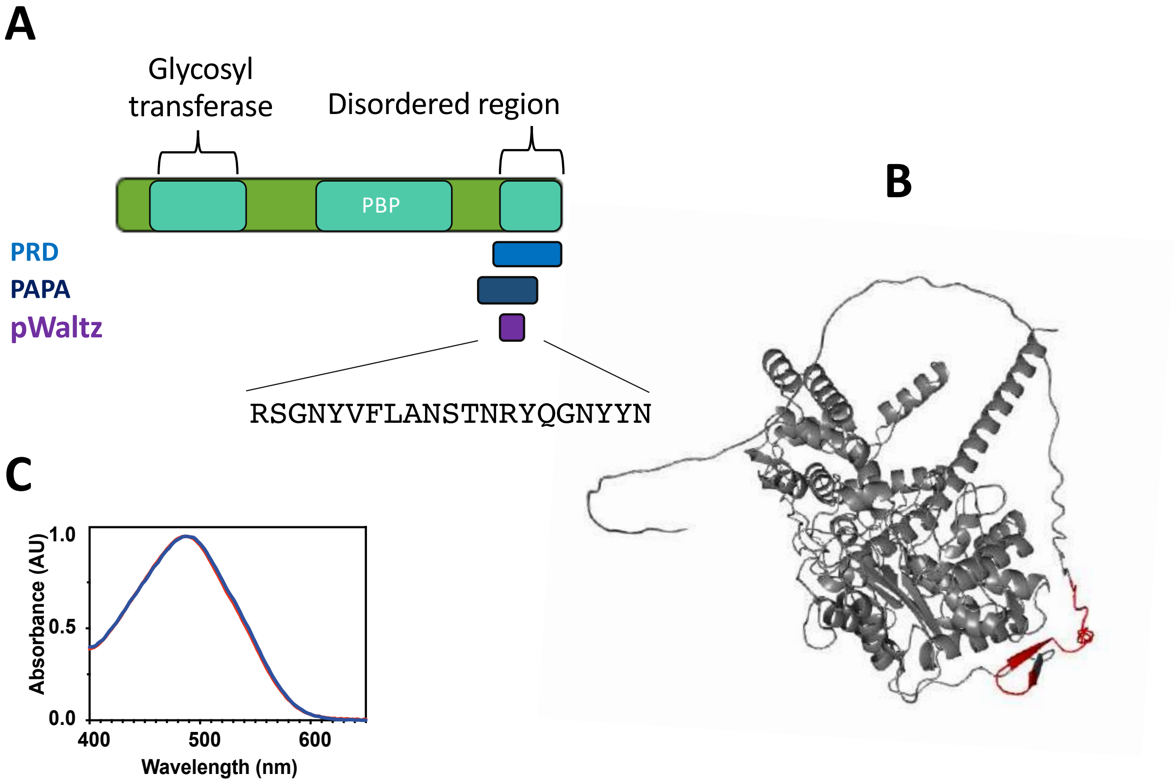


**Supplementary Figure 8. Predictive and experimental results associated to E7GXS5 (currently A0A448AIK9) containing C7.** A) Diagram sowing the prion-like regions predicted (PBP = penicillin-binding protein). B) Alpha-fold prediction of the whole protein structure. C) Congo-red binding analysis.

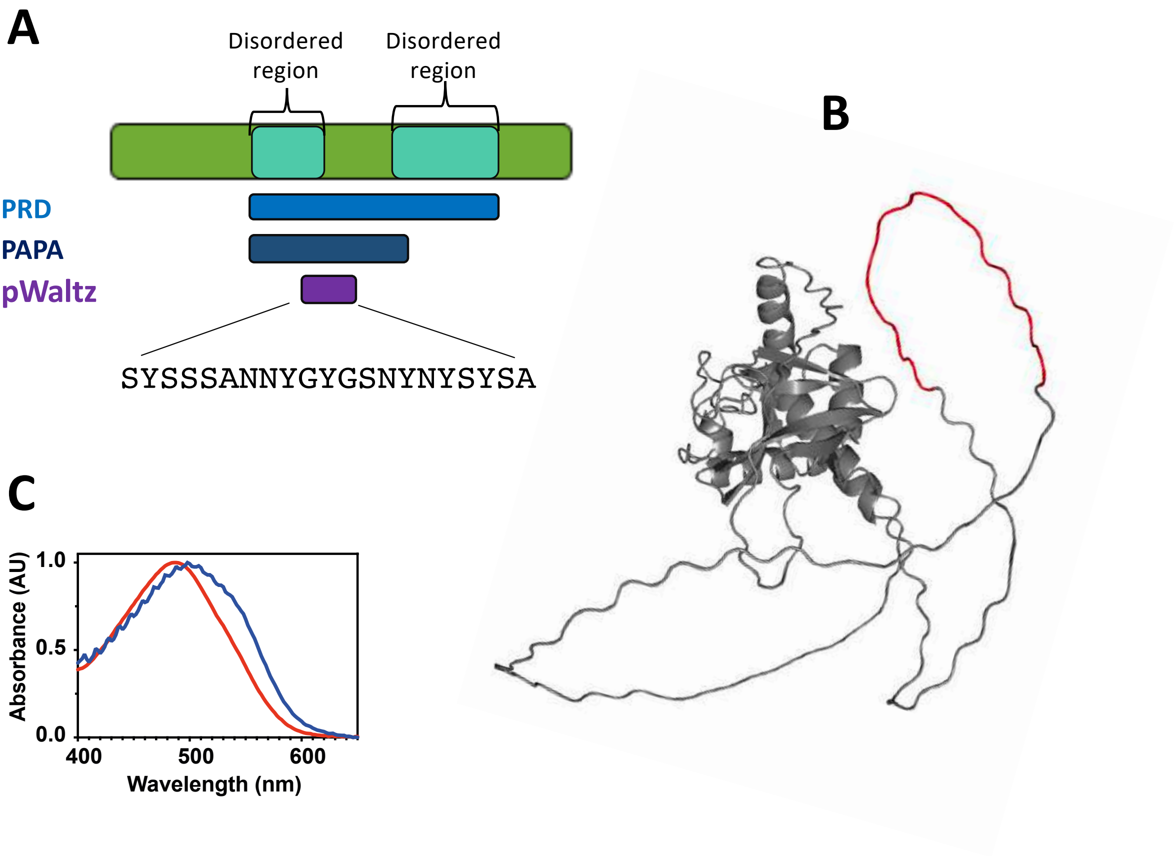


**Supplementary Figure 9. Predictive and experimental results associated to F0HUU2 containing C8.** A) Diagram sowing the prion-like regions predicted. B) Alpha-fold prediction of the whole protein structure. C) Congo-red binding analysis.

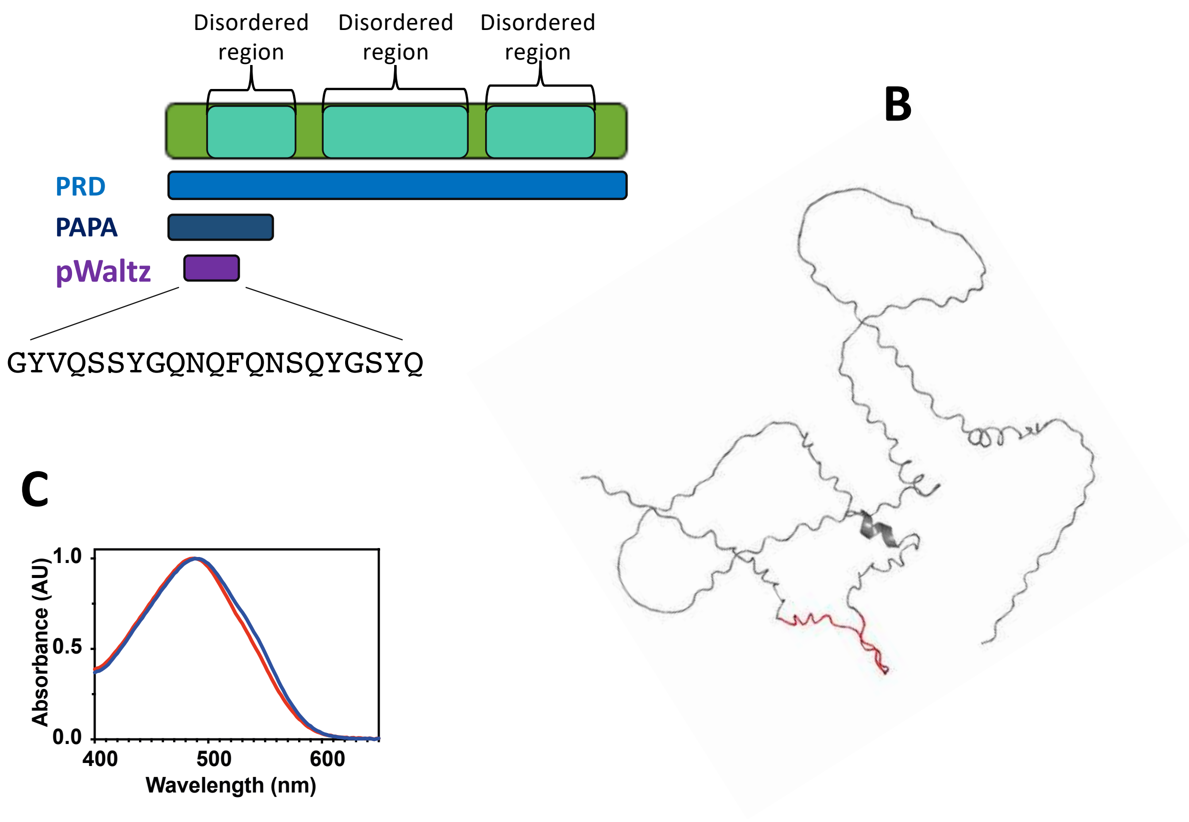


**Supplementary Figure 10. Predictive and experimental results associated to F5LFB6 containing C9.** A) Diagram sowing the prion-like regions predicted. B) Alpha-fold prediction of the whole protein structure. C) Congo-red binding analysis.

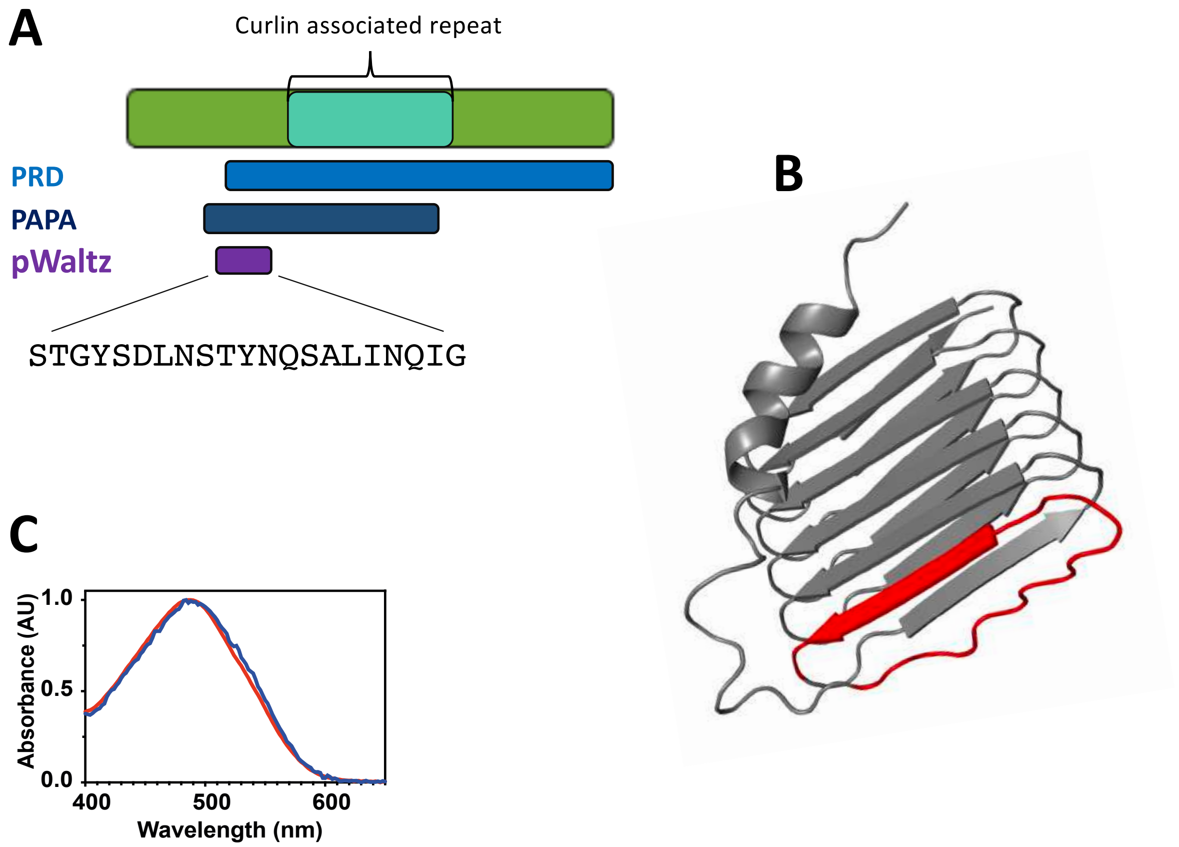


**Supplementary Figure 11. Predictive and experimental results associated to G9Y7N7 containing C10.** A) Diagram sowing the prion-like regions predicted. B) Alpha-fold prediction of the whole protein structure. C) Congo-red binding analysis.

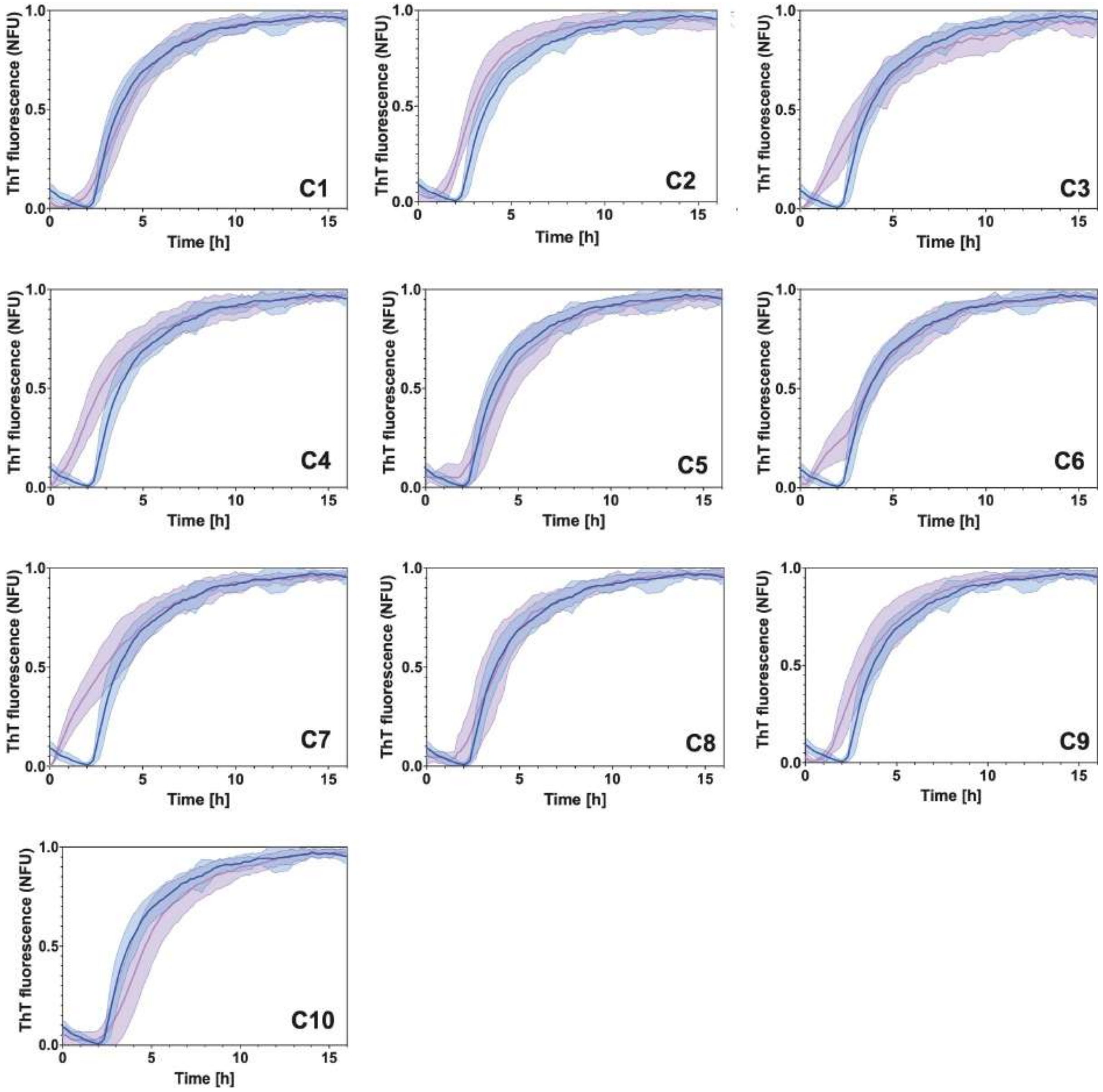


**Supplementary Figure 12. Aggregation kinetics plots of Aβ40 seeded with ten peptides derived from the gut microbiome.** The purple curve represents the aggregation kinetics of Aβ40 without any seeding, whereas the blue curve represents the aggregation kinetics in the presence of pre-aggregated peptides. The shaded regions show the standard error of the mean, which has been calculated from four independent replicates, with each replicate comprising three repeated samples.

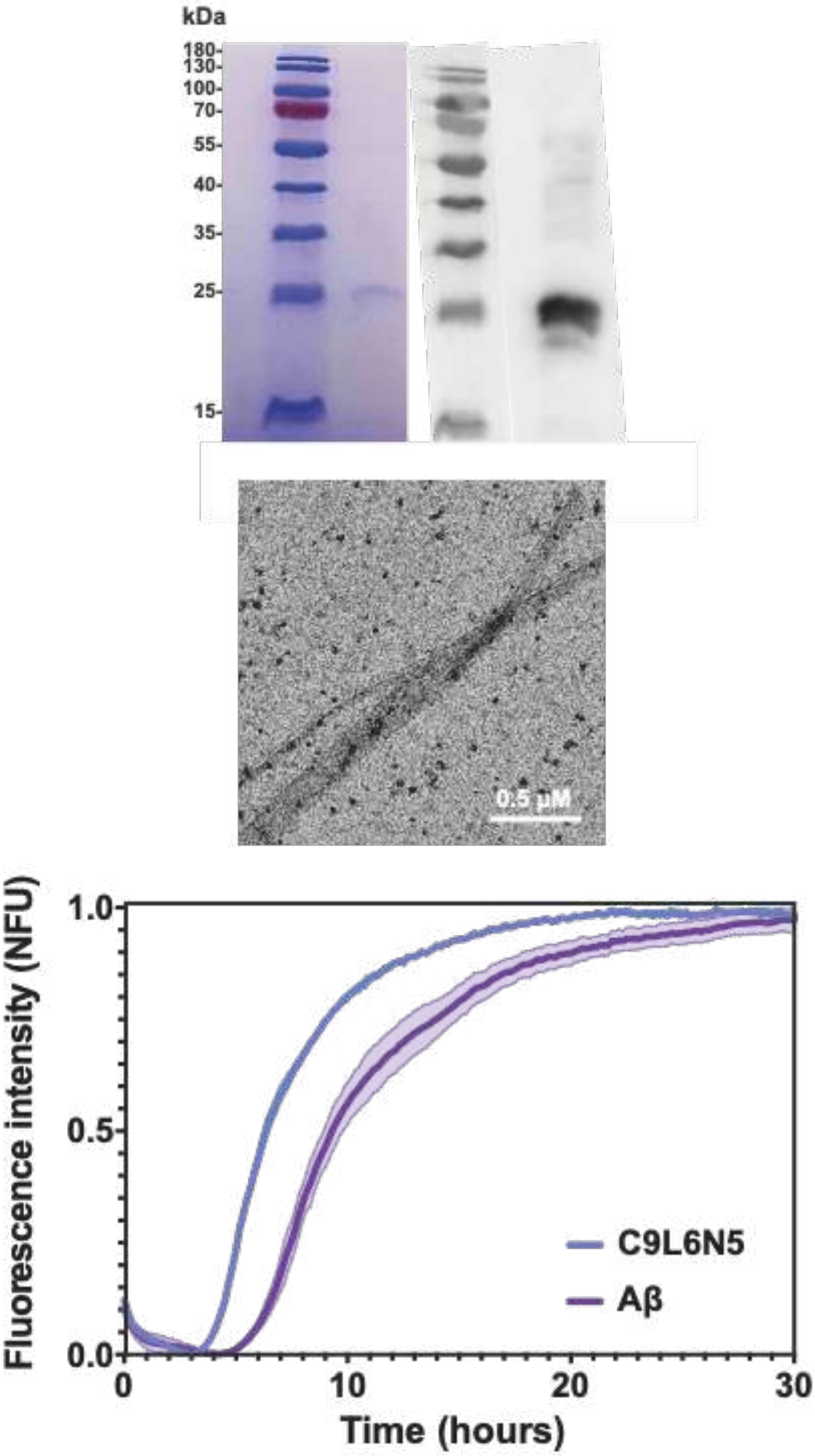


**Supplementary Figure 13. Purification and aggregation of C9L6N5 (containing C4).** Up, SDS-PAGE gel and western blotting (anti-His) showing one protein band after C9L6N5 purification. Middle, TEM image showing fibrillar aggregates of C9L6N5. Bottom, Aβ40 aggregation kinetics not seeded (purple) and seeded (blue) with aggregates of C9L6N5.

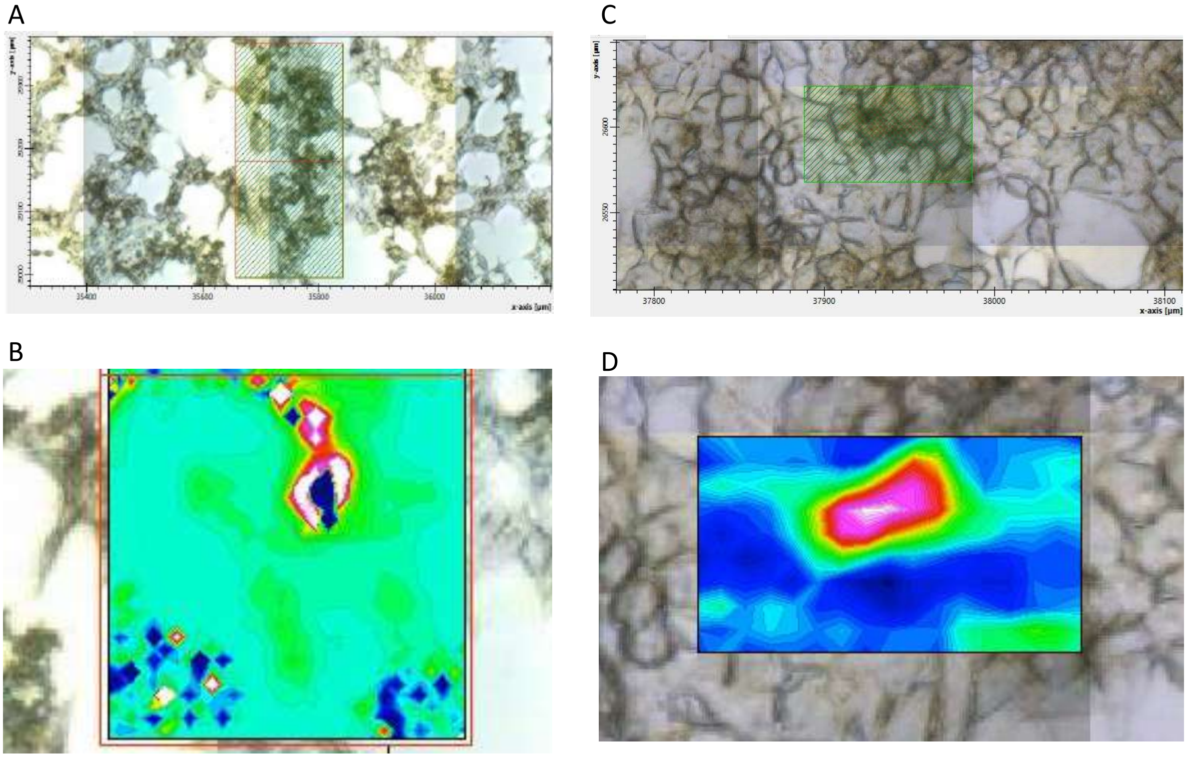


**Supplementary Figure 14. µFTIR maps of cells treated with aggregated peptides derived from the microbiota.** The upper panels show microscope images of cells incubated with the peptides (A) C2 and (C) C4. The lower panels illustrate the beta/alpha ratio (1740/2921 cm-1), utilized for detecting the presence of amyloid fibrils formed by the peptides (B) C2 and (D) C4. The images indicate the distribution of preformed aggregates throughout the samples. Amyloid deposits were identified in all samples, confirming the persistence of the preaggregated peptide's under cell culture conditions.

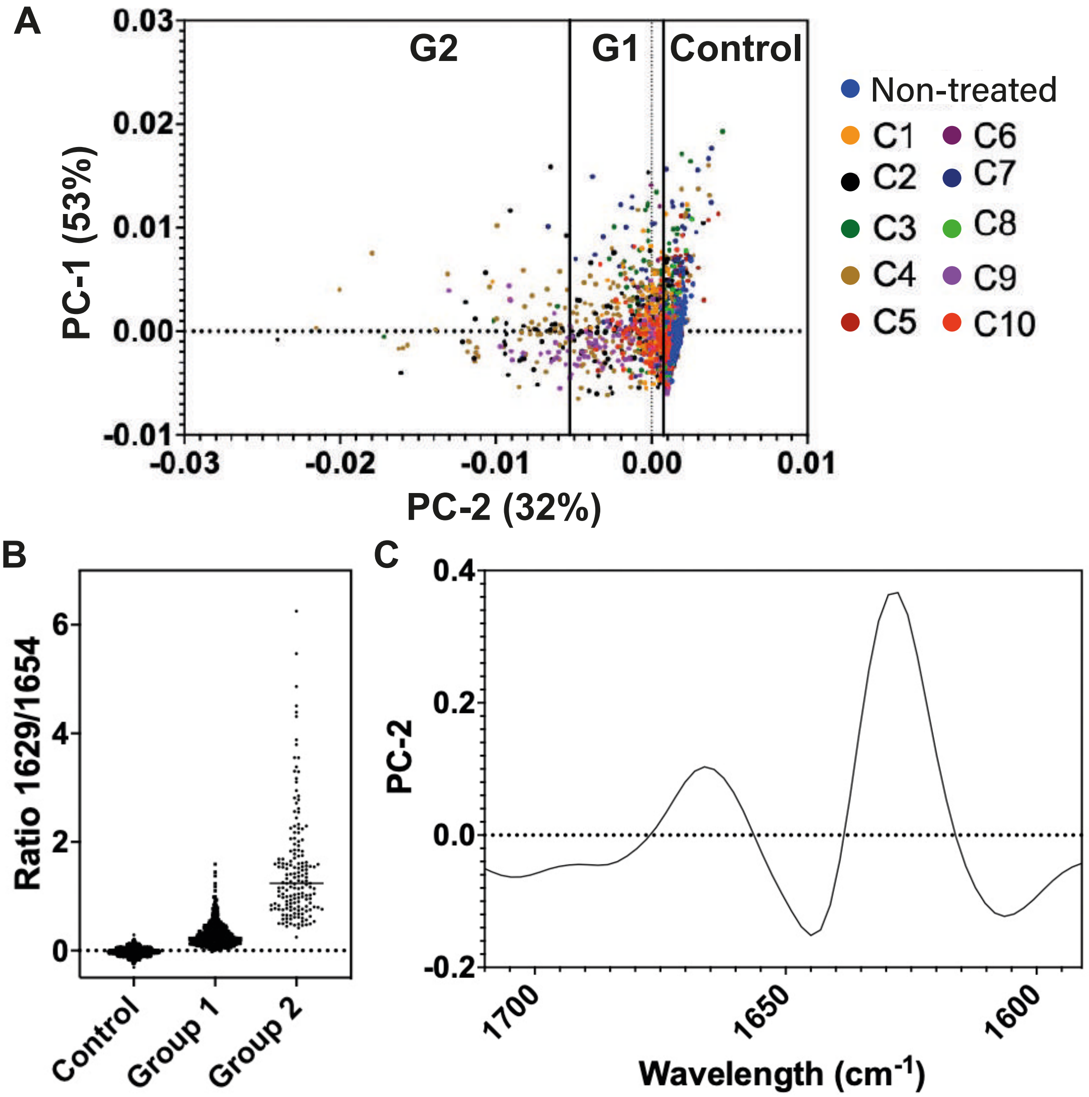


**Supplementary Figure 15. Results of μFTIR analysis showing that the absorbance of β-sheet differentiates between cells incubated with and without aggregates.** A) Principal component analysis of measurements in the amide I region, depicting the separation of three groups based on aggregate signal intensity. B) Distribution of the amyloid aggregation ratio (1629/1654) for the three groups derived from PCA analysis. The first group (Control, N=1181) corresponds to measurements taken in locations without amyloid aggregates, while group 1 (G1, N=465) and group 2 (G2, N=169) denote groups with progressively increasing aggregate signals. C) Summary μFTIR spectra of PC-2, representing 32% of the variances between the samples. The peak from 1650 cm-1 to 1600 cm-1 corresponds to β-sheet signal, indicating that the difference between the groups is linked to amyloid fibrils. Amyloid deposits (G1 and G2) were identified in all samples, confirming the persistence of aggregated peptides under cell culture conditions.

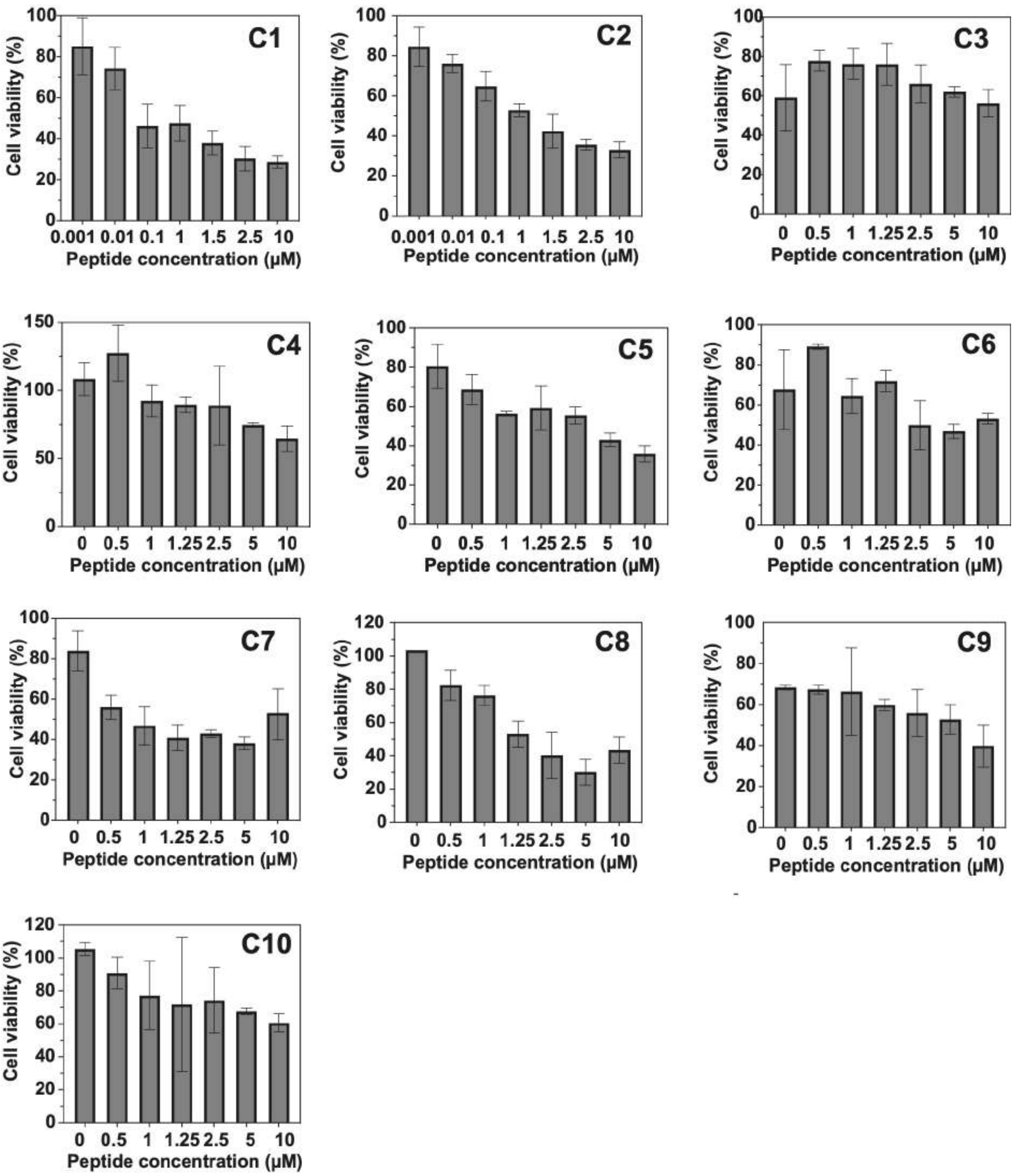


**Supplementary Figure 16. Cytotoxicity of amyloid-forming cores derived from the gut microbiome.** Cell viability of SH-SY5Y cells after 24 hours of exposure to various peptides (N=3). Due to their high toxicity, C1 and C2 have a wider concentration range. The error bars represent the standard error of the mean (SEM).

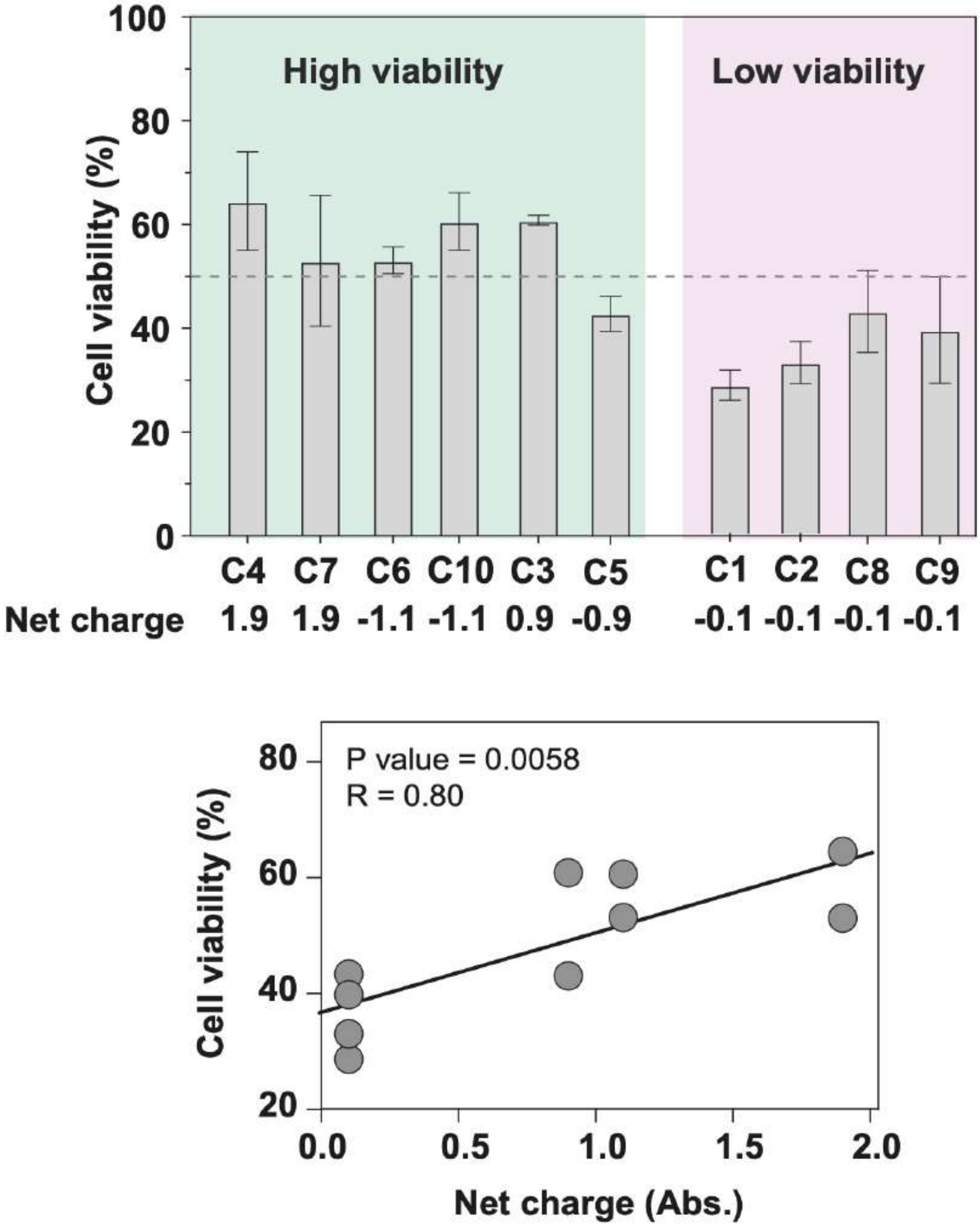


**Supplementary Figure 17. Comparison between the peptides charge and their cytotoxicity.** More charge (in absolute number, Abs.) is associated to more cell viability.

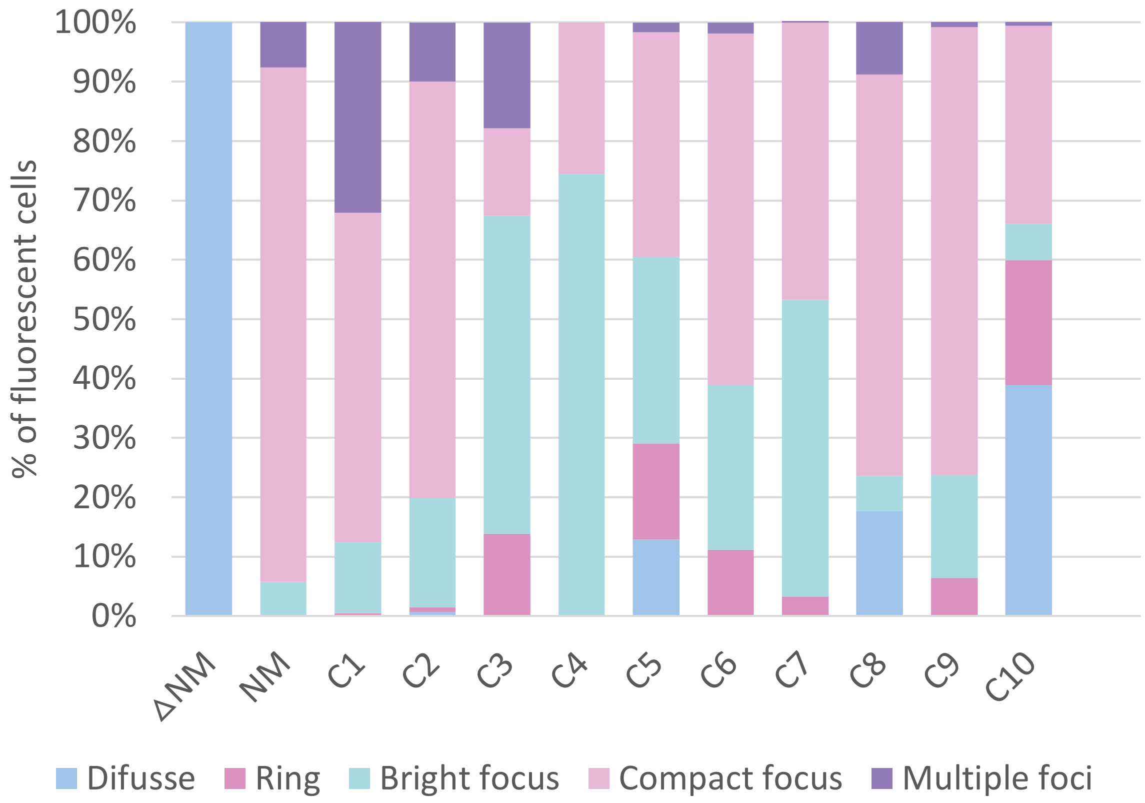


**Supplementary Figure 18. Location of the different Sup35p chimeras with amyloid-forming cores derived from the gut microbiome.** Percentage count of the various shapes detected by fluorescent microscopy, which the Sup35p chimeras adopted within the *S. cerevisiae*.

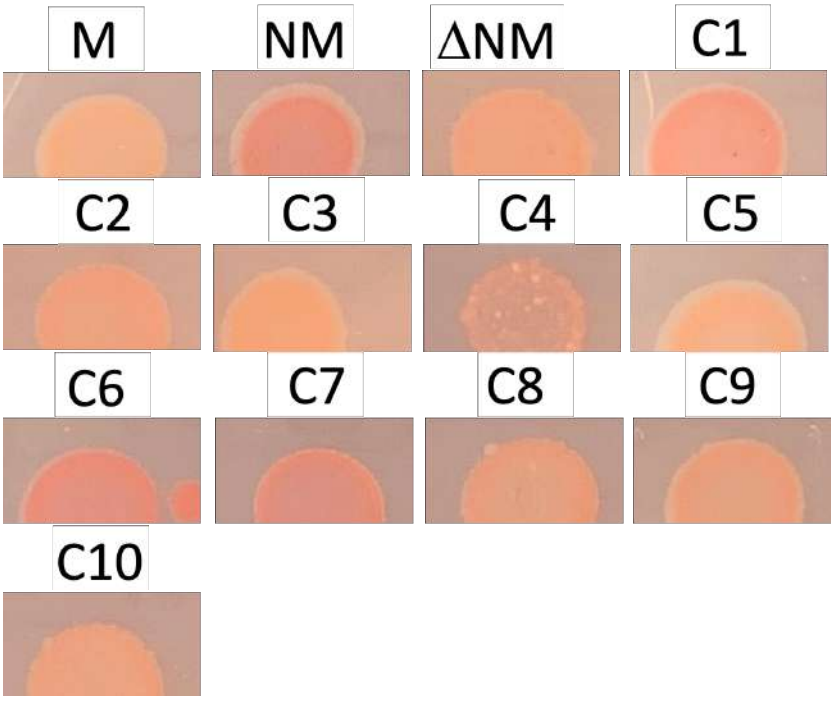


**Supplementary Figure 19. E. coli C-DAG colonies on Congo Red-containing plates.** The images show the colonies of the different *E. coli* expressing the Sup35 chimeras grown on congo-red containing plates (Methods).

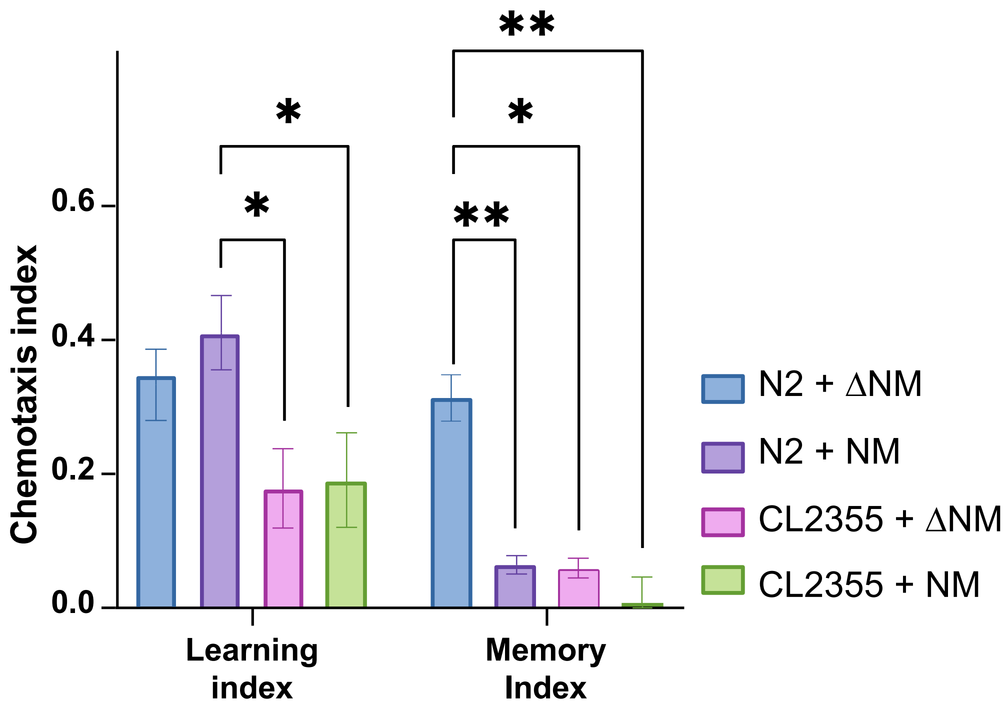


**Supplementary Figure 20. STAM comparison between N2 (wildtype) and CL2355 (Alzheimer's model).** Chemotaxis index for learning and memory assays measured for worms fed with ΔSup35 and Sup35NM (One-way ANOVA, N=3).

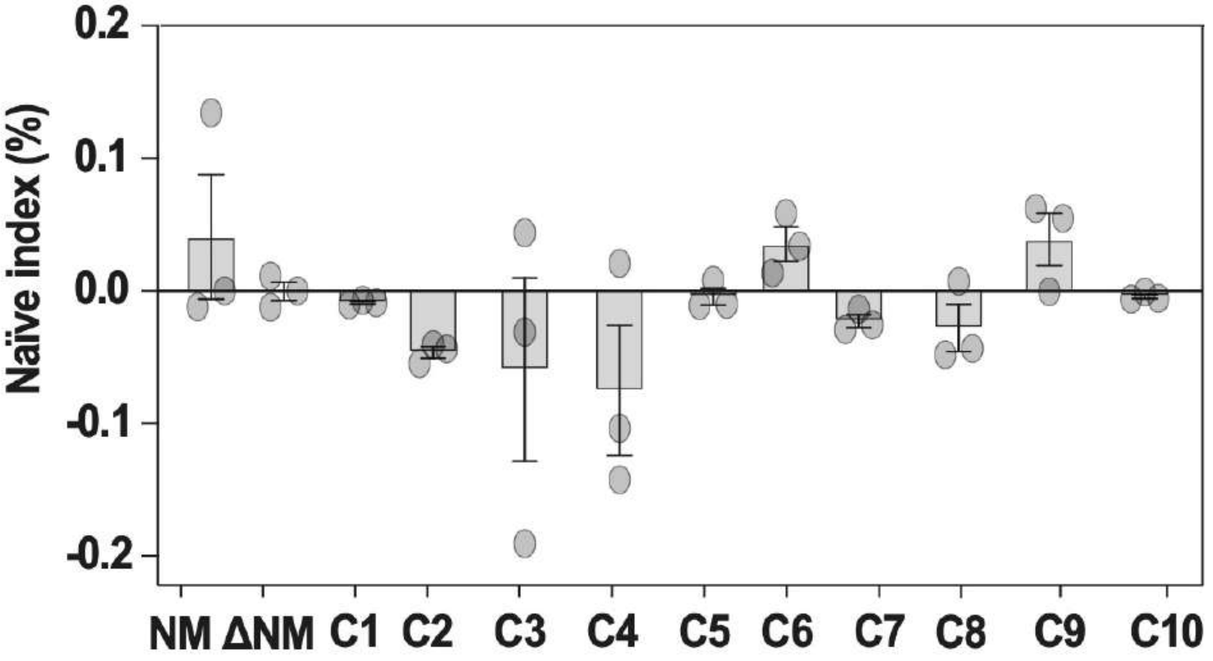


**Supplementary Figure 21. Chemotaxis index of naïve worms**. Chemotaxis index measured for unconditioned worms. (One-way ANOVA, N=3).

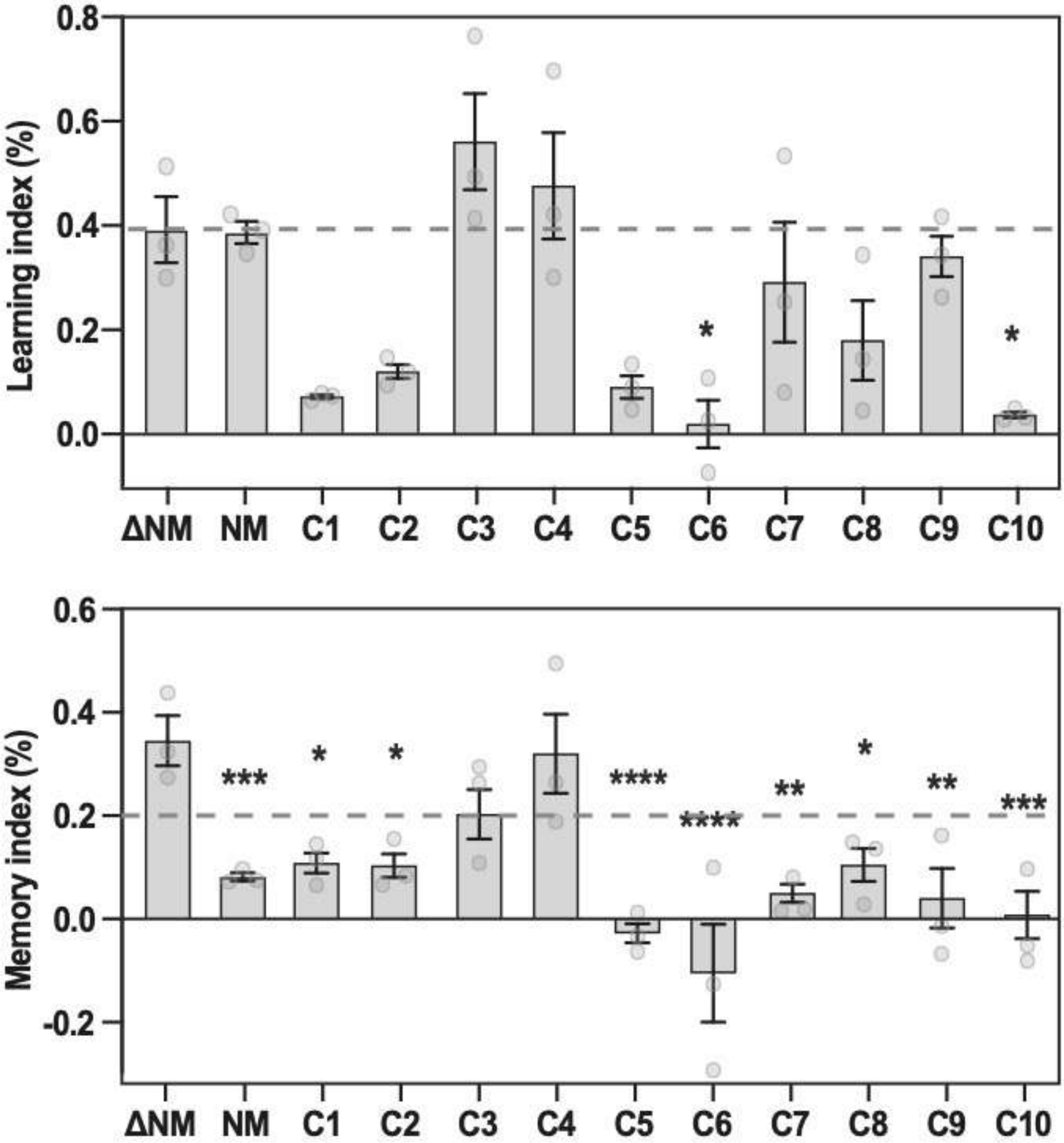


**Supplementary Figure 22. Chemotaxis index measured 1 hour after conditioning.** Top learning assay, bottom memory assay (one-way ANOVA, N=3).

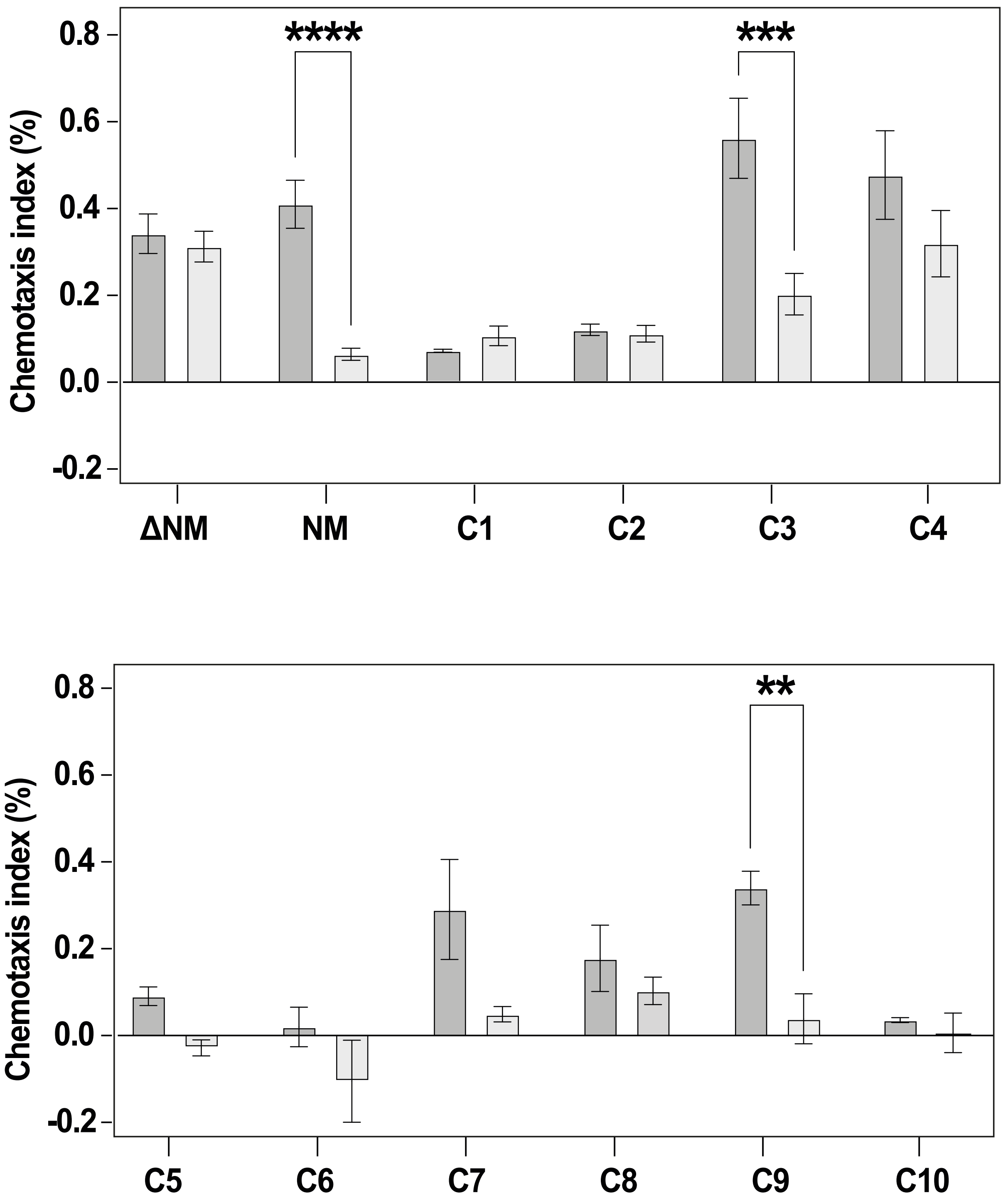


**Supplementary Figure 23. Comparison between cognitive effects.** Comparison of the learning index (in dark grey) and the memory index (in light grey).

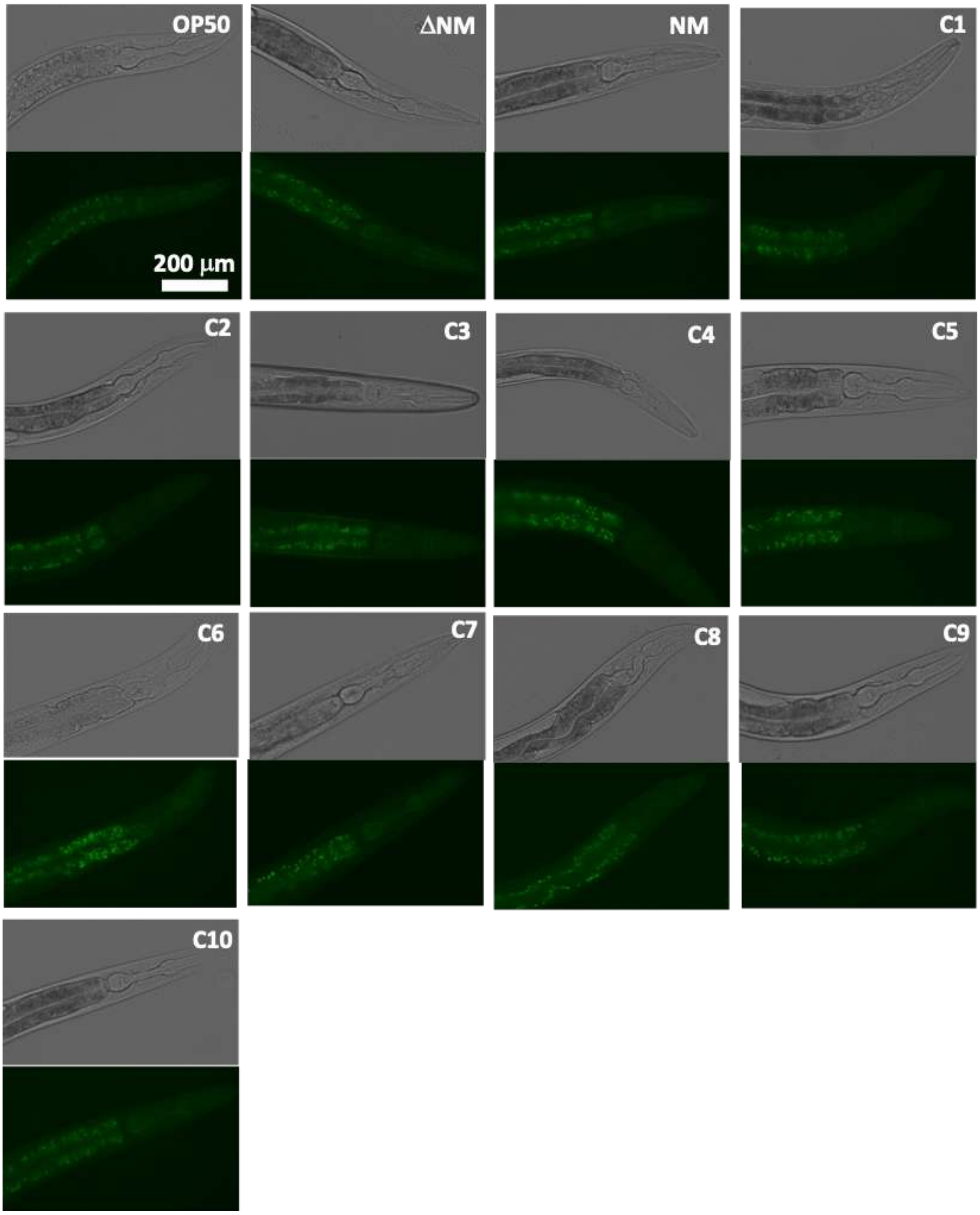


**Supplementary Figure 24. The intake of bacterial aggregates affects the gut granules autofluorescence.** Images showing the fluorescence and transmitted-light images of the frontal section of *C. elegans* fed with *E. coli* expressing different Sup35NM chimeras. The images are centered on the first two intestine rings, carefully avoiding the presence of the gonads to mitigate the interference.

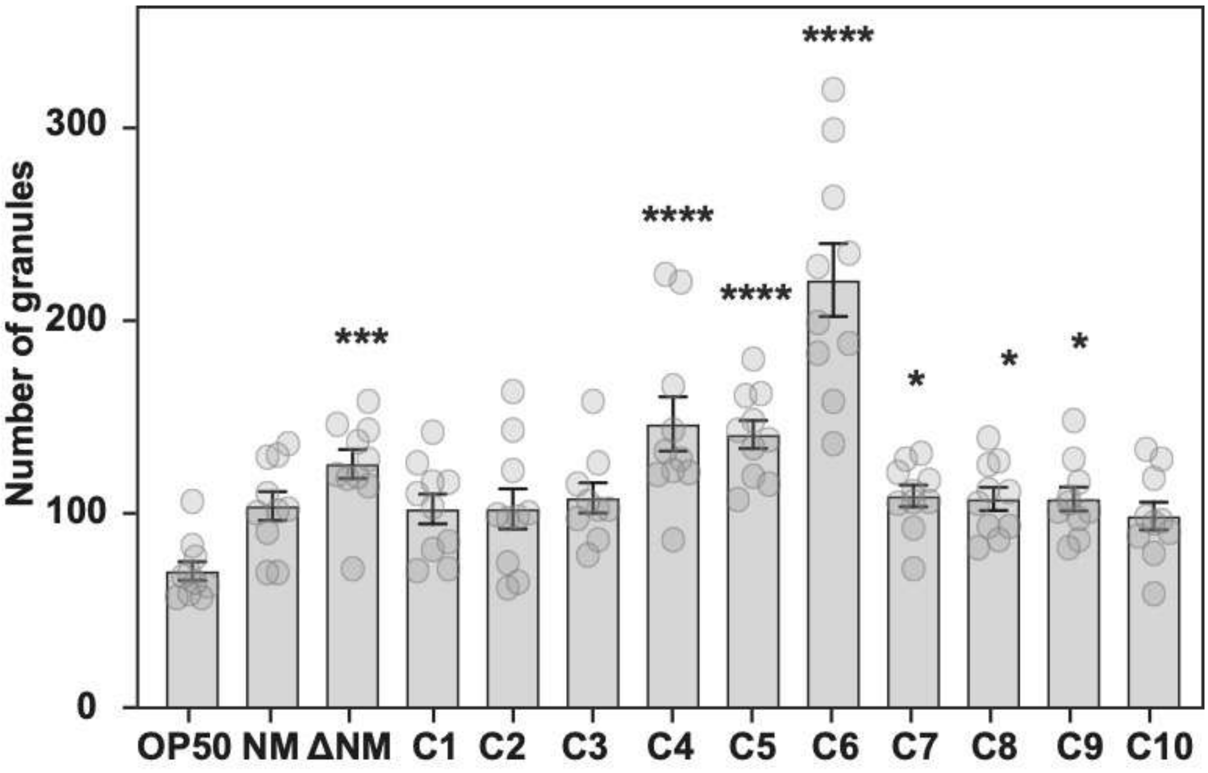


**Supplementary Figure 25.** ***C. elegans*** **gut granules counting.** Number distribution of gut granules in *C. elegans* fed *E. coli* expressing various Sup35NM variants. Error bars represent the SEM of 8-10 measurements. Statistical significance was determined using one-way ANOVA (*p < 0.05, **p < 0.01, ***p < 0.001, ****p < 0.0001).

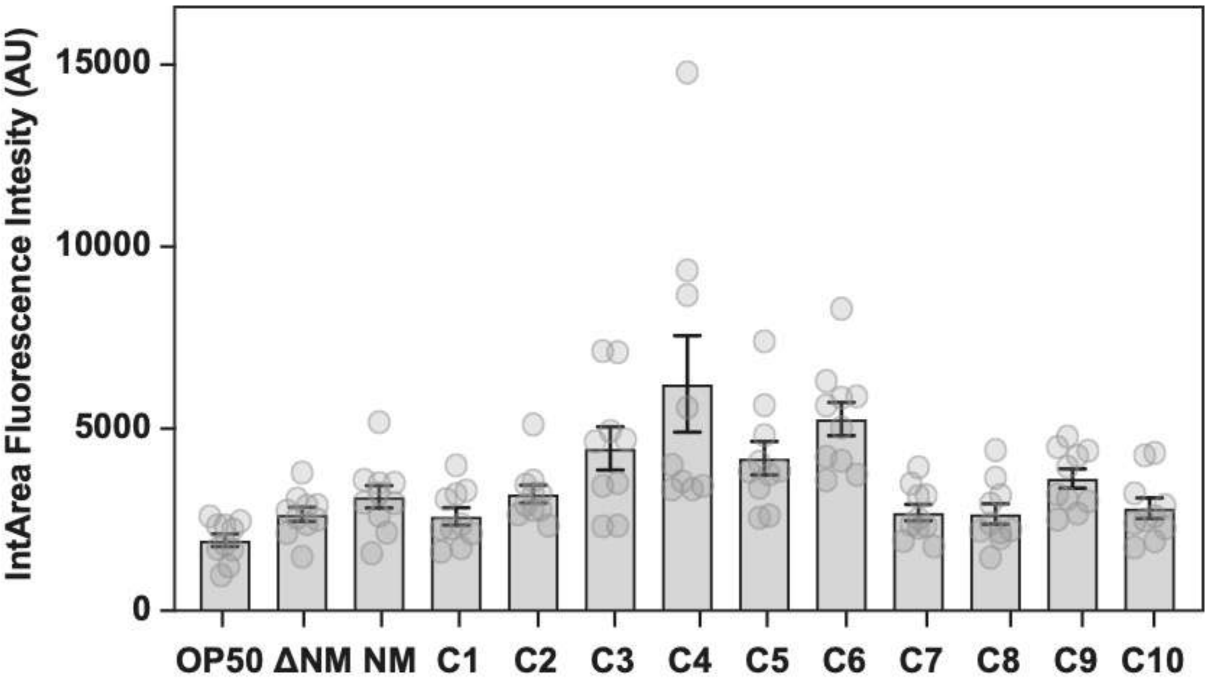


**Supplementary Figure 26. *C. elegans* gut granules fluorescent intensity.** Plot showing the distribution of the gut granules fluorescent intensity adjusted by area (IntArea, ImageJ) in the *C. elegans* fed with *E. coli* expressing different variants of Sup35NM. Error bars represent the SEM of 8-10 measurements.

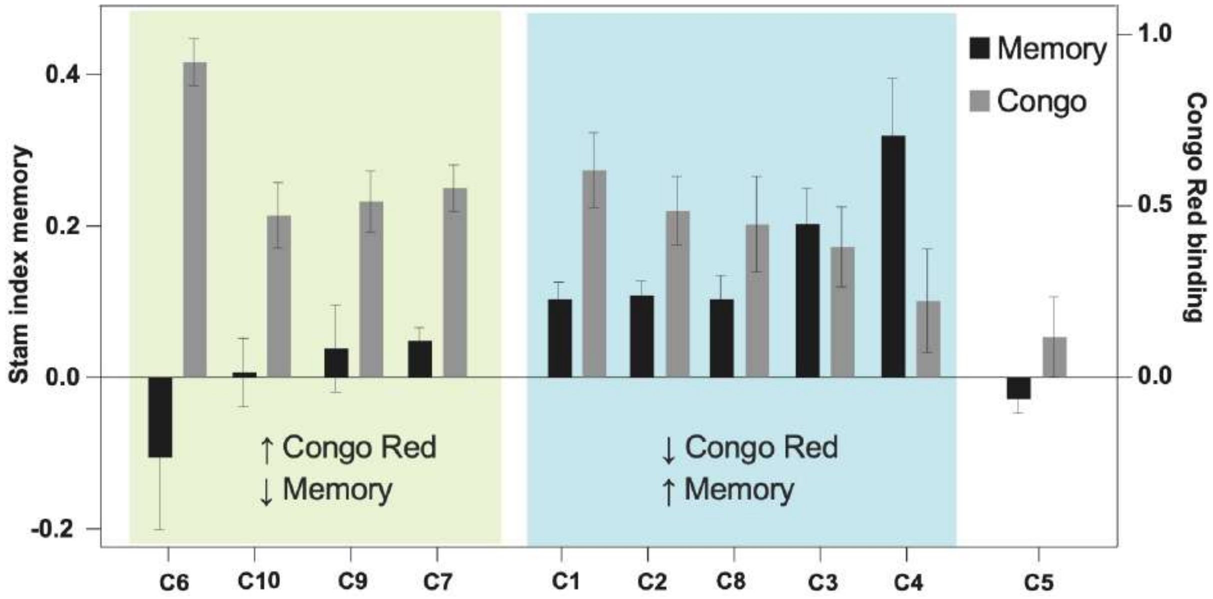


**Supplementary Figure 27. Memory index and Congo Red binding comparison.** Plot comparing the memory index and the congo red binding (from Figure 4B) of *E. coli* expressing different variants of Sup35NM.

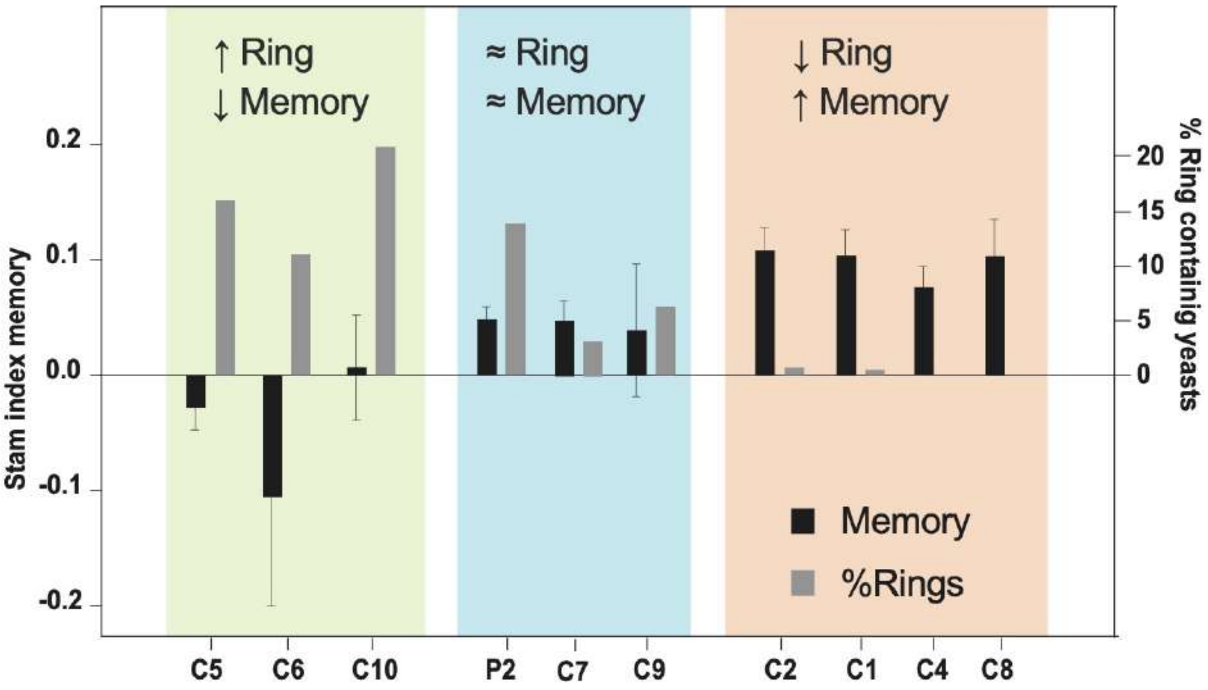


**Supplementary Figure 28. Memory index and presence of ring aggregates comparison.** Plot comparing the memory index and the presence of ring structures in yeast (from Figure 3B) expressing different variants of Sup35NM.

| Name | Gene name | Uniprot | pW | Amyloid core | Protein description | Species | GRAM | PSORT |
|---|---|---|---|---|---|---|---|---|
| **C1** | HMPREF1429_00551 | M3SJI9 | 75,856 | GNNNNSVISFNQTNFNQGTYN | putative vacuolating cytotoxin | Helicobacter pylori GAMchJs124i | - | Extracellular/Outer Membrane |
| **C2** | HMPREF1434_01202 | M5YJZ4 | 78,866 | SSVYWLNSVNENNNNKSYYIS | putative vacuolating cytotoxin | Helicobacter pylori GAM93Bi | - | Extracellular/Outer Membrane |
| **C3** | COPCOM_00275 | C0B555 | 78,1175 | MQAAFNFINNRRYQDAINVLN | DnaJ domain protein | Coprococcus comes ATCC 27758 | + | Cytoplasmic/Extracellular |
| **C4** | BLAHAN_05046 | C9L6N5 | 75,1871 | YQQALNVLSRIQNRNAQWYYL | DnaJ domain protein | Blautia hansenii DSM 20583 | + | Extracellular |
| **C5** | RO1_21080 | D4KZ46 | 77,2415 | HANEGNVNQYNNSYQNTNSYS | trypsin-like serine proteases | Roseburia intestinalis XB6B4 | + | Unknown** |
| **C6** | ROSEINA2194_02497 | C0FUS6 | 77,1301 | MDNNYYNNNNNNNTNSNSNYN | trypsin | Roseburia inulinivorans DSM 16841 | + | Extracellular |
| **C7** | HMPREF9459_01032 | E7GXS5* | 75,7126 | RSGNYVFLANSTNRYQGNYYN | penicillin-binding protein 1A | Streptococcus anginosus 1_2_62CV | + | Extracellular |
| **C8** | HMPREF5505_0689 | F0HUU2 | 74,5819 | SYSSSANNYGYGSNYNYSYSA | aggregation promoting factor | Lactobacillus delbrueckii subsp. lactis DSM 20072 | + | Extracellular |
| **C9** | HMPREF9413_3202 | F5LFB6 | 73,8652 | GYVQSSYGQNQFQNSQYGSYQ | putative RNA-binding protein FUS | Paenibacillus sp. HGF7 | +/- | Extracellular |
| **C10** | HMPREF0454_02561 | G9Y7N7 | 73,7219 | STGYSDLNSTYNQSALINQIG | curlin associated repeat | Hafnia alvei ATCC 51873 | - | Extracellular |

**Supplementary Data 1. Table showing ten amyloid-forming cores selected from gut microbiome.** pW indicates the pWALTZ score. *It is an obsolete entry in UniProt. The correct actual sequence is "A0A448AIK9," which includes the change I63V with respect to "E7GXS5". **Single-pass membrane according to UniProt.

**Supplementary Data 2. Predictions performed on the Sup35p variants studied in this work.**

Presented here are the sequences of the Sup35NM chimeras, with the newly detected sequences highlighted by pWALTZ (Material and Methods). Sequences in yellow indicate a precise match with the introduced amyloid-forming cores. Sequences in blue signify a partial match or mismatch with the introduced amyloid-forming cores, including the regions in Sup35NM and Sup35M highlighted in blue. The Sup35p nucleation region is denoted in bold and underlined. The introduced amyloid-forming cores are marked in bold.

List of pWALTZ detected sequence on the region detected by PAPA.

```
FULL  RGNYKNFNYNNNLQGYQAGFQ 73.9926 (residues from 98 to 118)
Delta RGNYKNFNYNNNLQGYQAGFQ 73.9926 (residues from 58 to 68)
C1    GNNNNSVISFNQTNFNQGTYN 78.8660 (residues from 2 to 22)
C2    NNNKSYYISAGGYYQNYQGYS 77.6716 (residues from 14 to 34)
C3    MQAAFNFINNRRYQDAINVLN 78.1175 (residues from 2 to 22)
C4    RNAQWYYLAGGYYQNYQGYSG 78.2290 (residues from 15 to 35)
C5    YNNSYQNTNSYSAGGYYQNYQ 80.0127 (residues from 11 to 31)
C6    MDNNYYNNNNNNNTNSNSNYN 77.1301 (residues from 2 to 22)
C7    RSGNYVFLANSTNRYQGNYYN 75.7126 (residues from 2 to 22)
C9    NQFQNSQYGSYQAGGYYQNYQ 77.5441 (residues from 11 to 31)
C8    GYGSNYNYSYSAAGGYYQNYQ 78.4679 (residues from 11 to 31)
C10   TYNQSALINQIGAGGYYQNYQ 77.2256 (residues from 11 to 31)
```

In **bold** the 21 peptides studied.
In yellow when they matches 100% with the new pWALTZ prediction.
In blue when they doesn't match 100% with the new pWALTZ prediction
In **<u>bold and underlined</u>** the Sup35p nucleation region.

>Sup35NM
M**<u>SDSNQGNNQQNYQQYSQNGNQQQGNNRYQGYQAYNAQAQP</u>**AGGYYQNYQGYSGYQQGGYQQ
YNPDAGYQQQYNPQGGYQQYNPQGGYQQQFNPQGGRGNYKNFNYNNNLQGYQAGFQPQSQGM
SLNDFQKQQKQAAPKPKKTLKLVSSSGIKLANATKKVGTKPAESDKKEEEKSAETKEPTKEP
TKVEEPVKKEEKPVQTEEKTEEKSELPKVEDLKISESTHNTNNANVTSADALIKEQEEEVDD
EVVNDMFGGKDHVSLIFMGHVDAGKSTMGGNLLYLTGSVDKRTIEKYEREAKDAGRQGWYLS
WVMDTNKEERNDGKTIEVGKAYFETEKRRYTILDAPGHKMYVSEMIGGASQADVGVLVISAR
KGEYETGFERGGQTREHALLAKTQGVNKMVVVVNKMDDPTVNWSKERYDQCVSNVSNFLRAI
GYNIKTDVVFMPVSGYSGANLKDHVDPKECPWYTGPTLLEYLDTMNHVDRHINAPFMLPIAA
KMKDLGTIVEGKIESGHIKKGQSTLLMPNKTAVEIQNIYNETENEVDMAMCGEQVKLRIKGV
EEEDISPGFVLTSPKNPIKSVTKFVAQIAIVELKSIIAAGFSCVMHVHTAIEEVHIVKLLHK
LEKGTNRKSKKPPAFAKKGMKVIAVLETEAPVCVETYQDYPQLGRFTLRDQGTTIAIGKIVK
IAE

PAPA

Score = 0.10, Position = 4

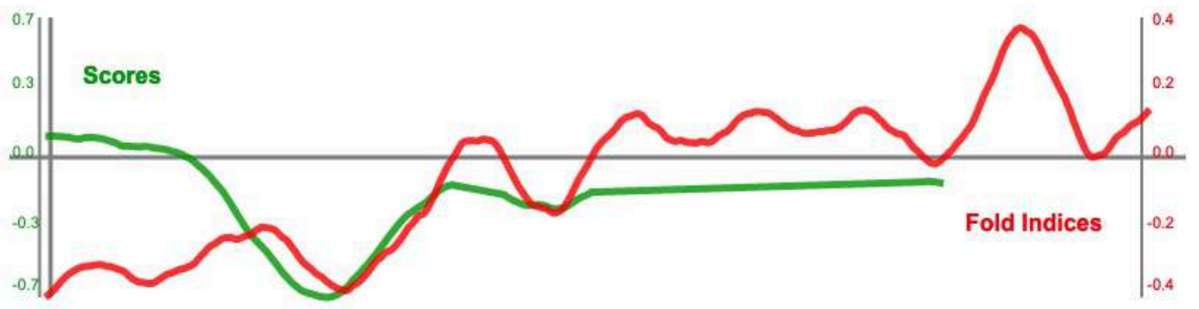


PLAAC

| COREscore | LLR | PAPAprop | PAPAfi |
|---|---|---|---|
| 23.306 | 23.306 | 0.100 | -0.423 |

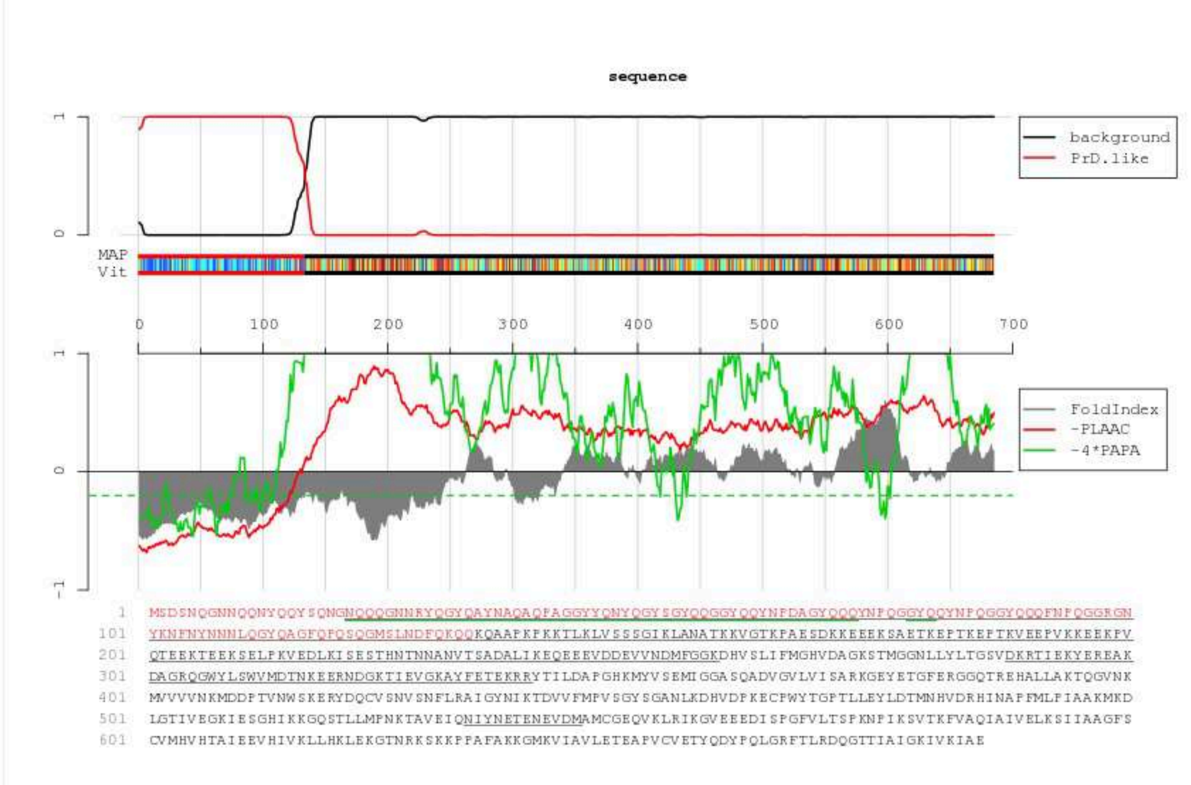

>Sup35M
MAGGYYQNYQGYSGYQQGGYQQYNPDAGYQQQYNPQGGYQQYNPQGGYQQQFNPQGGRGNYK
NFNYNNNLQGYQAGFQPQSQGMSLNDFQKQQKQAAPKPKKTLKLVSSSGIKLANATKKVGTK
PAESDKKEEEKSAETKEPTKEPTKVEEPVKKEEKPVQTEEKTEEKSELPKVEDLKISESTHN
TNNANVTSADALIKEQEEEVDDEVVNDMFGGKDHVSLIFMGHVDAGKSTMGGNLLYLTGSVD
KRTIEKYEREAKDAGRQGWYLSWVMDTNKEERNDGKTIEVGKAYFETEKRRYTILDAPGHKM
YVSEMIGGASQADVGVLVISARKGEYETGFERGGQTREHALLAKTQGVNKMVVVVNKMDDPT
VNWSKERYDQCVSNVSNFLRAIGYNIKTDVVFMPVSGYSGANLKDHVDPKECPWYTGPTLLE
YLDTMNHVDRHINAPFMLPIAAKMKDLGTIVEGKIESGHIKKGQSTLLMPNKTAVEIQNIYN
ETENEVDMAMCGEQVKLRIKGVEEEDISPGFVLTSPKNPIKSVTKFVAQIAIVELKSIIAAG
FSCVMHVHTAIEEVHIVKLLHKLEKGTNRKSKKPPAFAKKGMKVIAVLETEAPVCVETYQDY
PQLGRFTLRDQGTTIAIGKIVKIAE

PAPA

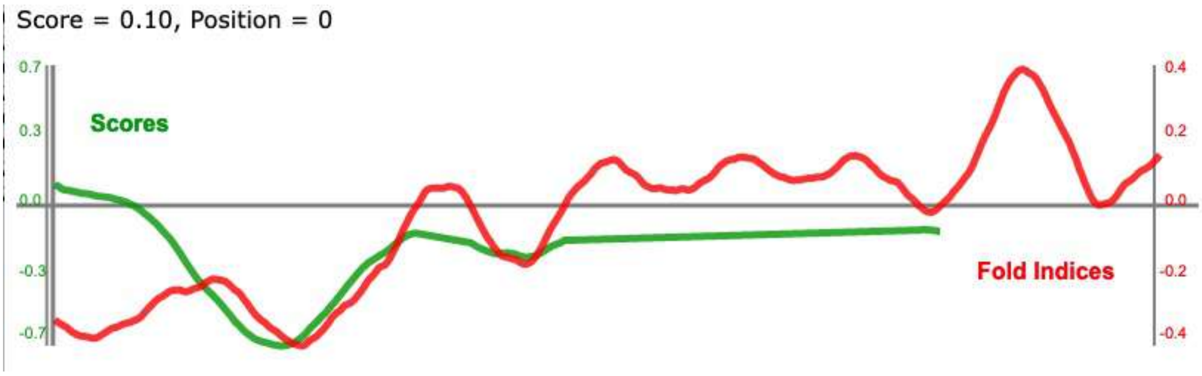


PLAAC

| COREscore | LLR | PAPAprop | PAPAfi |
|---|---|---|---|
| 19.025 | 19.025 | 0.099 | -0.342 |

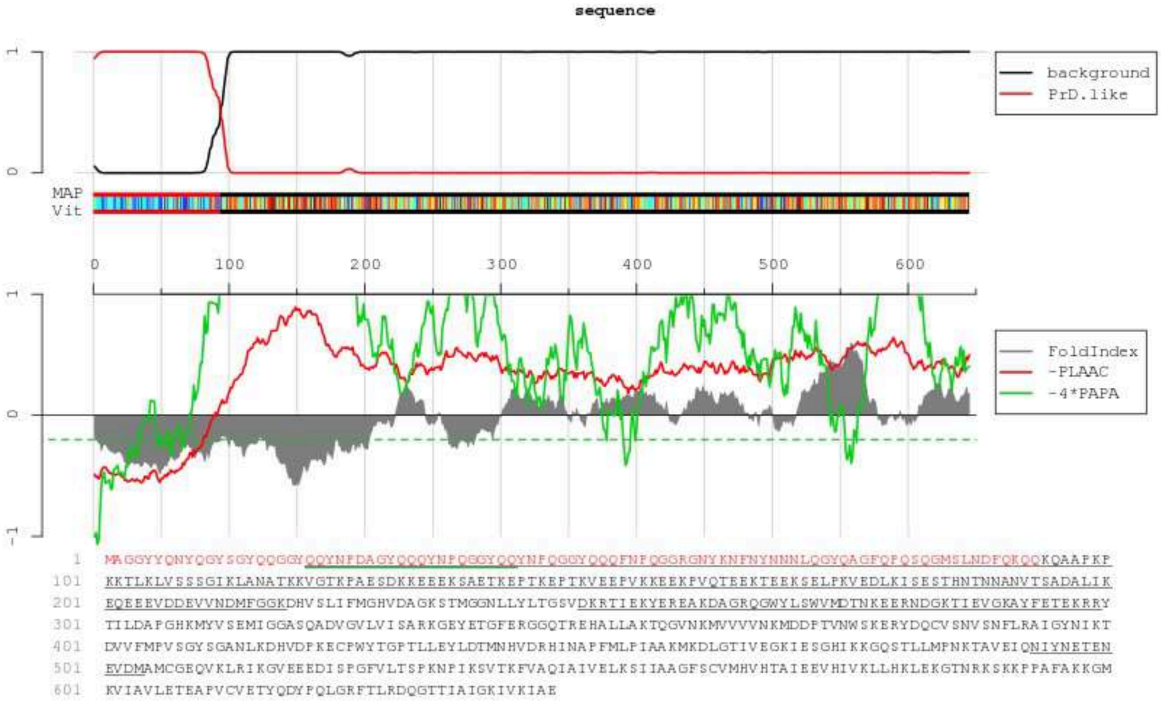

>C1

**MGNNNNSVISFNQTNFNQGTYNA**GGYYQNYQGYSGYQQGGYQQYNPDAGYQQQYNPQGGYQQ
YNPQGGYQQQFNPQGGRGNYKNFNYNNNLQGYQAGFQPQSQGMSLNDFQKQQKQAAPKPKKT
LKLVSSSGIKLANATKKVGTKPAESDKKEEEKSAETKEPTKEPTKVEEPVKKEEKPVQTEEK
TEEKSELPKVEDLKISESTHNTNNANVTSADALIKEQEEEVDDEVVNDMFGGKDHVSLIFMG
HVDAGKSTMGGNLLYLTGSVDKRTIEKYEREAKDAGRQGWYLSWVMDTNKEERNDGKTIEVG
KAYFETEKRRYTILDAPGHKMYVSEMIGGASQADVGVLVISARKGEYETGFERGGQTREHAL
LAKTQGVNKMVVVVNKMDDPTVNWSKERYDQCVSNVSNFLRAIGYNIKTDVVFMPVSGYSGA
NLKDHVDPKECPWYTGPTLLEYLDTMNHVDRHINAPFMLPIAAKMKDLGTIVEGKIESGHIK
KGQSTLLMPNKTAVEIQNIYNETENEVDMAMCGEQVKLRIKGVEEEDISPGFVLTSPKNPIK
SVTKFVAQIAIVELKSIIAAGFSCVMHVHTAIEEVHIVKLLHKLEKGTNRKSKKPPAFAKKG
MKVIAVLETEAPVCVETYQDYPQLGRFTLRDQGTTIAIGKIVKIAE

PAPA

Score = 0.20, Position = 0

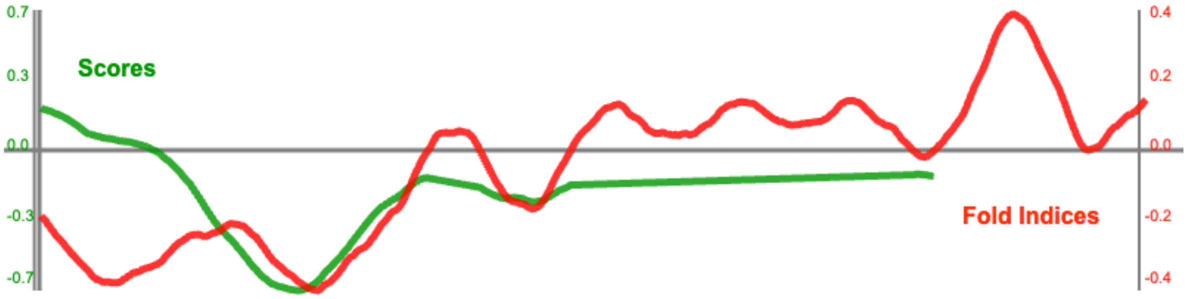


PLAAC

| COREscore | LLR | PAPAprop | PAPAfi | (column descriptions) |
|---|---|---|---|---|
| 19.025 | 19.025 | 0.195 | -0.197 | |

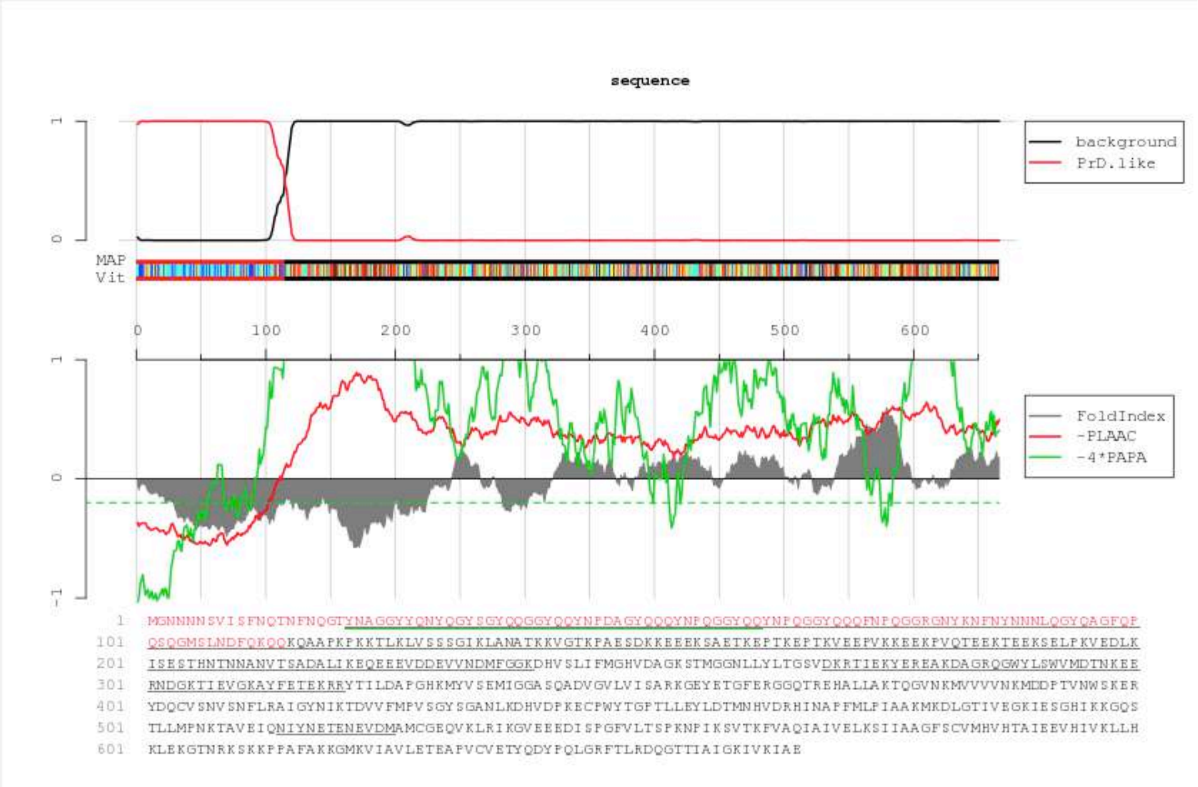

>C2
**MSSVYWLNSVNENNNNKSYYIS**AGGYYQNYQGYSGYQQGGYQQYNPDAGYQQQYNPQGGYQQ
YNPQGGYQQQFNPQGGRGNYKNFNYNNNLQGYQAGFQPQSQGMSLNDFQKQQKQAAPKPKKT
LKLVSSSGIKLANATKKVGTKPAESDKKEEEKSAETKEPTKEPTKVEEPVKKEEKPVQTEEK
TEEKSELPKVEDLKISESTHNTNNANVTSADALIKEQEEEVDDEVVNDMFGGKDHVSLIFMG
HVDAGKSTMGGNLLYLTGSVDKRTIEKYEREAKDAGRQGWYLSWVMDTNKEERNDGKTIEVG
KAYFETEKRRYTILDAPGHKMYVSEMIGGASQADVGVLVISARKGEYETGFERGGQTREHAL
LAKTQGVNKMVVVVNKMDDPTVNWSKERYDQCVSNVSNFLRAIGYNIKTDVVFMPVSGYSGA
NLKDHVDPKECPWYTGPTLLEYLDTMNHVDRHINAPFMLPIAAKMKDLGTIVEGKIESGHIK
KGQSTLLMPNKTAVEIQNIYNETENEVDMAMCGEQVKLRIKGVEEEDISPGFVLTSPKNPIK
SVTKFVAQIAIVELKSIIAAGFSCVMHVHTAIEEVHIVKLLHKLEKGTNRKSKKPPAFAKKG
MKVIAVLETEAPVCVETYQDYPQLGRFTLRDQGTTIAIGKIVKIAE

PAPA

Score = 0.18, Position = 0

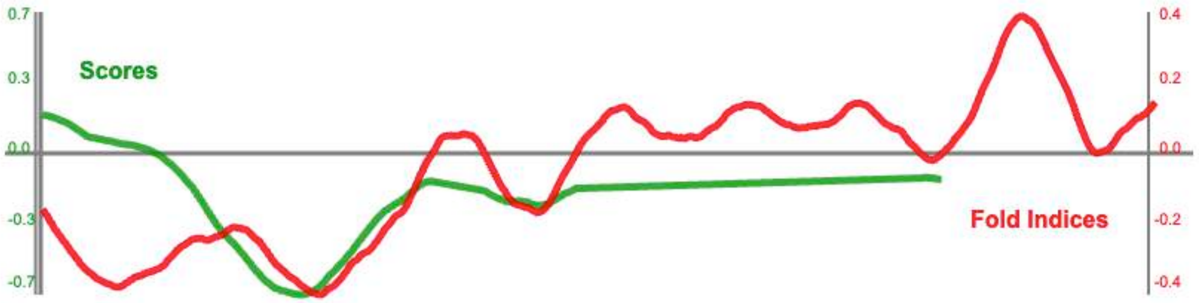


PLAAC

| COREscore | LLR | PAPAprop | PAPAfi | (column descriptions) |
|---|---|---|---|---|
| 19.025 | 19.025 | 0.180 | -0.167 | |

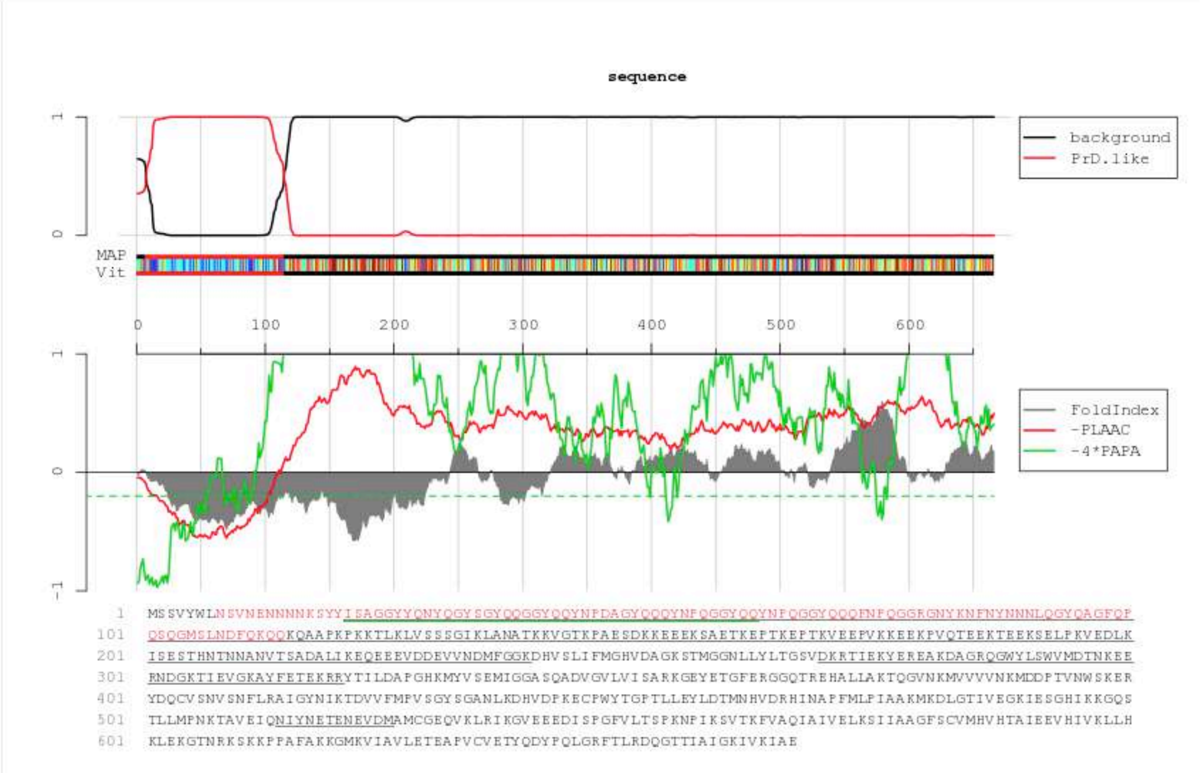

>C3

**M**MQAAFNFINNRRYQDAINVLNAGGYYQNYQGYSGYQQGGYQQYNPDAGYQQQYNPQGGYQQ
YNPQGGYQQQFNPQGGRGNYKNFNYNNNLQGYQAGFQPQSQGMSLNDFQKQQKQAAPKPKKT
LKLVSSSGIKLANATKKVGTKPAESDKKEEEKSAETKEPTKEPTKVEEPVKKEEKPVQTEEK
TEEKSELPKVEDLKISESTHNTNNANVTSADALIKEQEEEVDDEVVNDMFGGKDHVSLIFMG
HVDAGKSTMGGNLLYLTGSVDKRTIEKYEREAKDAGRQGWYLSWVMDTNKEERNDGKTIEVG
KAYFETEKRRYTILDAPGHKMYVSEMIGGASQADVGVLVISARKGEYETGFERGGQTREHAL
LAKTQGVNKMVVVVNKMDDPTVNWSKERYDQCVSNVSNFLRAIGYNIKTDVVFMPVSGYSGA
NLKDHVDPKECPWYTGPTLLEYLDTMNHVDRHINAPFMLPIAAKMKDLGTIVEGKIESGHIK
KGQSTLLMPNKTAVEIQNIYNETENEVDMAMCGEQVKLRIKGVEEEDISPGFVLTSPKNPIK
SVTKFVAQIAIVELKSIIAAGFSCVMHVHTAIEEVHIVKLLHKLEKGTNRKSKKPPAFAKKG
MKVIAVLETEAPVCVETYQDYPQLGRFTLRDQGTTIAIGKIVKIAE

PAPA

Score = 0.15, Position = 0

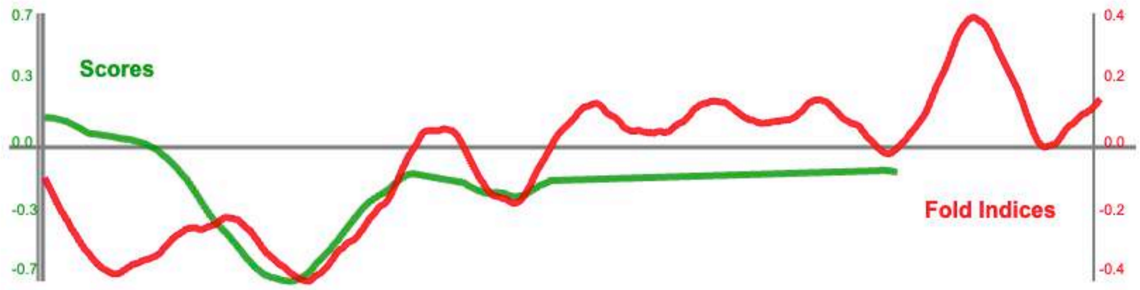


PLAAC

| COREscore | LLR | PAPAprop | PAPAfi | (column descriptions) |
|---|---|---|---|---|
| 19.025 | 19.025 | 0.146 | -0.095 | |

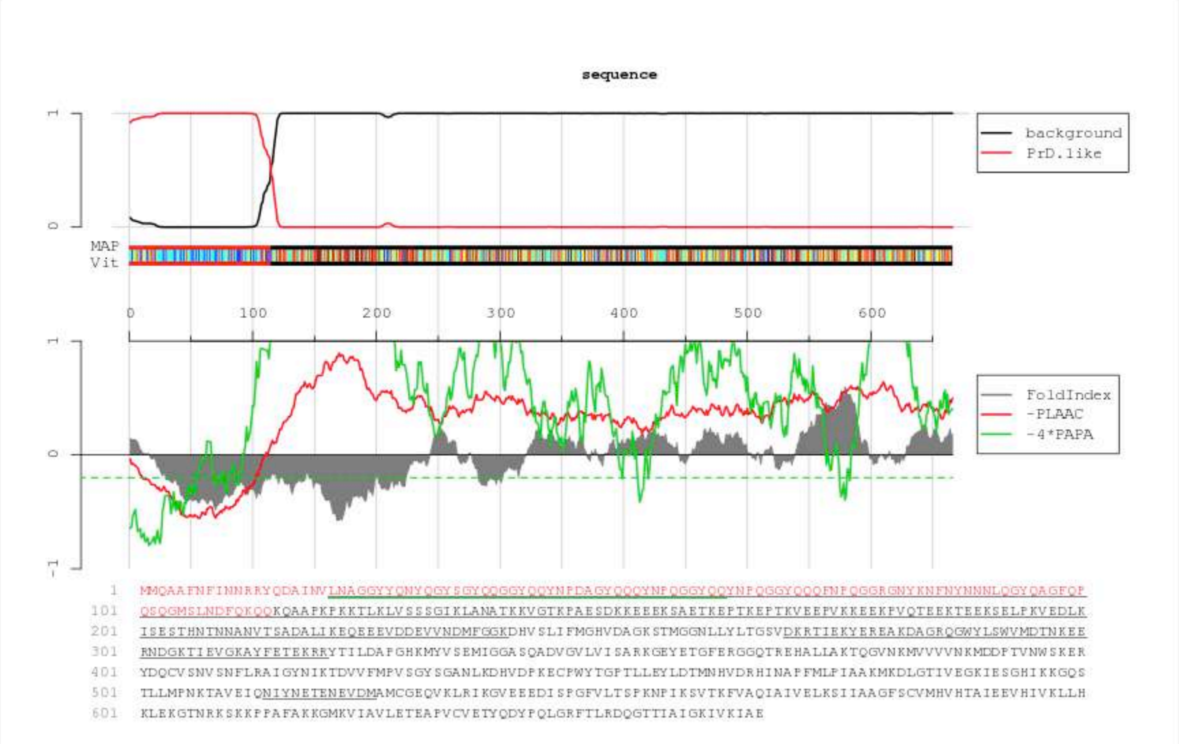

>C4

**MYQQALNVLSRIQNRNAQWYYL**AGGYYQNYQGYSGYQQGGYQQYNPDAGYQQQYNPQGGYQQ
YNPQGGYQQQFNPQGGRGNYKNFNYNNNLQGYQAGFQPQSQGMSLNDFQKQQKQAAPKPKKT
LKLVSSSGIKLANATKKVGTKPAESDKKEEEKSAETKEPTKEPTKVEEPVKKEEKPVQTEEK
TEEKSELPKVEDLKISESTHNTNNANVTSADALIKEQEEEVDDEVVNDMFGGKDHVSLIFMG
HVDAGKSTMGGNLLYLTGSVDKRTIEKYEREAKDAGRQGWYLSWVMDTNKEERNDGKTIEVG
KAYFETEKRRYTILDAPGHKMYVSEMIGGASQADVGVLVISARKGEYETGFERGGQTREHAL
LAKTQGVNKMVVVVNKMDDPTVNWSKERYDQCVSNVSNFLRAIGYNIKTDVVFMPVSGYSGA
NLKDHVDPKECPWYTGPTLLEYLDTMNHVDRHINAPFMLPIAAKMKDLGTIVEGKIESGHIK
KGQSTLLMPNKTAVEIQNIYNETENEVDMAMCGEQVKLRIKGVEEEDISPGFVLTSPKNPIK
SVTKFVAQIAIVELKSIIAAGFSCVMHVHTAIEEVHIVKLLHKLEKGTNRKSKKPPAFAKKG
MKVIAVLETEAPVCVETYQDYPQLGRFTLRDQGTTIAIGKIVKIAE

PAPA

Score = 0.17, Position = 0

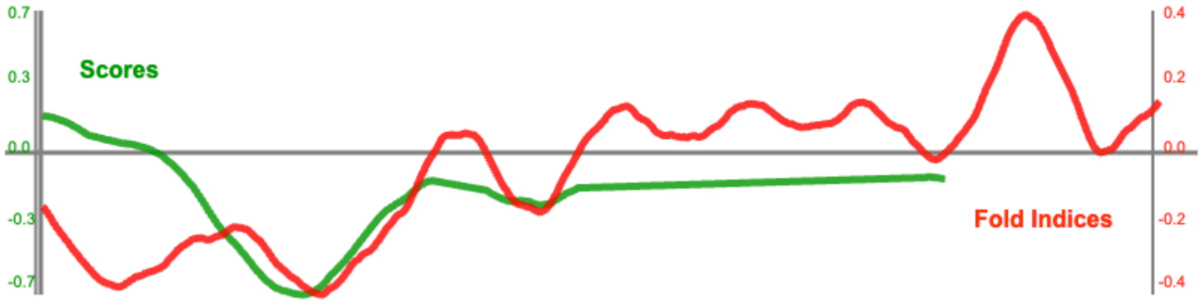


PLAAC

| COREscore | LLR | PAPAprop | PAPAfi | (column descriptions) |
|---|---|---|---|---|
| 19.025 | 19.025 | 0.171 | -0.161 | |

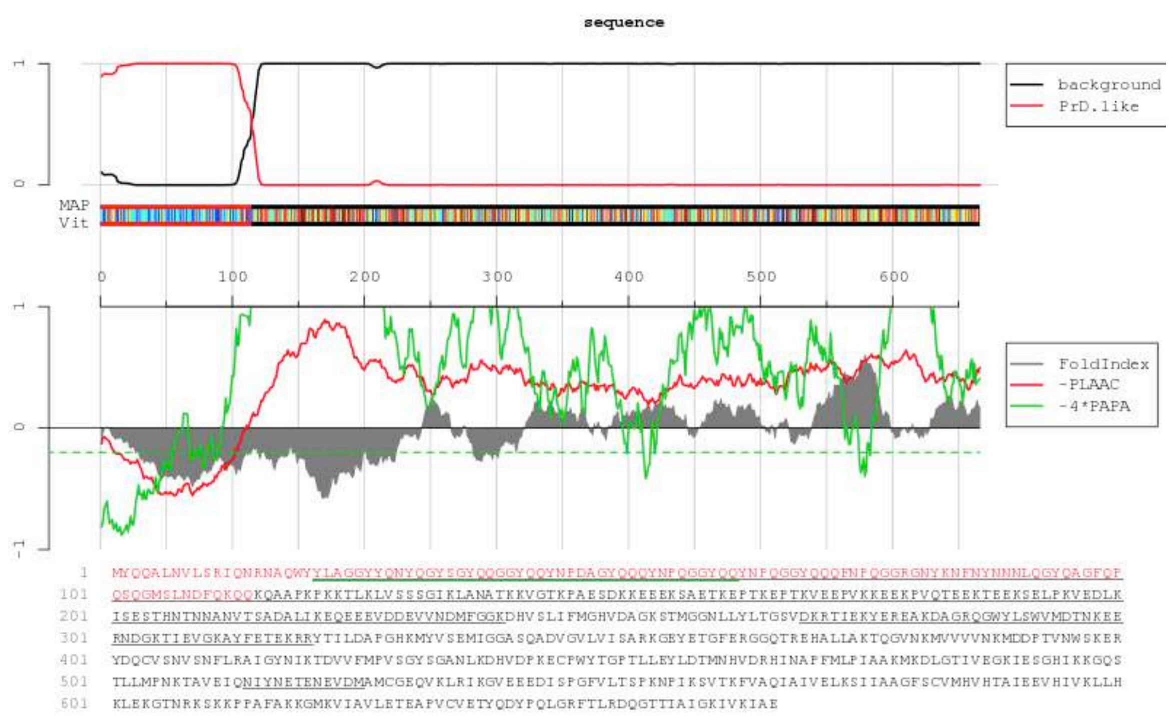

>C5

**MHANEGNVNQYNNSYQNTNSYS**AGGYYQNYQGYSGYQQGGYQQYNPDAGYQQQYNPQGGYQQ
YNPQGGYQQQFNPQGGRGNYKNFNYNNNLQGYQAGFQPQSQGMSLNDFQKQQKQAAPKPKKT
LKLVSSSGIKLANATKKVGTKPAESDKKEEEKSAETKEPTKEPTKVEEPVKKEEKPVQTEEK
TEEKSELPKVEDLKISESTHNTNNANVTSADALIKEQEEEVDDEVVNDMFGGKDHVSLIFMG
HVDAGKSTMGGNLLYLTGSVDKRTIEKYEREAKDAGRQGWYLSWVMDTNKEERNDGKTIEVG
KAYFETEKRRYTILDAPGHKMYVSEMIGGASQADVGVLVISARKGEYETGFERGGQTREHAL
LAKTQGVNKMVVVVNKMDDPTVNWSKERYDQCVSNVSNFLRAIGYNIKTDVVFMPVSGYSGA
NLKDHVDPKECPWYTGPTLLEYLDTMNHVDRHINAPFMLPIAAKMKDLGTIVEGKIESGHIK
KGQSTLLMPNKTAVEIQNIYNETENEVDMAMCGEQVKLRIKGVEEEDISPGFVLTSPKNPIK
SVTKFVAQIAIVELKSIIAAGFSCVMHVHTAIEEVHIVKLLHKLEKGTNRKSKKPPAFAKKG
MKVIAVLETEAPVCVETYQDYPQLGRFTLRDQGTTIAIGKIVKIAE

PAPA

Score = 0.16, Position = 0

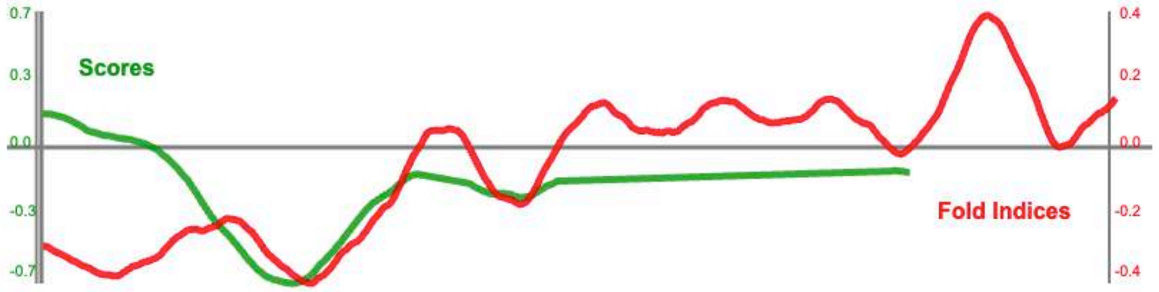


PLAAC

| COREscore | LLR | PAPAprop | PAPAfi | (column descriptions) |
|---|---|---|---|---|
| 19.025 | 19.025 | 0.163 | -0.305 | |

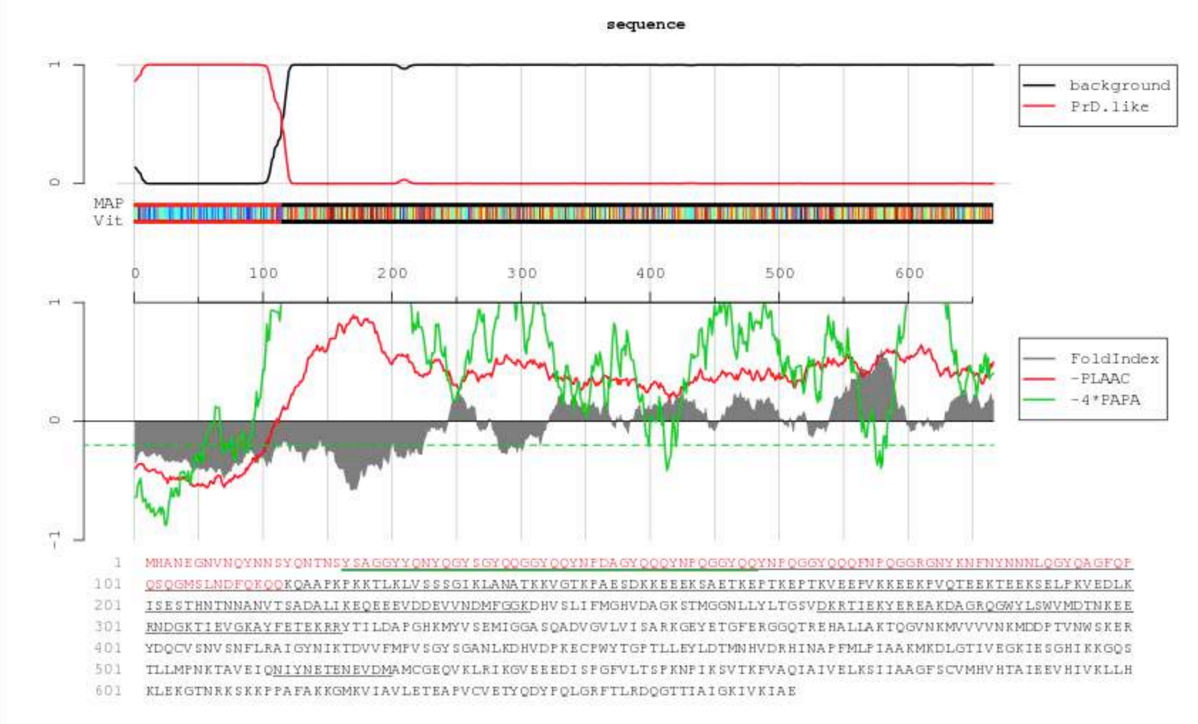

>C6
MMDNNYYNNNNNNNNTNSNSNYNAGGYYQNYQGYSGYQQGGYQQYNPDAGYQQQYNPQGGYQQ
YNPQGGYQQQFNPQGGRGNYKNFNYNNNLQGYQAGFQPQSQGMSLNDFQKQQKQAAPKPKKT
LKLVSSSGIKLANATKKVGTKPAESDKKEEEKSAETKEPTKEPTKVEEPVKKEEKPVQTEEK
TEEKSELPKVEDLKISESTHNTNNANVTSADALIKEQEEEVDDEVVNDMFGGKDHVSLIFMG
HVDAGKSTMGGNLLYLTGSVDKRTIEKYEREAKDAGRQGWYLSWVMDTNKEERNDGKTIEVG
KAYFETEKRRYTILDAPGHKMYVSEMIGGASQADVGVLVISARKGEYETGFERGGQTREHAL
LAKTQGVNKMVVVVNKMDDPTVNWSKERYDQCVSNVSNFLRAIGYNIKTDVVFMPVSGYSGA
NLKDHVDPKECPWYTGPTLLEYLDTMNHVDRHINAPFMLPIAAKMKDLGTIVEGKIESGHIK
KGQSTLLMPNKTAVEIQNIYNETENEVDMAMCGEQVKLRIKGVEEEDISPGFVLTSPKNPIK
SVTKFVAQIAIVELKSIIAAGFSCVMHVHTAIEEVHIVKLLHKLEKGTNRKSKKPPAFAKKG
MKVIAVLETEAPVCVETYQDYPQLGRFTLRDQGTTIAIGKIVKIAE

PAPA

Score = 0.16, Position = 0

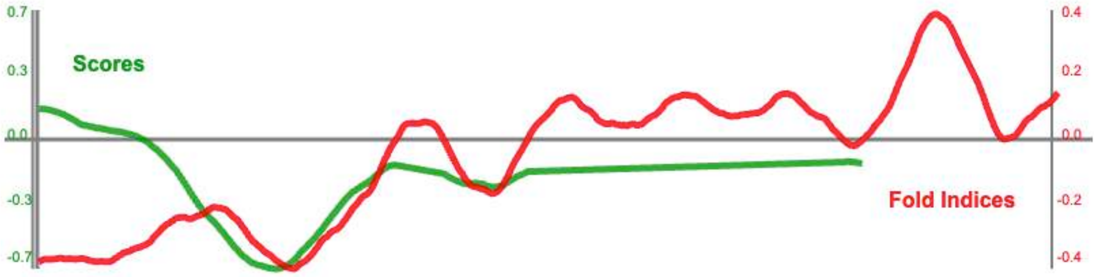


PLAAC

sequence

| COREscore | LLR | PAPAprop | PAPAfi | (column descriptions) |
|---|---|---|---|---|
| 20.492 | 20.492 | 0.157 | -0.398 | |

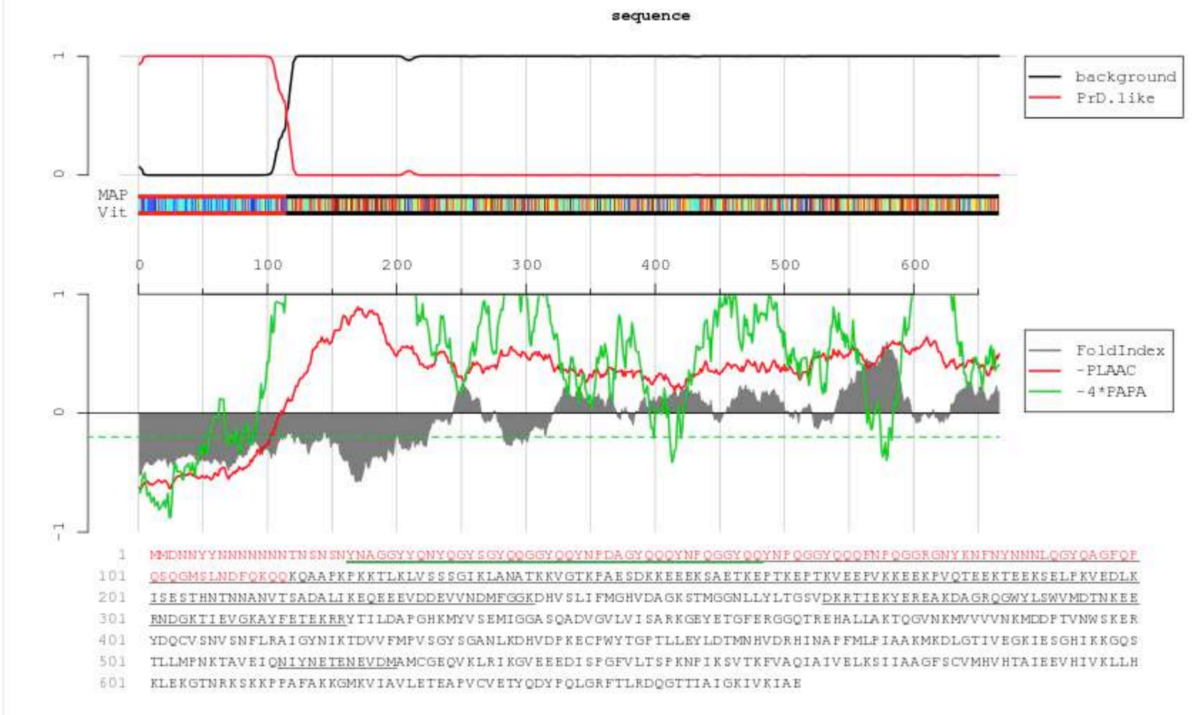

>C7
M**RSGNYVFLANSTNRYQGNYYN**AGGYYQNYQGYSGYQQGGYQQYNPDAGYQQQYNPQGGYQQ
YNPQGGYQQQFNPQGGRGNYKNFNYNNNLQGYQAGFQPQSQGMSLNDFQKQQKQAAPKPKKT
LKLVSSSGIKLANATKKVGTKPAESDKKEEEKSAETKEPTKEPTKVEEPVKKEEKPVQTEEK
TEEKSELPKVEDLKISESTHNTNNANVTSADALIKEQEEEVDDEVVNDMFGGKDHVSLIFMG
HVDAGKSTMGGNLLYLTGSVDKRTIEKYEREAKDAGRQGWYLSWVMDTNKEERNDGKTIEVG
KAYFETEKRRYTILDAPGHKMYVSEMIGGASQADVGVLVISARKGEYETGFERGGQTREHAL
LAKTQGVNKMVVVVNKMDDPTVNWSKERYDQCVSNVSNFLRAIGYNIKTDVVFMPVSGYSGA
NLKDHVDPKECPWYTGPTLLEYLDTMNHVDRHINAPFMLPIAAKMKDLGTIVEGKIESGHIK
KGQSTLLMPNKTAVEIQNIYNETENEVDMAMCGEQVKLRIKGVEEEDISPGFVLTSPKNPIK
SVTKFVAQIAIVELKSIIAAGFSCVMHVHTAIEEVHIVKLLHKLEKGTNRKSKKPPAFAKKG
MKVIAVLETEAPVCVETYQDYPQLGRFTLRDQGTTIAIGKIVKIAE

PAPA

Score = 0.18, Position = 0

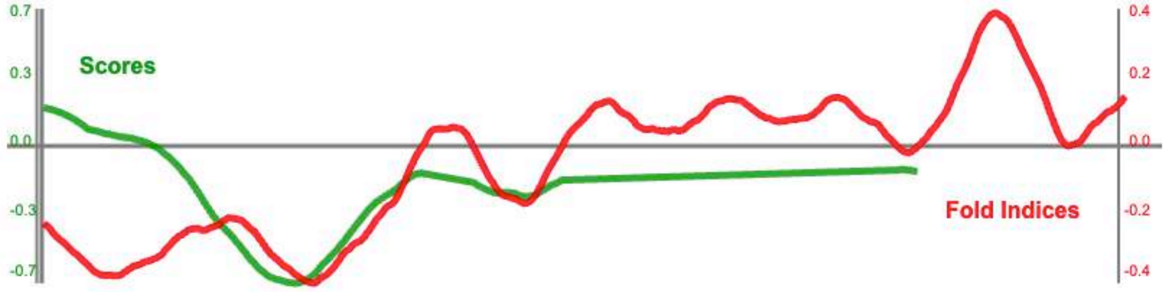


PLAAC

| COREscore | LLR | PAPAprop | PAPAfi | (column descriptions) |
|---|---|---|---|---|
| 19.025 | 19.025 | 0.183 | -0.237 | |

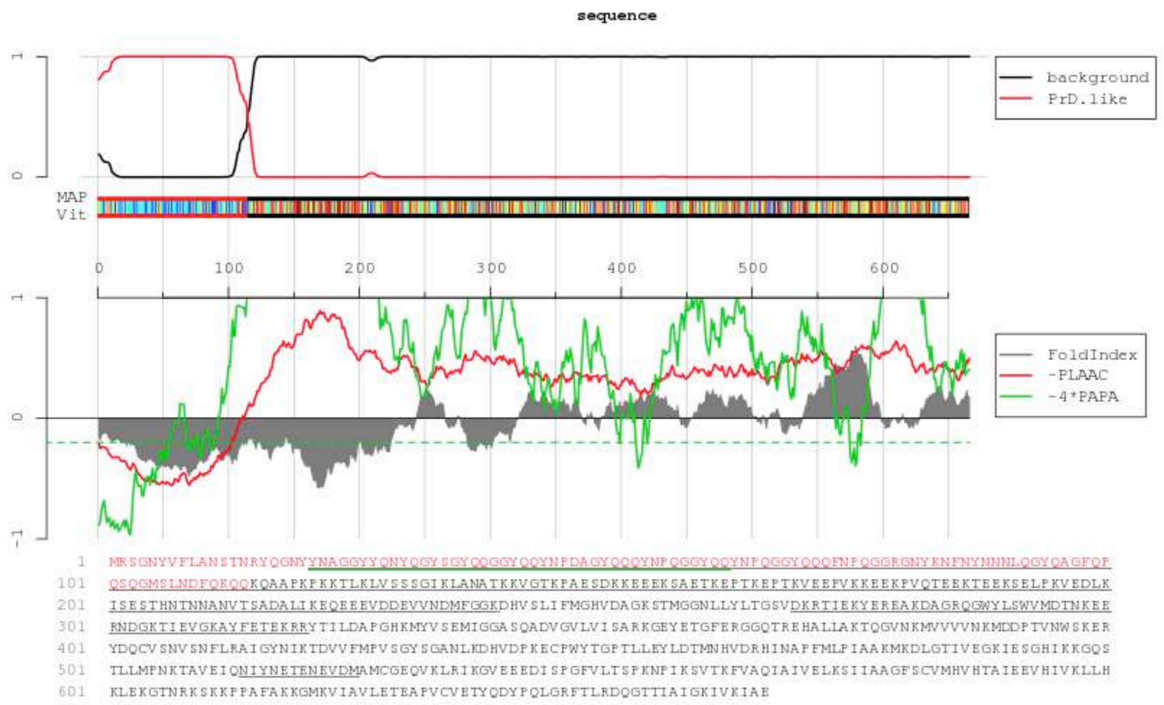

>C8

**MSYSSSANNYGYGSNYNYSYSA**AGGYYQNYQGYSGYQQGGYQQYNPDAGYQQQYNPQGGYQQ
YNPQGGYQQQFNPQGGRGNYKNFNYNNNLQGYQAGFQPQSQGMSLNDFQKQQKQAAPKPKKT
LKLVSSSGIKLANATKKVGTKPAESDKKEEEKSAETKEPTKEPTKVEEPVKKEEKPVQTEEK
TEEKSELPKVEDLKISESTHNTNNANVTSADALIKEQEEEVDDEVVNDMFGGKDHVSLIFMG
HVDAGKSTMGGNLLYLTGSVDKRTIEKYEREAKDAGRQGWYLSWVMDTNKEERNDGKTIEVG
KAYFETEKRRYTILDAPGHKMYVSEMIGGASQADVGVLVISARKGEYETGFERGGQTREHAL
LAKTQGVNKMVVVVNKMDDPTVNWSKERYDQCVSNVSNFLRAIGYNIKTDVVFMPVSGYSGA
NLKDHVDPKECPWYTGPTLLEYLDTMNHVDRHINAPFMLPIAAKMKDLGTIVEGKIESGHIK
KGQSTLLMPNKTAVEIQNIYNETENEVDMAMCGEQVKLRIKGVEEEDISPGFVLTSPKNPIK
SVTKFVAQIAIVELKSIIAAGFSCVMHVHTAIEEVHIVKLLHKLEKGTNRKSKKPPAFAKKG
MKVIAVLETEAPVCVETYQDYPQLGRFTLRDQGTTIAIGKIVKIAE

PAPA

Score = 0.20, Position = 0

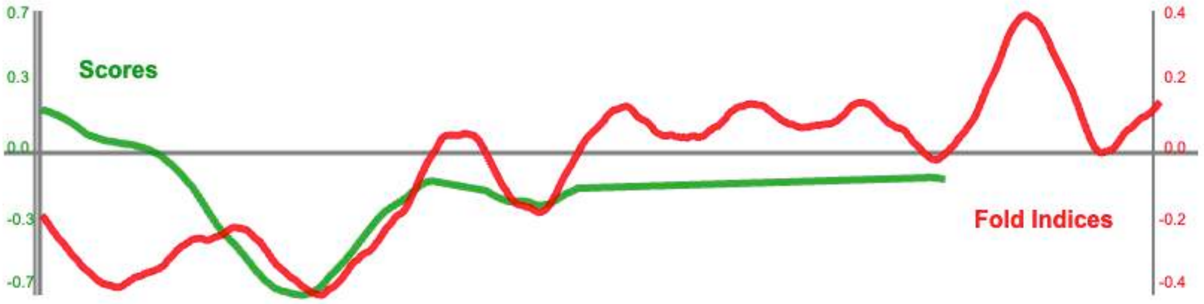


PLAAC

| COREscore | LLR | PAPAprop | PAPAfi | (column descriptions) |
|---|---|---|---|---|
| 19.025 | 19.025 | 0.201 | -0.186 | |

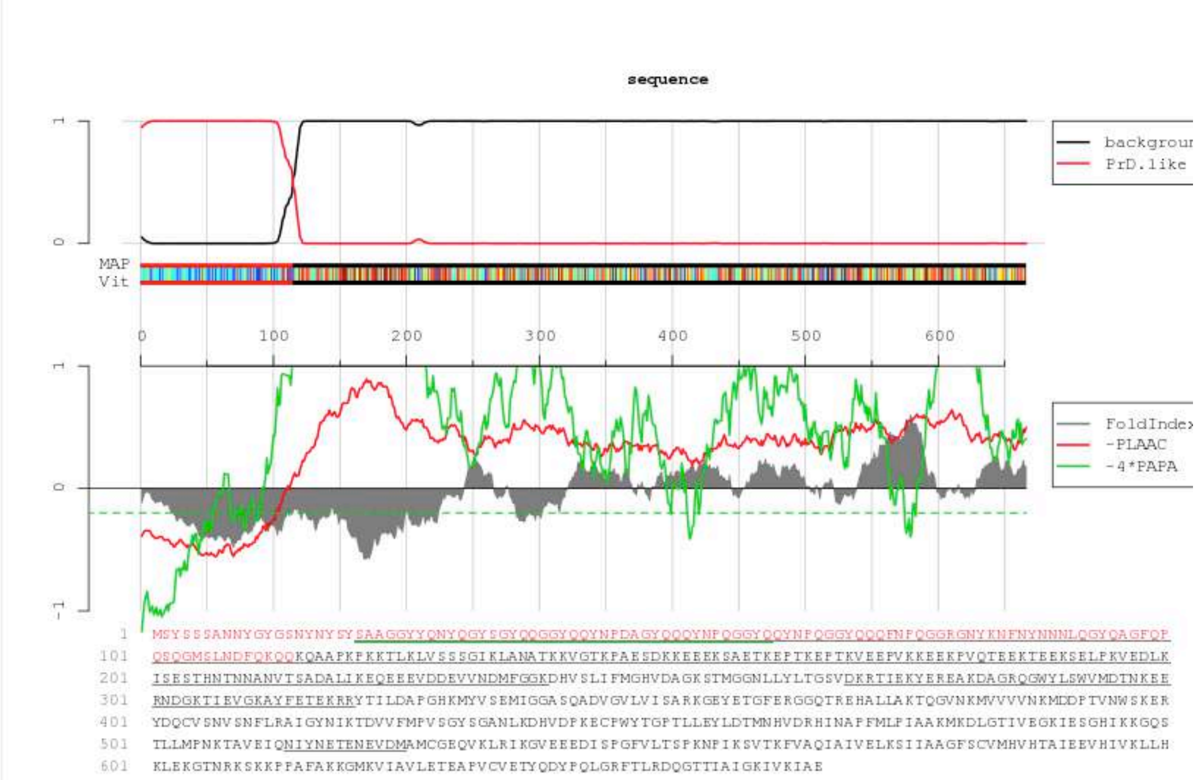

>C9

**MGYVQSSYGQNQFQNSQYGSYQ**AGGYYQNYQGYSGYQQGGYQQYNPDAGYQQQYNPQGGYQQ
YNPQGGYQQQFNPQGGRGNYKNFNYNNNLQGYQAGFQPQSQGMSLNDFQKQQKQAAPKPKKT
LKLVSSSGIKLANATKKVGTKPAESDKKEEEKSAETKEPTKEPTKVEEPVKKEEKPVQTEEK
TEEKSELPKVEDLKISESTHNTNNANVTSADALIKEQEEEVDDEVVNDMFGGKDHVSLIFMG
HVDAGKSTMGGNLLYLTGSVDKRTIEKYEREAKDAGRQGWYLSWVMDTNKEERNDGKTIEVG
KAYFETEKRRYTILDAPGHKMYVSEMIGGASQADVGVLVISARKGEYETGFERGGQTREHAL
LAKTQGVNKMVVVVNKMDDPTVNWSKERYDQCVSNVSNFLRAIGYNIKTDVVFMPVSGYSGA
NLKDHVDPKECPWYTGPTLLEYLDTMNHVDRHINAPFMLPIAAKMKDLGTIVEGKIESGHIK
KGQSTLLMPNKTAVEIQNIYNETENEVDMAMCGEQVKLRIKGVEEEDISPGFVLTSPKNPIK
SVTKFVAQIAIVELKSIIAAGFSCVMHVHTAIEEVHIVKLLHKLEKGTNRKSKKPPAFAKKG
MKVIAVLETEAPVCVETYQDYPQLGRFTLRDQGTTIAIGKIVKIAE

PAPA

Score = 0.21, Position = 0

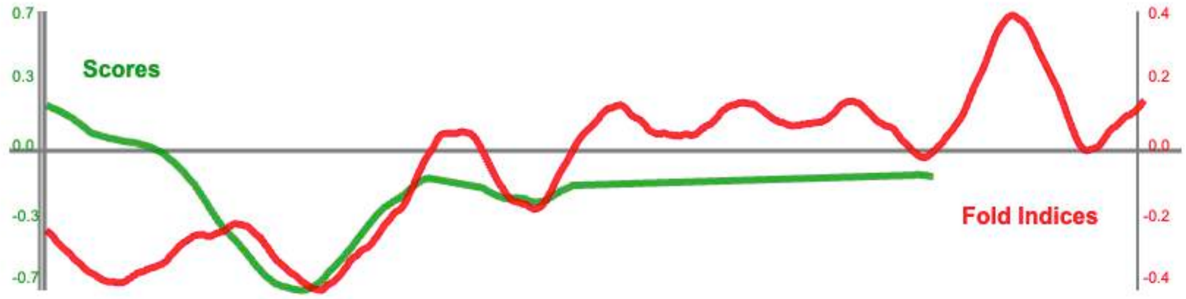


PLAAC

| COREscore | LLR | PAPAprop | PAPAfi | (column descriptions) |
|---|---|---|---|---|
| 19.025 | 19.025 | 0.213 | -0.240 | |

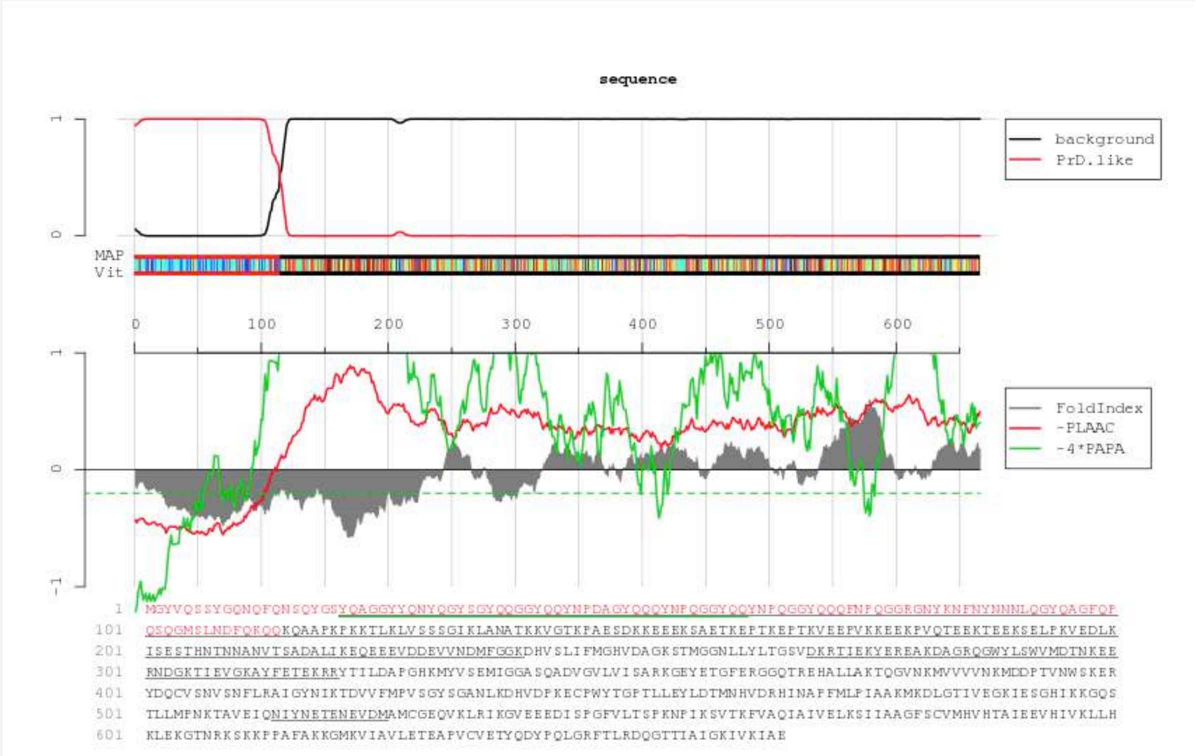

>C10

**MSTGYSDLNSTYNQSALINQIG**AGGYYQNYQGYSGYQQGGYQQYNPDAGYQQQYNPQGGYQQ
YNPQGGYQQQFNPQGGRGNYKNFNYNNNLQGYQAGFQPQSQGMSLNDFQKQQKQAAPKPKKT
LKLVSSSGIKLANATKKVGTKPAESDKKEEEKSAETKEPTKEPTKVEEPVKKEEKPVQTEEK
TEEKSELPKVEDLKISESTHNTNNANVTSADALIKEQEEEVDDEVVNDMFGGKDHVSLIFMG
HVDAGKSTMGGNLLYLTGSVDKRTIEKYEREAKDAGRQGWYLSWVMDTNKEERNDGKTIEVG
KAYFETEKRRYTILDAPGHKMYVSEMIGGASQADVGVLVISARKGEYETGFERGGQTREHAL
LAKTQGVNKMVVVVNKMDDPTVNWSKERYDQCVSNVSNFLRAIGYNIKTDVVFMPVSGYSGA
NLKDHVDPKECPWYTGPTLLEYLDTMNHVDRHINAPFMLPIAAKMKDLGTIVEGKIESGHIK
KGQSTLLMPNKTAVEIQNIYNETENEVDMAMCGEQVKLRIKGVEEEDISPGFVLTSPKNPIK
SVTKFVAQIAIVELKSIIAAGFSCVMHVHTAIEEVHIVKLLHKLEKGTNRKSKKPPAFAKKG
MKVIAVLETEAPVCVETYQDYPQLGRFTLRDQGTTIAIGKIVKIAE

PAPA

Score = 0.14, Position = 0

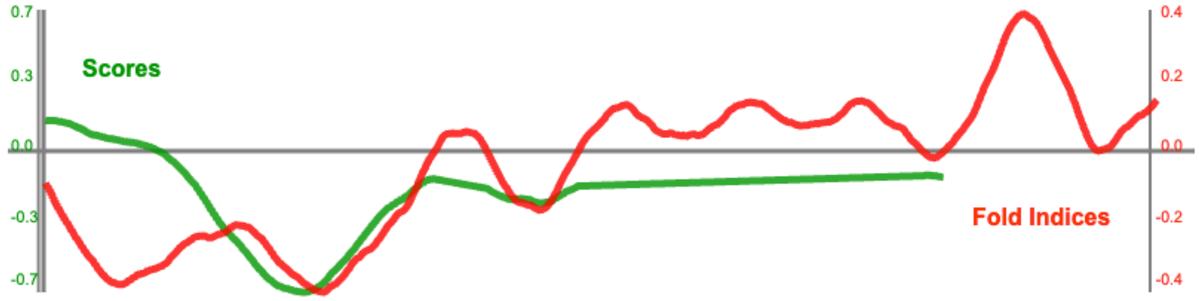


PLAAC

| COREscore | LLR | PAPAprop | PAPAfi | (column descriptions) |
|---|---|---|---|---|
| 19.025 | 19.025 | 0.142 | -0.096 | |

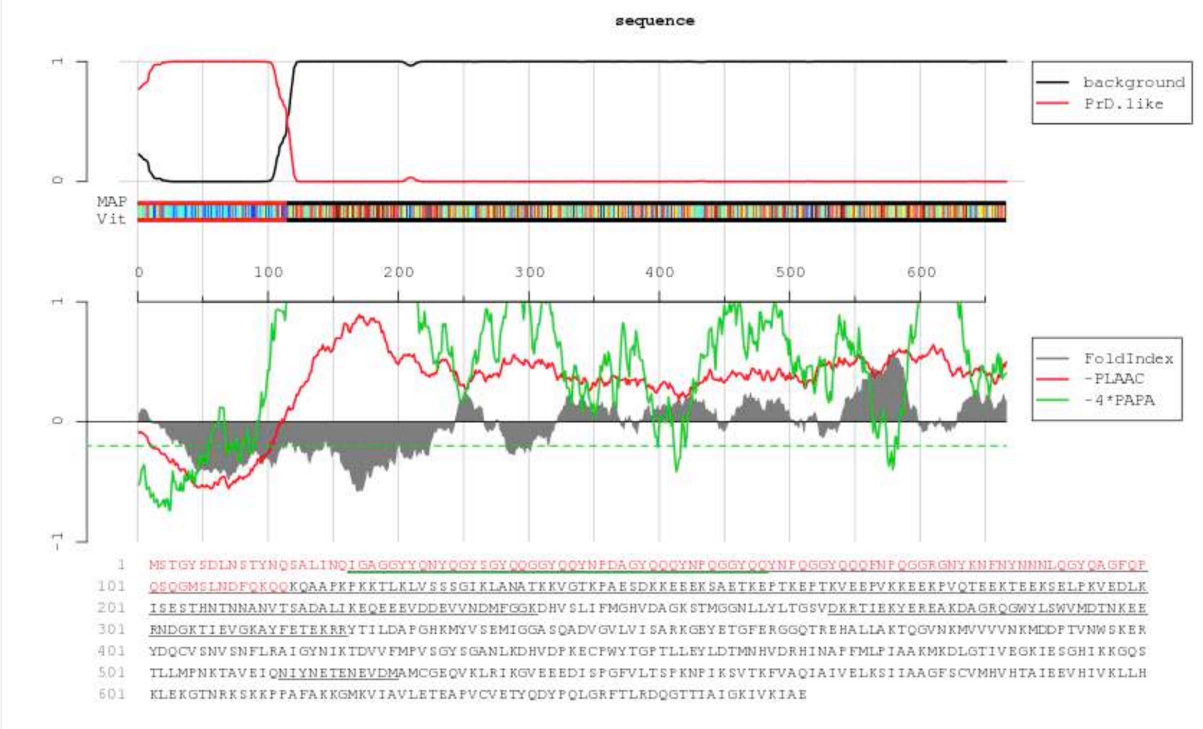

**Supplementary Data 2.** Protein sequences of the Sup35p and the Sup35NM fused to GFP expressed in *S. cerevisiae*.

**Sup35p**

MSDSNQGNNQQNYQQYSQNGNQQQGNNRYQGYQAYNAQAQPAGGYYQNYQGYSGYQQGGYQQ
YNPDAGYQQQYNPQGGYQQYNPQGGYQQQFNPQGGRGNYKNFNYNNNLQGYQAGFQPQSQGM
SLNDFQKQQKQAAPKPKKTLKLVSSSGIKLANATKKVGTKPAESDKKEEEKSAETKEPTKEP
TKVEEPVKKEEKPVQTEEKTEEKSELPKVEDLKISESTHNTNNANVTSADALIKEQEEEVDD
EVVNDMFGGKDHVSLIFMGHVDAGKSTMGGNLLYLTGSVDKRTIEKYEREAKDAGRQGWYLS
WVMDTNKEERNDGKTIEVGKAYFETEKRRYTILDAPGHKMYVSEMIGGASQADVGVLVISAR
KGEYETGFERGGQTREHALLAKTQGVNKMVVVVNKMDDPTVNWSKERYDQCVSNVSNFLRAI
GYNIKTDVVFMPVSGYSGANLKDHVDPKECPWYTGPTLLEYLDTMNHVDRHINAPFMLPIAA
KMKDLGTIVEGKIESGHIKKGQSTLLMPNKTAVEIQNIYNETENEVDMAMCGEQVKLRIKGV
EEEDISPGFVLTSPKNPIKSVTKFVAQIAIVELKSIIAAGFSCVMHVHTAIEEVHIVKLLHK
LEKGTNRKSKKPPAFAKKGMKVIAVLETEAPVCVETYQDYPQLGRFTLRDQGTTIAIGKIVK
IAE

Red: amyloid core substituted by the bacterial peptides; purple: N-domain; brown: M-domain; blue: C-terminal domain (functional as translator terminator eRF3)

**NMGFP fusion**

MSDSNQGNNQQNYQQYSQNGNQQQGNNRYQGYQAYNAQAQPAGGYYQNYQGYSGYQQGGYQQ
YNPDAGYQQQYNPQGGYQQYNPQGGYQQQFNPQGGRGNYKNFNYNNNLQGYQAGFQPQSQGM
SLNDFQKQQKQAAPKPKKTLKLVSSSGIKLANATKKVGTKPAESDKKEEEKSAETKEPTKEP
TKVEEPVKKEEKPVQTEEKTEEKSELPKVEDLKISESTHNTNNANVTSADALIKEQEEEVDD
EVVNDMF
GSAGSAAGSGMSRSKGEELFTGVVPILVELDGDVNGHKFSVSGEGEGDATYGKLTLKFICTT
GKLPVPWPTLVTTLTYGVQCFSRYPDHMKRHDFFKSAMPEGYVQERTISFKDDGNYKTRAEV
KFEGDTLVNRIELKGIDFKEDGNILGHKLEYNYNSHNVYITADKQKNGIKANFKIRHNIEDG
SVQLADHYQQNTPIGDGPVLLPDNHYLSTQSALSKDPNEKRDHMVLLEFVTAAGITHGMDEL
YKDSHHVSMDYKDDDDKI

Red: amyloid core substituted by the bacterial peptides; purple: N-domain; brown: M-domain; cyan: linker; green: GFP

| | GFP phenotype | | | | | | | |
|---|---|---|---|---|---|---|---|---|
| | 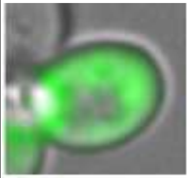 | 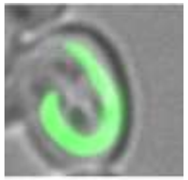 | 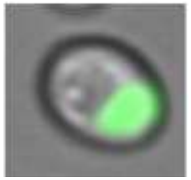 | 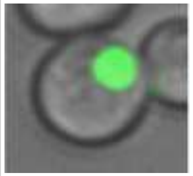 | 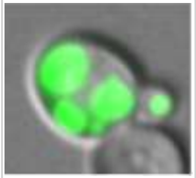 | | 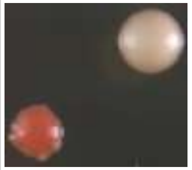 | 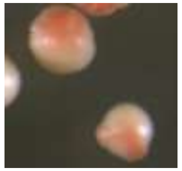 |
| Sequence | Difusse | Ring | Bright focus | Compact focus | Multiple foci | PSI+ | Stable | Unstable |
| NMΔ | 100% | 0,00% | 0% | 0% | 0% | 31% | | +++ |
| NM | 0% | 0,00% | 5,80% | 86,63% | 7,65% | 100% | +++ | |
| C1 | 0% | 0,48% | 11,96% | 55,50% | 32,06% | 100% | +++ | |
| C2 | 0,70% | 0,71% | 18,44% | 70,21% | 9,93% | 98% | +++ | |
| C3 | 0% | 13,86% | 53,46% | 14,85% | 17,82% | 65% | | + |
| C4 | 0% | 0,00% | 74,46% | 25,53% | 0% | 91% | ++ | |
| C5 | 12,90% | 16,12% | 31,45% | 37,90% | 1,61% | 81% | | + |
| C6 | 0% | 11,10% | 27,77% | 59,25% | 1,81% | 66% | | ++ |
| C7 | 0% | 3,26% | 50% | 46,70% | 1,08% | 88% | +++ | |
| C8 | 17,65% | 0,00% | 5,88% | 67,65% | 8,82% | 91% | ++ | |
| C9 | 0% | 6,35% | 17,46% | 75,40% | 0,79% | 67% | ++ | |
| C10 | 38,89% | 20,99% | 6,17% | 33,33% | 0,62% | 75% | | ++ |

**Supplementary Data 3. Table showing the phenotypic characteristics associated to Sup35 expression.** In green, the population distribution (in percentage) of the Sup35NM-GFP location. In red, the percentage of white colonies ([PSI+]). In blue, the proportion of colonies without colored sectors (stable) or with sectors (unstable): + > 1%, ++ > 30%, ++ > 50%.

**Supplementary Data 4.** DNA sequences of the constructs encoding the Sup35p variants (Sup35NM, ΔNM and chimeras) expressed in *E. coli*.

**Green:** First methionine
Pink: CsgAss signal sequence
Black: linker (Not1+serine)
**Blue bold:** The nucleation region
Blue: Sup35NM
**Red:** The amyloid-cores
**Black bold:** 6xHistidines

Sup35NM
**M**KLLKVAAIAAIVFSGSALAGVVPQYGGGGNHGGGGNNSGPNAAAS**DSNQGNNQQNYQQYSQ**
**NGNQQQGNNRYQGYQAYNAQAQP**AGGYYQNYQGYSGYQQGGYQQYNPDAGYQQQYNPQGGYQ
QYNPQGGYQQQFNPQGGRGNYKNFNYNNNLQGYQAGFQPQSQGMSLNDFQKQQKQAAPKPKK
TLKLVSSSGIKLANATKKVGTKPAESDKKEEEKSAETKEPTKEPTKVEEPVKKEEKPVQTEE
KTEEKSELPKVEDLKISESTHNTNNANVTSADALIKEQEEEVDDEVVND**HHHHHH**

ΔSup35NM
**M**KLLKVAAIAAIVFSGSALAGVVPQYGGGGNHGGGGNNSGPNAAASAGGYYQNYQGYSGYQQ
GGYQQYNPDAGYQQQYNPQGGYQQYNPQGGYQQQFNPQGGRGNYKNFNYNNNLQGYQAGFQP
QSQGMSLNDFQKQQKQAAPKPKKTLKLVSSSGIKLANATKKVGTKPAESDKKEEEKSAETKE
PTKEPTKVEEPVKKEEKPVQTEEKTEEKSELPKVEDLKISESTHNTNNANVTSADALIKEQE
EEVDDEVVND**HHHHHH**

Sup35NM chimeras
**M**KLLKVAAIAAIVFSGSALAGVVPQYGGGGNHGGGGNNSGPNAAAS**XXXXXXXXXXXXXXXX**
**XXXXX**AGGYYQNYQGYSGYQQGGYQQYNPDAGYQQQYNPQGGYQQYNPQGGYQQQFNPQGGR
GNYKNFNYNNNLQGYQAGFQPQSQGMSLNDFQKQQKQAAPKPKKTLKLVSSSGIKLANATKK
VGTKPAESDKKEEEKSAETKEPTKEPTKVEEPVKKEEKPVQTEEKTEEKSELPKVEDLKISE
STHNTNNANVTSADALIKEQEEEVDDEVVND**HHHHHH**
